# Small molecule modulators of TOX protein re-invigorate T cell activity

**DOI:** 10.1101/2025.03.03.641115

**Authors:** Bocheng Wu, Heng Jui Chang, Prashant Singh, Alexander Hostetler, Yichen Xiang, Shenghao Guo, Fiona Yihan Wang, Julia J. Zhong, Becky S. Leifer, Richard P. Schiavoni, Nan Jiang, Amit Choudhary, Peter M. K. Westcott, Angela N. Koehler

## Abstract

The TOX protein (thymocyte selection-associated high mobility group box) is a critical transcription factor implicated in both T acute lymphoblastic leukemia (T-ALL) and CD8^+^ T cell exhaustion. Gene perturbation studies suggest that inhibiting TOX may have therapeutic implications for both leukemia and T cell exhaustion. However, due to its complex molecular mechanisms and intrinsically disordered structure, TOX has not been effectively targeted by small molecules to date. In this study, we used small molecule microarray (SMM) screening and biochemical assays to identify a series of TOX protein-protein interaction (PPI) inhibitors. We identified **KI-TOX-A3** as a TOX protein binder and potent TOX PPI inhibitor. In T-ALL, **KI-TOX-A3** revealed selective cytotoxicity and proteosome-dependent TOX degradation. In CD8^+^ T cells, **KI-TOX-A3** potently reversed T cell exhaustion by decreasing surface inhibitory receptors, increasing expression of effector cytokines, and enhancing cancer cell killing activity. We also demonstrate the utility of **KI-TOX-A3** to probe potential epigenetic regulatory mechanisms of TOX via KAT7 acetylation in T cells.

## Introduction

The transcription factor TOX is a pivotal regulator in lymphocytic malignancies, including leukemia and lymphoma. In leukemia, the Myc-driven T-acute lymphoblastic leukemia (T-ALL) implanted in zebrafish showed high *TOX* gene expression, which was positively correlated with MYC expression [1]. TOX was found to promote genome instability and cell growth of T-ALL by binding to the DNA repair heterodimer protein Ku70/80, inhibiting its DNA repair function in the nonhomologous end-joining (NHEJ) pathway[1]. The knockdown of *Tox* decreases the genome instability of T-ALL and induced cytotoxicity to T-ALL cells accordingly. In addition, over-expression of TOX can be found in many other leukemia and lymphoma cells. In the acute myeloid leukemia (AML), high expression of TOX was found to be associated with poor patient overall survival (OS) and may be used as a biomarker to predict clinical outcomes[2]. In lymphoma, TOX is known to be highly expressed in mycosis fungoides (MF) [3], precursor T lymphoblastic lymphoma[4], angioimmunoblastic T-cell lymphoma (AITL)[4] and cutaneous T-cell lymphoma (CTCL)[5]. In CTCL, TOX negatively regulates the tumor suppressor Runx3[5]. In B-cell lymphoma, TOX is overexpressed in precursor B lymphoblastic lymphoma (B-LBL), diffuse large B cell lymphoma (DLBCL), follicular lymphoma (FL), and a small proportion of Burkitt lymphoma (BL) patients[4]. Overall, TOX seems to play a critical role in blood cancers and is a potential therapeutic target.

In T cells, basal expression of TOX is crucial for thymic T cell development [6]. TOX is required for the CD4^+^ lineage transcriptional program. At the molecular level, zinc finger and BTB domain-containing protein 7B (thPOK) upregulation and CD8 lineage repression were compromised in the absence of TOX [7]. In cancer, TOX is required in CD8^+^ T cells to differentiate into the tumor-specific T cells and to perform tumor infiltration [8].

However, in the context of chronic stimulation, such as cancer or chronic viral infection, high levels of TOX drive CD8+ T cell exhaustion [8-11]. T cell exhaustion is a hyporesponsive state characterized by reduced effector cytokines (e.g., TNF-α, IFN-γ) and increased co-inhibitory receptors (co-IRs, e.g., PD-1, LAG-3, TIM3) [12-14]. Exhausted T cells (T_ex_) have lower rates of proliferation and reduced cytotoxicity against cognate antigen-expressing cells [12,15]. T cell exhaustion was defined to have three stages: progenitor exhausted T cells (T_ex_^prog^), intermediate exhausted T cells (T_ex_^int^), and terminal exhausted T cells (T_ex_^term^), while TOX starts to be overexpressed in the stage of T_ex_^int^ [16].

The molecular mechanisms of TOX in T cell exhaustion process remain poorly understood. TOX was found to manipulate chromatin accessibility and regulate gene expressions epigenetically, including upregulation of multiple IRs, such as PD-1, LAG-3, TIM-3, and downregulation of effector cytokines, such as TNF-α and IFN-γ [9,17]. However, the direct DNA binding activity of TOX is unclear. One study identified a 10-mer binding motif of TOX that is rich in GC sequences [18], while other studies suggest that TOX lacks key hydrophobic amino acids to interact with DNA and therefore is more likely to regulate transcription via protein-protein interactions at its transactivation domain[19]. Indeed, multiple protein-protein interactions (PPIs) with TOX have been identified, including with the HBO1 complex via interaction with KAT7—a histone acetyltransferase that regulates H3K14 and H4K12 acetylation and many other epigenetic regulation enzymes such as Dnmt1[9]. However, the role of TOX in this interaction is not fully understood.

Structurally, TOX is a difficult drug target as it contains conformationally dynamic regions and is often referred to as an intrinsically disordered protein (IDP). While conventional drug discovery efforts have successfully targeted some transcription factors (TFs), a significant subset remain notoriously elusive, deemed “undruggable” due to either disordered regions or involvement in essential cellular functions [20]. Currently, there is no crystal or Cryo-EM structure of TOX, due to its conformational plasticity. The nuclear magnetic resonance (NMR) structure of TOX in the Protein Data Bank (PDB) named 2CO9 only contains the structured HMG-box DNA binding domain (DBD). Based on the AlphaFold predicted full-length structure (AF-A0A077ZAP3), no obvious drug binding hydrophobic pockets are apparent[21-23]. However, targeting such IDPs is not impossible. IDPs are highly flexible and many IDPs can interact with proteins and small molecules via conformational changes [24,25].

Overall, TOX regulation of oncogenesis and T cell exhaustion nominate it as a highly promising therapeutic target for both blood cancer treatment and immunotherapy. A small molecule modulator of TOX is highly desirable, both for potential therapeutic development and as a tool for deeper mechanistic studies of TOX biology. Here, we present the discovery of a potent TOX PPI inhibitor through a systematic chemical probe screening approach. We found that the small molecule induces apoptosis of leukemia cells and rescues T cells from exhaustion, consistent with potent inhibitory action against TOX protein.

## Results

### Discovery of TOX regulatory small molecules

To facilitate discovery of chemical modulators of a challenging target such as TOX, we used the small molecule microarray (SMM) screening method for discovering protein-small molecule interactions. This technique has been applied to discover promising chemicals against many target proteins, such as IL-4[26], CDK9 [27], Max[28], ETV1[29], and ORAI1[30]. Following the conventional method [31], we employed small-molecule microarrays (SMMs) to screen pure full-length protein TOX (expressed by HEK293T) against a library of 65,000 immobilized compounds (Figure 1B). The library includes chemical fragments, drug-like molecules, precompetitive drug-like structures, and an overrepresentation of rapamycin as an FKBP12 binder unrelated to TOX protein to serve as a null distribution. After excluding non-specific binders to other proteins screened against the same chemical library, 280 hits were identified based on a cut-off Z-score of 3 (Fig. 1C). The Z-score indicates the significance of fluorescent signal difference from TOX antibody and the background noise, which can be interpreted as chemical binding score.

**Figure 1.**
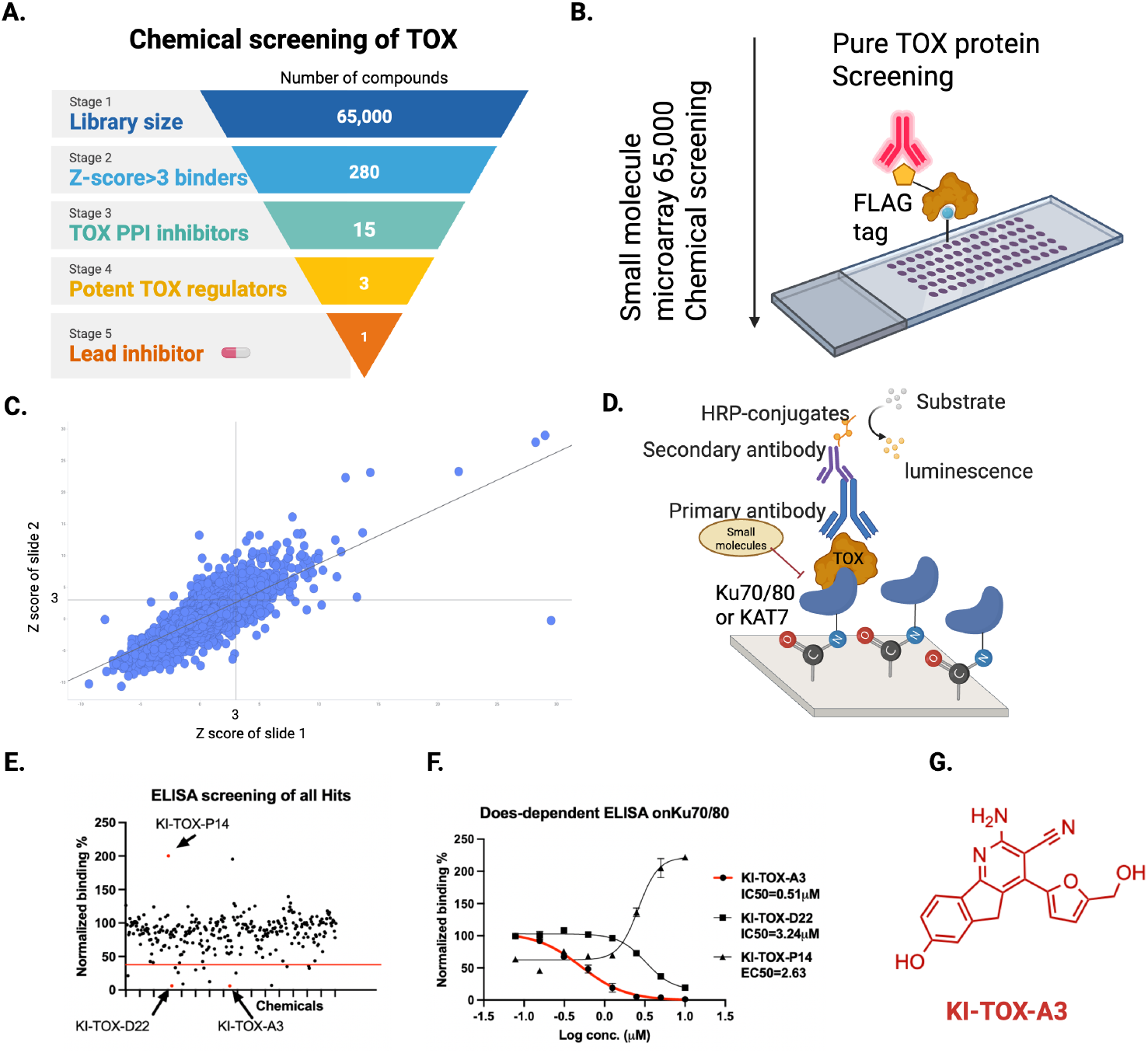
Screening of TOX binders and inhibitors. (A) Screening workflow. (B). The methodology and design of small molecule microarray using pure mammalian TOX protein. (C). The SMM hit chart with a cut-off gate at Z score=3. (D) Design of a developed protein-protein interaction assay as the secondary screening method. (E) ELISA screening of all 280 hits with set gate of >70% inhibition. (F) Three candidates that showed repeated dose dependent TOX PPI regulations. (G) Structure of KI-TOX-A3.

To further filter hits to those capable of regulating TOX through direct interaction, we designed an ELISA assay to detect PPI formation and inhibition. PPIs are critical to the function of many transcription factors, that can induce a transcription complex and promote cellular transcription[20]. As aforementioned, TOX has PPIs with KU70/80 [1] and KAT7 [9]. The assay was designed to bind TOX to the coated partner protein KU70/80 on an NHS-active white plate, as illustrated in Fig. 1D. Using a single-dose concentration (10 µM) screen, 15 SMM assay positives exhibited more than 70% inhibition of the TOX binding with KU70/80 (Fig. 1E). Subsequently, a dose-dependent binding assay was performed using these 15 candidates, ranging from 10 µM to 0.156 µM further confirms the final hits candidates. Three compounds showed significant impact on the PPI of TOX (Figs. 1F and S1B). The candidate **KI-TOX-A3** demonstrated a promising dose-dependent inhibition of the TOX-KAT7 interaction, with an IC_50_ of 0.51 µM. Another candidate, **KI-TOX-D22**, showed a consistent IC_50_ of 3.2 µM on the TOX-KAT7 interaction. Interestingly, **KI-TOX-P14** exhibited mild inhibition (approximately 40%) at concentrations from 0.125 µM to 1.25 µM, followed by a signal increase up to 250% of the positive control, suggesting a potential inverse effect at higher concentrations (Fig. 1F). In the supplementary information, we provide additional details on other inhibitors and their inhibition curves. While several compounds showed promising inhibition activity, we selected **KI-TOX-A3** (Fig.1G) as the primary candidate due to its potent and dose-dependent inhibition performance.

### Target validation of KI-TOX-A3 inhibition of TOX

As TOX binding to KAT7 has been previously demonstrated [9] and confirmed by our binding assay, we performed a TOX-KAT7 proximity ligation assay in Molt-4 cells to visualize the intracellular inhibition evidence (Fig. 2a). We chose Molt-4 because this cell overexpresses TOX protein according to the Human Protein Atlas[32]. Briefly, we treated Molt-4 cells with **KI-TOX-A3, KI-TOX-D22**, and **KI-TOX-P14** at 0, 5 and 10 µM in 1% (v/v) DMSO for 24h. We quantified TOX-KAT7 interaction by confocal microscopy, revealing that **KI-TOX-A3** and **KI-TOX-D22** significantly reduced TOX-KAT7 interaction at 5 and 10 µM. As expected, **KI-TOX-P14** slightly enhanced TOX-KAT7 interaction at higher concentration (10 µM) (Fig. 2A).

**Figure 2.**
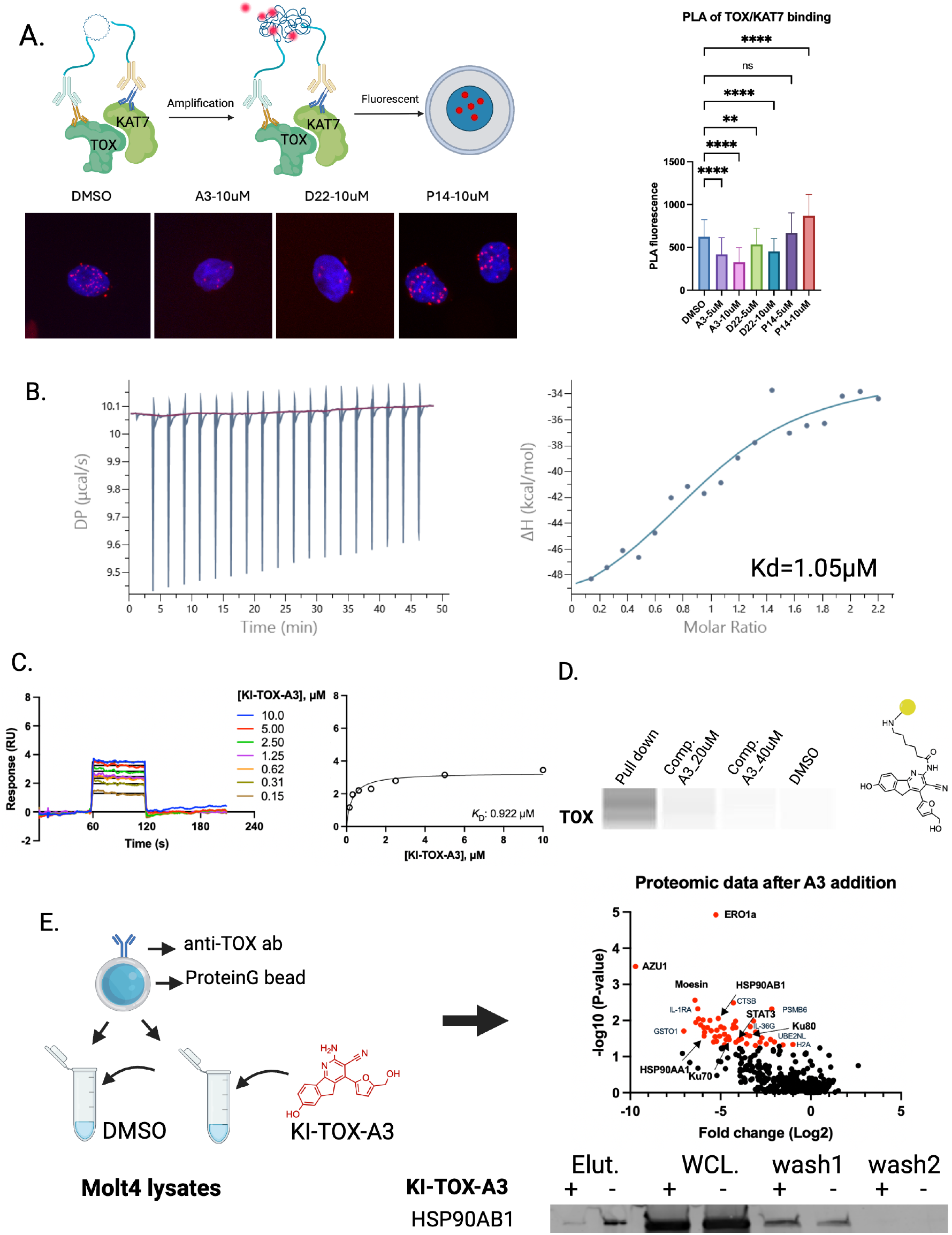
Candidate probes bind to TOX protein. (A). The proximity ligation assay (PLA) revealed candidate modification towards the protein-protein interaction of TOX and KAT7 in Molt-4 cells. (B). Isothermal Titration Calorimetry (ITC) was used to estimate the binding affinity of **KI-TOX-A3** towards TOX protein and concluded with a *K*_*D*_=1.05µM. (C) SPR binding studies demonstrated **KI-TOX-A3** binding to TOX protein with a *K*_*D*_=0.92µM. (D). Molt-4 pulldown assay showed that the TOX bound towards **KI-TOX-A3-0239** on the NHS-bead and lost its binding with KI-TOX-A3 competition at 20 and 40µM. (E). An anti-Tox IP assay shows that the **KI-TOX-A3** blocks binding partners of TOX in Molt-4 lysates. Bars show mean plus standard deviation; * P < 0.05; ** P < 0.0021; ***P < 0.0002; ****P<0.0001).

To confirm that the binding activity of the **KI-TOX-A3** to the TOX protein, we expressed TOX protein using the C43 clone of E.Coli as the host. The His-SUMO-TOX was stably made, and the protocol has been fully provided in the SI. We successfully estimated the KI-TOX-A3 binding to TOX protein using Isothermal Titration Calorimetry (ITC). Briefly, we titrated 100μM KI-TOX-A3 into a pool of concentrated TOX protein and obtained KD=1.05μM, which is close to the PPI inhibition data we collected in Fig. 1F and 2A. To provide further evidence of TOX binding, we also performed a surface plasmon resonance (SPR) binding assay involving full-length recombinant biotinylated human TOX protein using a high-affinity streptavidin (SA) sensor chip. The protein plasmid design and protocols of expression, purification and biotinylation are listed in SI of methods. The sensor gram curve shown in figure 2C shows that **KI-TOX-A3** binds to TOX with a *K*_*D*_=0.92µM, close enough to the ITC estimation (Fig. 2B) and the IC_50_ of the *in vitro* binding assay (Fig. 1F and 2A). In addition, **KI-TOX-P14** binds to TOX, saturating in the 5 µM and the *K*_*D*_=0.79 µM (Fig. S1D).

Next, we performed competition studies to further evaluate the specificity of **KI-TOX-A3**. We performed a lysate immunoprecipitation (IP) assay with the **KI-TOX-A3** derivative (**KI-TOX-A3-0239**, shown in fig. 2D and S2D) incubated with different concentrations of **KI-TOX-A3**. The methylene linker was utilized to provide distance from the bead to enable protein binding, and the primary amine is crucial to form a covalent bond with the bead. The detailed pulldown protocol is provided in the methods section. Soluble competition with **KI-TOX-A3** at 20 µM and 40 µM decreased the intensity of the TOX band, indicating that the **KI-TOX-A3** competes out TOX protein binding in the pulldown process, suggestive of selective TOX binding with **KI-TOX-A3**.

To explore the interactomes of TOX in Molt-4 cell line, we used pulldown assays involving the TOX protein in the whole cell lysate of Molt-4 cells with treatment of DMSO or **KI-TOX-A3** After pulldown, the eluted TOX interactomes were analyzed by LC-MS (Fig. 2E). Indeed, **KI-TOX-A3** reproducibly inhibited protein interactions of TOX (Fig. 2E). The volcano plot shows hits with p-value>0.05 and fold change >1 or <-1 are all negatively regulated (red dots), indicating that **KI-TOX-A3** cause a loss of PPI of TOX. We observed that HSP90AA1 and HSP90AB1 were pulled down in the DMSO control group but not the **KI-TOX-A3** group, indicating that **KI-TOX-A3** may inhibit TOX-HSP binding. Chaperone cycles including HSP90s are critical for correct protein folding, especially overexpressed proteins in the context of cancer [33-35]. This implies that **KI-TOX-A3** could potentially destabilize the TOX protein in T-ALL cells. In addition to HSPs, **KI-TOX-A3** perturbed other potential TOX PPIs in T-ALL cells (Table S1). This implies that the **KI-TOX-A3** modulates multiple interactions within the TOX interactome. Future studies will involve validating the bulk of these putative PPIs.

### KI-TOX-A3 demonstrates potent intracellular activity against cancer cells

Next, we sought to understand the intracellular effects of **KI-TOX-A3** in cancer cells known to be dependent on TOX. We first probed the cytotoxicity of **KI-TOX-A3** against the T-ALL (Jurkat, Molt-4, HBP-ALL), CTCL (HH, Hut78), and B-cell lymphoma (K562, Raji) cell lines (Fig. 3A and 3B). **KI-TOX-A3** was selectively cytotoxic to all TOX-dependent cell lines (T-ALL and CTCL cell lines) and relatively less toxic to TOX-independent cells with limited levels of TOX protein [4]. These data indicate on-target selective inhibition of TOX intracellularly (Fig. 3A and B).

**Figure 3.**
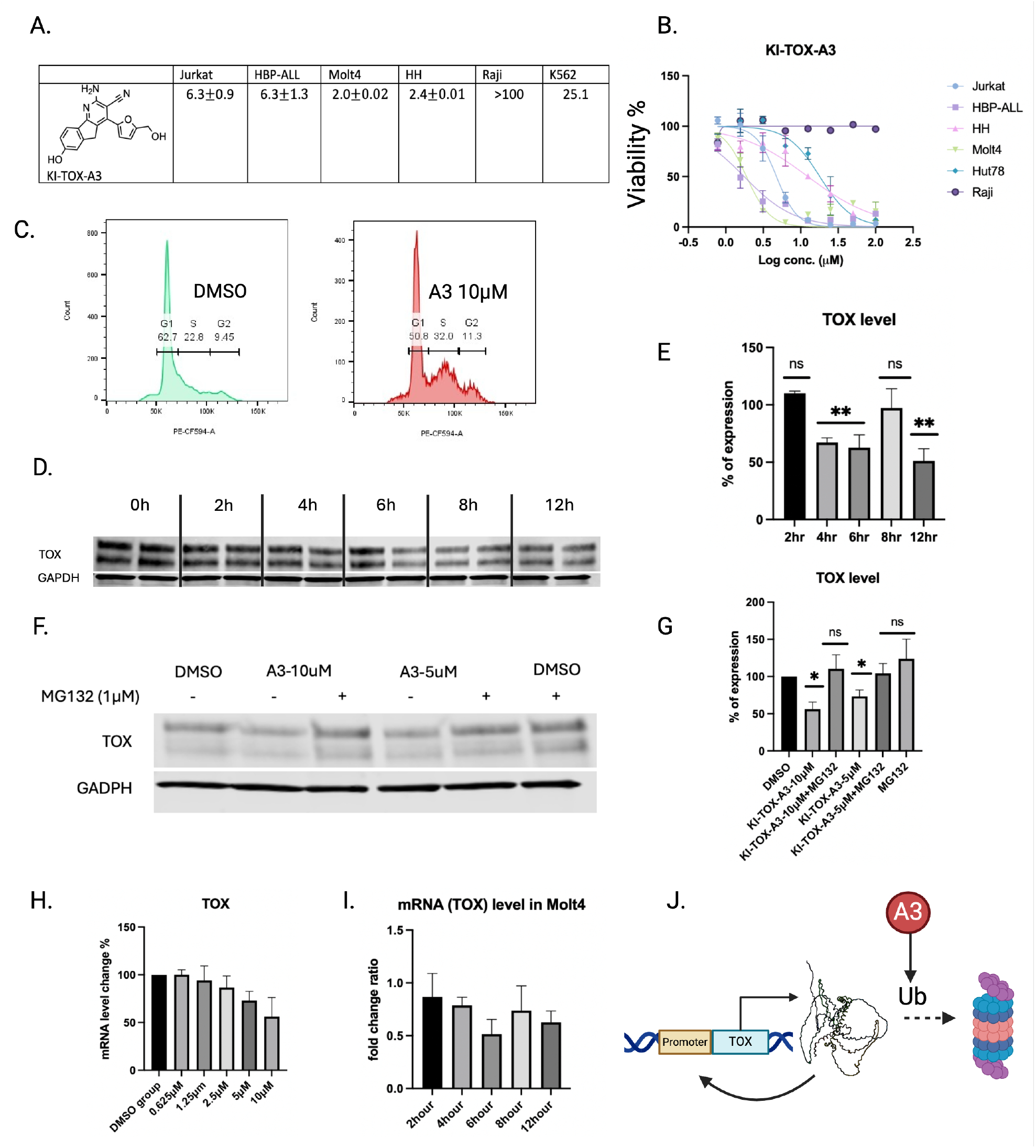
Intracellular effect of candidate TOX chemical probes. (A). The cytotoxicity of **KI-TOX-A3, KI-TOX-D22**, and **KI-TOX-P14** against cancer cells after 96h treatment. (B). The **KI-TOX-A3** inhibits T-ALL cell proliferation but not Raji (a B-cell lymphoma). (C). The cell cycle arrest effect from **KI-TOX-A3** against Jurkat cells. (D and F). **KI-TOX-A3** can downregulate TOX protein after 6h treatment. (E and G) TOX is rescued by the proteasome inhibitor MG132 (1 µM). (H) The **KI-TOX-A3** can downregulate TOX protein and mRNA level dose-dependently and (I) timely in Jurkat cells, and (J) A hypothesized mechanism of action of TOXi **KI-TOX-A3** against TOX protein.

It has previously been shown that shRNA knockdown of *Tox* triggered S phase arrest and impaired proliferation of T-ALL cells including Jurkat cells [1]. Accordingly, we assessed the cell cycle effects of **KI-TOX-A3** in Jurkat cells. Excitingly, we noticed that 10 µM **KI-TOX-A3** treatment could also trigger S phase cell cycle arrest that matches the previous observation (Fig. 3C). We also investigated TOX protein expression in Molt-4 cells by Western blot. Interestingly, we found downregulation of TOX protein after a 4h treatment (Fig. 3D and E), indicating a potential effect of 10 µM **KI-TOX-A3** on proteosome-mediated degradation of TOX. To test this, we performd a 12-h treatment of 5 and 10 µM **KI-TOX-A3** in Molt-4 cells and found that the 1µM MG-132, a proteosome inhibitor [36], rescued TOX protein levels, which implies that the **KI-TOX-A3** might downregulate TOX protein via degradation (Fig. 3F and G). Many transcription factors utilize positive autoregulation, a process where changes in protein levels directly impact transcription levels of the corresponding gene [37]. To clarify whether the TOX protein level change would impact the mRNA level of TOX, we performed qPCR to monitor TOX mRNA levels. We observed the mRNA level of TOX decreased in a dose-dependent manner after 12h-treatment (Fig. 3H). We also tested time-course changes and found that TOX mRNA start to decrease significantly after 4h treatment, but not at earlier time points. This is slightly later than protein level downregulation, which is a further suggestive of autoregulation (Fig. 3I). Overall, we hypothesize that the TOX inhibitor **KI-TOX-A3** binds to TOX which further leads to PPI inhibition, especially its interactions with chaperone partners. This ligand-protein interaction impacts TOX protein levels, which may impact the TOX gene in transcription level with TF positive autoregulation (Fig. 3J).

### KI-TOX-A3 Binding site investigation

We sought to understand the molecular basis of **KI-TOX-A3** binding to TOX. Based on the competition IP of TOX using the **KI-TOX-A3** derivative shown in figure 2, we believe that the amino acid is not critical for TOX binding and permissive to substitution.

Before initiating SAR studies, we briefly investigated the potential binding site(s) on TOX. .To explore potential binding in the DNA-binding domain (DBD) region, we conducted an AlphaFold simulation of TOX-KAT7 binding, where the DBD region is postulated to play a role in PPI formation. Previous studies have shown that TOX protein lacking the DBD region has reduced binding to KU70/80[1]. We used an electrophoretic mobility shift assay (EMSA) involving a DNA probe to investigate TOX binding capacity and TOX inhibitor effect. The DNA strand that we used is not the consensus sequence of TOX protein. Instead, this is a DNA strand (GGCGGGGGCG) that TOX can bind using DNA adenine methyltransferase identification (DamID) sequencing method [18]. In TOX titration data, we added TOX with increasing concentration from 0μM to 5μM with 50nM DNA probe. From the EMSA data, we noticed that the DNA probe was successfully bound by TOX as judged by the intensity of the bands. The TOX-DNA binding was decreased dose-dependently with lower concentration of TOX (Fig.4A, left). Later, we added different doses of small molecule **KI-TOX-A3** to 0.5μM TOX and 50nM DNA mixture. Interestingly, **KI-TOX-A3** did not inhibit TOX-DNA binding from 0.04 to 20µM (Fig. 4A, right), whereas compound treatment in this dose range did modulate the PPI with KAT7. This implies that **KI-TOX-A3** does not interrupt the highly conserved HMG-box helix sequence DBD region binding towards DNA. This also suggests that our candidate TOX inhibitors do not impact the other HMG-box contained transcription factors, and it raises the possibility that **KI-TOX-A3** binds to the transactivation domain (TAD) of TOX to affect its inhibition of PPIs.

**Figure 4.**
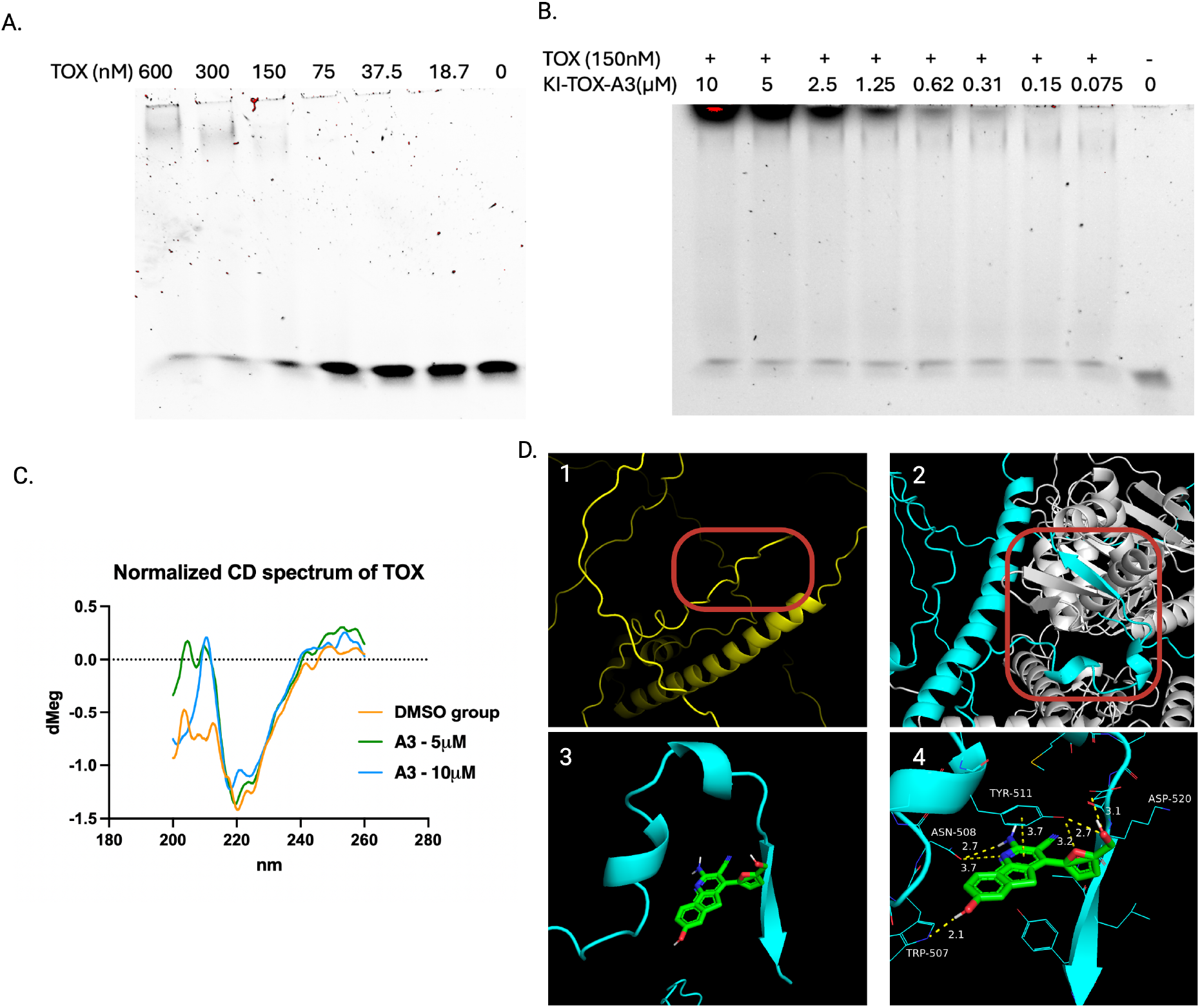
A structure-binding investigation of the current lead **KI-TOX-A3**. (a) EMSA data showed that the DNA probe (sequence provided in the SI) of TOX successfully binds to TOX protein dose dependently. (b) The **KI-TOX-A3** from 0 to 10μM do not inhibit the DNA binding domain function using the DNA probe. (c) The CD experiment shows that **KI-TOX-A3** causes conformational change in TOX protein from free loop to α-helix. (d1) TOX C-term has a free loop, (d2) while the TOX-KAT7 interaction simulation elucidated by AlphaFold v3 shows that the C-terminal changes into a secondary structure; (d3). The KI-TOX-A3 was docked on the C-terminal of TOX where the KAT7 was removed) by Auto dock 1.5.3 with PYRX. (d4) **KI-TOX-A3** showed binding with hydrogen bond formation and π-π interaction with the C-terminal of the TOX.

To understand further how TOX and **KI-TOX-A3** interact, we utilized circular dichroism (CD), a biophysical method used to observe secondary structure changes by the differential absorption of left and right circularly polarized light [38]. Interestingly, in the range of 210nm to 200nm, we noticed a significant upward positive shift of CD signal in the **KI-TOX-A3** treated groups (Figure 4C). A negative signal corresponds to disordered coil, while a positive signal indicates a right-handed α-helix. This change implies that **KI-TOX-A3** induces a secondary structure formation, likely to be an α-helix.

To assess potential ligand binding loci on the TAD of TOX, we executed an AlphaFold v3 prediction on TOX-KAT7 binding and removed the KAT7 protein from the binding site for ligand docking[39]. Our AlphaFold v3 analyses suggest that TOX captures KAT7 via inserting the N-terminus into the KAT7 helix, while the C-terminus of TOX resides nearby KAT7, inducing a conformational change from random coil to helix and beta-sheet (highlighted in red circle and blue line in Fig. 4C). This indicates that **KI-TOX-A3** might have a N- or C-terminal binding site. In addition, we removed KAT7 and ran an unbiased Auto dock simulation [40] with **KI-TOX-A3** on the full-length TOX protein. We noticed that the C-terminus shows stronger ligand binding preference based on the binding affinity calculated from docking. The amino group of the **KI-TOX-A3** does not have direct interaction with the TOX protein, while the phenol and methanol groups form hydrogen bonds with Trp-507, Asn-508 and Asp-520. We also noticed that the pyridine ring forms π-π interaction with Tyr-511, and the oxygen on furan group is a hydrogen receptor of Try-511.

### Chemical SAR revealed potential binding poses

Finally, we performed a pilot structure-activity relationship (SAR) by synthesizing 13 derivatives of the parent compound (Table 1). Synthesis methods are reported in SI of method, and the synthesis scheme is in Fig.S2. To examine the potency of the analogs, we utilized the Cell Titer-Glo (CTG) assay as a surrogate to assess cell viability in the TOX-dependent cell lines Jurkat, Molt-4, HBP-ALL, HH, and Hut78, and TOX-independent cells K562 and Raji. We also utilized the PPI binding inhibition assay (Fig. 1D) as a target-directed measure of activity. Selected modifications such as relocating hydroxyl groups in **KI-TOX-A3-0177** and **KI-TOX-A3-0179**, switching furan to thiophene in **KI-TOX-A3-0182**, to pyrrole in **KI-TOX-A3-0185**, removing the hydroxyl group near the furan in **KI-TOX-A3-0181**, and substituting into benzyl in **KI-TOX-A3-0184** or phenol in **KI-TOX-A3-0186** all decreased cytotoxicity and PPI inhibition. Possible reasons could be the bulky groups hinder interactions, or possibly a loss of essential H-bonding in the interactions. Expanding the five-member ring to a six-member ring in **KI-TOX-A3-0176** also impaired cytotoxicity and PPI inhibition, confirming the critical and fixed nature of the binding orientation. Interestingly, the modifications on the amine with isocyanate extension or amide coupling only mildly impacted TOX PPI inhibition (**KI-TOX-A3-0189, KI-TOX-A3-0190 and KI-TOX-A3-0246**) and slightly improved cytotoxicity against TOX-dependent leukemia cells. To conclude, we found that the SAR results are highly aligned to the docking information, indicating a highly possible binding site of **KI-TOX-A3** at the C-terminus of TOX. Based on this pilot SAR, more lead optimization studies can be done.

**Table 1.**
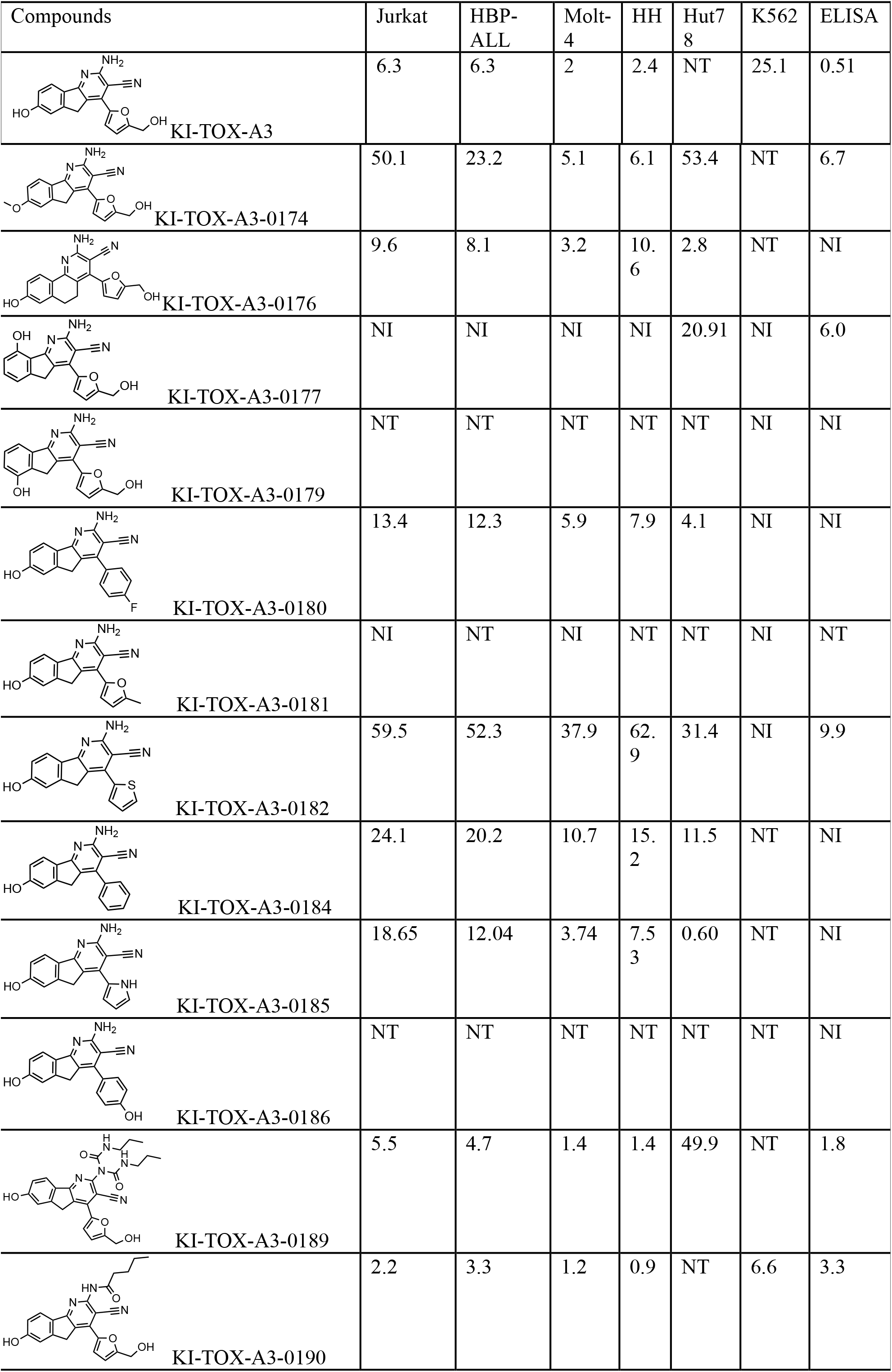
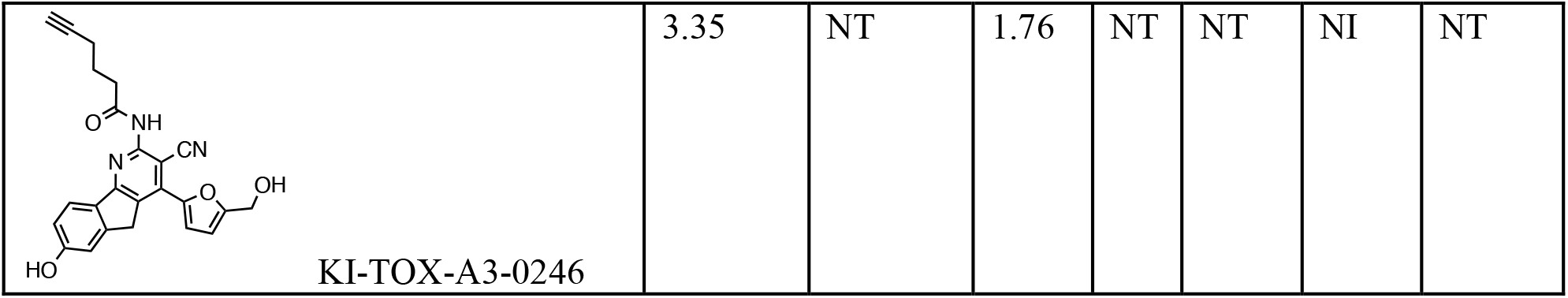
SAR study of the **KI-TOX-A3** (also see Fig. S3). The modification indicates that the -NH2 group is the accessible region of the **KI-TOX-A3**. Both furan and phenyl groups are critical for TOX binding. Unit: μM; NT: Not tested; NI: No inhibition at 100μM (anti-proliferation) or 10μM (ELISA).

| Compounds | Jurkat | HBP-ALL | Molt-4 | HH | Hut78 | K562 | ELISA |
| --- | --- | --- | --- | --- | --- | --- | --- |
| **KI-TOX-A3** | 6.3 | 6.3 | 2 | 2.4 | NT | 25.1 | 0.51 |
| KI-TOX-A3-0174 | 50.1 | 23.2 | 5.1 | 6.1 | 53.4 | NT | 6.7 |
| KI-TOX-A3-0176 | 9.6 | 8.1 | 3.2 | 10.6 | 2.8 | NT | NI |
| KI-TOX-A3-0177 | NI | NI | NI | NI | 20.91 | NI | 6.0 |
| KI-TOX-A3-0179 | NT | NT | NT | NT | NT | NI | NI |
| KI-TOX-A3-0180 | 13.4 | 12.3 | 5.9 | 7.9 | 4.1 | NI | NI |
| KI-TOX-A3-0181 | NI | NT | NI | NT | NT | NI | NT |
| KI-TOX-A3-0182 | 59.5 | 52.3 | 37.9 | 62.9 | 31.4 | NI | 9.9 |
| KI-TOX-A3-0184 | 24.1 | 20.2 | 10.7 | 15.2 | 11.5 | NT | NI |
| KI-TOX-A3-0185 | 18.65 | 12.04 | 3.74 | 7.53 | 0.60 | NT | NI |
| KI-TOX-A3-0186 | NT | NT | NT | NT | NT | NT | NI |
| KI-TOX-A3-0189 | 5.5 | 4.7 | 1.4 | 1.4 | 49.9 | NT | 1.8 |
| KI-TOX-A3-0190 | 2.2 | 3.3 | 1.2 | 0.9 | NT | 6.6 | 3.3 |
| KI-TOX-A3-0246 | 3.35 | NT | 1.76 | NT | NT | NI | NT |

### The TOX inhibitors re-invigorate exhausted T cells

Given that genetic knockout of *Tox* could mitigate T cell exhaustion in previous studies[1], we next tested whether **KI-TOX-A3, KI-TOX-D22** and **KI-TOX-P14** could likewise recover T cell activity in the context of chronic stimulation. First, we developed an *in vitro* T cell exhaustion protocol for mammalian CD8^+^ T cells that is optimized from a previous study [41] (Fig. 5A) (see Methods). In brief, primary CD8^+^ T cells from peripheral blood were stimulated with anti-CD2/3/28 stimulator either once or repeatedly to induce exhaustion. After 5 stimulation events, the T cells were found to have elevated TOX expression (Fig. 5B).

**Figure 5.**
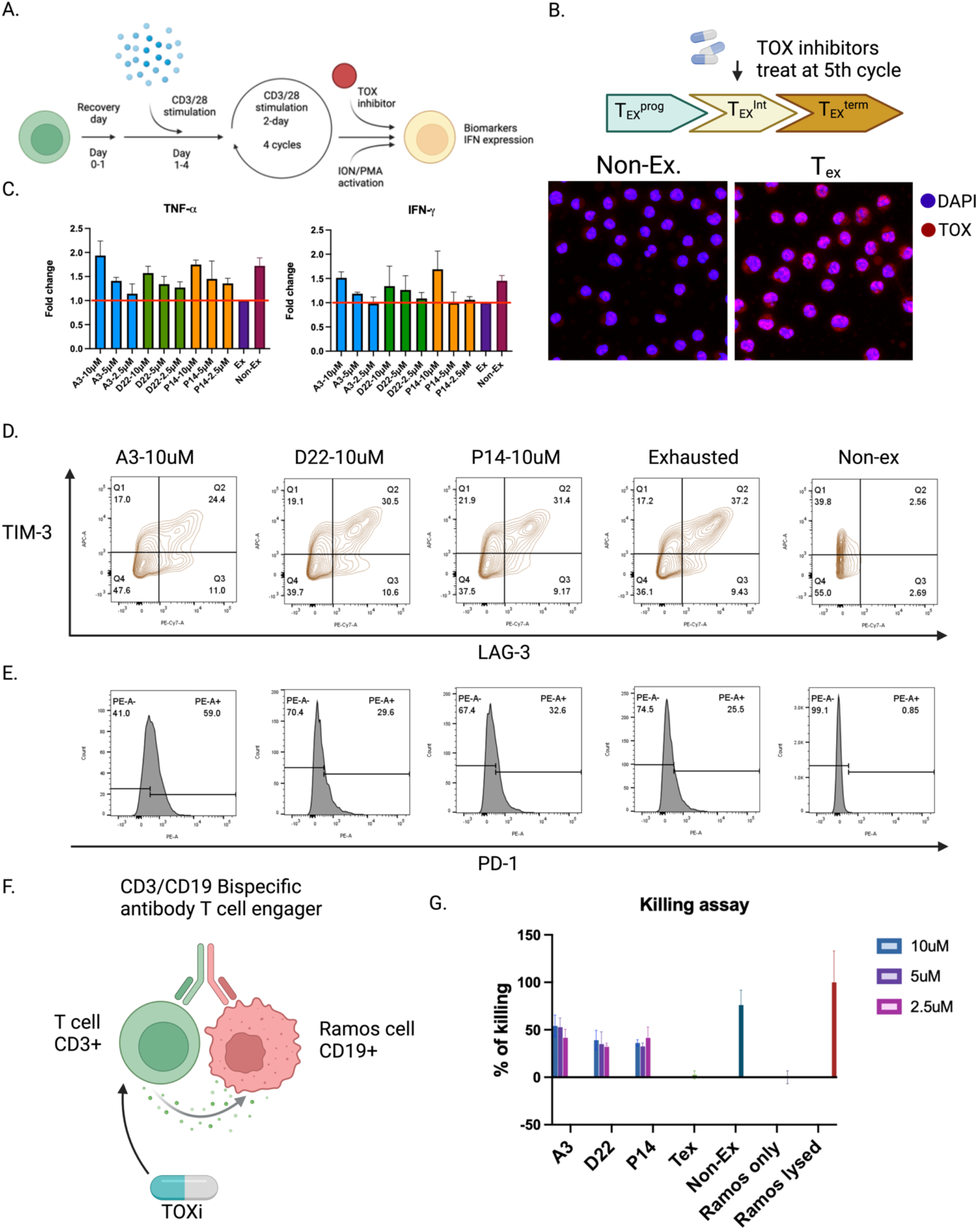
T cell activity was modulated by the TOX regulators. (A). Schematic of the exhaustion protocol to induce CD8^+^ T cell exhaustion in vitro. (B). T cells under 5^th^ cycle of stimulation showed high level of TOX using immunostaining and microscopy. (C). The changes of mRNA of TNF-α and IFN-γ in the T cells. The fold changes are relatively normalized by T_ex_ cells. (D) The inhibitory receptor biomarkers TIM3, LAG3 expression in CD8^+^ exhausted and non-exhausted T cells. (E). PD-1 expression is upregulated by TOX inhibitors (TOXi). (F and G). Co-culture killing assay shows that the T cells after 24h treatment of TOXi can be re-activated and kill the CD19-Ramos cells under the use of Bispecific CD3/CD19 antibody.

We incubated CD8^+^ T cells with the TOX inhibitors. The concentrations used were selected based on results from cytotoxicity assays in figure S6, in which the three compounds did not reduce T cell proliferation at 12.5 µM (24h). After treatment, we assessed the T cell exhaustion biomarkers TIM-3 and LAG-3 and observed reductions of both co-expressed inhibitory receptors after treatment with inhibitors (Fig. 5C). This reduction is consistent with potential for reversing T cell exhaustion state. We also observed similar reductions in co-IR expression following treatment with the TOX inhibitor after the 6th stimulation cycle (Fig. S6E).

Unlike knockdown or knockout of *Tox*, **KI-TOX-A3** increased expression level (Fig.5E) and mRNA levels of PD-1 (Fig. S6C) after 24h of treatment. This was also observed in the 6^th^ cycle stimulation population in Fig. S6E. While the mechanism of upregulation is not yet clear, **KI-TOX-A3** treatment does not eliminate TOX expression and thus performed differently from gene perturbation[42]. Although PD-1 is a known exhaustion marker regulated by TOX[13], it is also recognized as a T cell activation marker and its expression in the absence of other IRs is indicative of enhanced function [43].

To further validate inhibition of exhaustion and recovery of T cell function, we performed a co-culture killing assay[44]. In this assay, we pre-treated T_ex_ cells and non-exhausted T cells with TOX inhibitors at 2.5, 5, and 10µM or 1% v/v DMSO. After the T cells were washed and re-cultured with CD19-Ramos cells along with bispecific T cell engagers (BiTE) to enhance CD3^+^ T cell recognition of the CD19 antigen (Fig. 5F). As expected, T_ex_ exhibited limited cytotoxicity against the CD19-Ramos cells, while the non-exhausted T cells exhibited 80-90% killing. The treated T_ex_ cells demonstrated partial recovery with ∼50% improved cytotoxic activity against CD19-Ramos cells (Fig. 5G). This suggests that TOX is a potential therapeutic target for cancer treatment, and the TOX inhibitors described herein have potential to improve the antitumor activity of exhausted T cells.

### Tox negatively regulates KAT7 function through downregulation of H3K14ac

To further assess how these TOX PPI inhibitors modulate T cell exhaustion, we studied KAT7 activity. As part of the HBO1 complex, KAT7 is critical for T cell proliferation, differentiation, and survival[45]. However, the function of the HBO1 complex in T cell exhaustion is poorly understood. Since we know that the TOX binds to HBO1 complex through direct interaction with KAT7 protein[9], we hypothesized that a TOX-KAT7 interaction may play an important role in T cell exhaustion. Given the critical role of KAT7 in the epigenetic regulation of genes via histone 3 lysine 14 (H3K14) and histone 14 lysine 12 (H4K12) acetylation[46,47], we probed H3K14 acetylation in T cells and found lower levels of acetylation T_ex_ compared to non-exhausted T cells. The treated groups showed dose-dependent recovery of this acetylation (Fig. 6A). We hypothesize that KAT7 plays a critical role in maintaining T cell activation, though more mechanistic studies are needed to discover the function of H3k14 acetylation in T cell activation and exhaustion.

**Figure 6.**
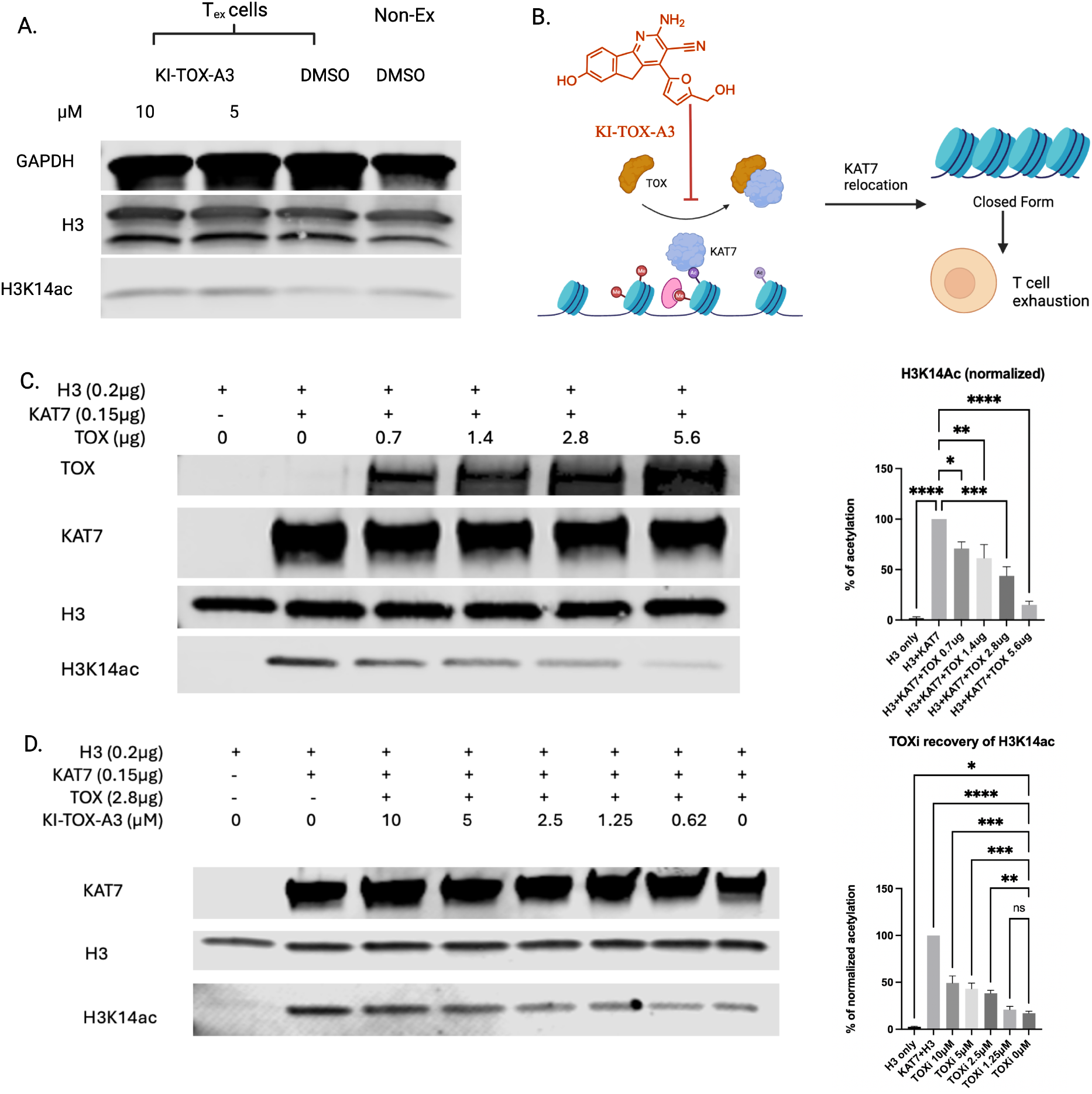
TOX acts as an inhibitor of the KAT7 acetylation activity on the H3K14 site. (A). The exhausted T cells have relatively low expression of H3K14ac, while **KI-TOX-A3** recovered its H3k14ac level to that observed in non-exhausted T cells. (B) Model describing how TOX inhibitors can affect the ability of TOX to interact with KAT7 and alter T cell exhaustion status. (C) *In vitro* assay to study TOX protein as an inhibitor of KAT7 protein by dosing TOX protein in the solution of KAT7, acetyl-CoA and H3 protein, followed by Western blot of H3K14ac, which is regulated by KAT7. (D) The addition of **KI-TOX-A3** helps increase the acetylation on H3 protein after TOX protein adding to the solution.

To this end, we performed an *in vitro* assay to evaluate whether TOX is an inhibitor of KAT7 protein function. Briefly, histone 3 (H3), acetyl-CoA, KAT7 and TOX were mixed and incubated at room temperature in a working buffer reported in method section, followed by a Western blot to monitor H3K14ac levels. We observed that the TOX protein blocks the acetylation function of KAT7. In figure 6B, H3 was successfully acetylated with KAT7 and acetyl CoA mixture, while KAT7 without H3, or H3 without KAT7 does not yield H3K14ac. As hypothesized, the addition of TOX inhibited KAT7 function of H3 acetylation. Therefore, TOX protein may play a role in the epigenetic regulation of T cells through H3k14ac downregulation and followed by chromatin accessibility restriction.

Finally, we added our lead compound **KI-TOX-A3** at various concentrations to explore whether the compound perturbed the TOX-KAT7 interaction. As expected, **KI-TOX-A3** blocked TOX inhibition of KAT7, rescuing KAT7 function as judged by increased H3K14ac levels in this *in vitro* assay (Fig. 6C). From the Western blot, **KI-TOX-A3** has a low micromole IC_50_ similar to the range observed for the compound in the protein binding inhibition data in figure 1F and native gel assay in figure 2B. These data indicate that our TOX inhibitors can recover T cell activity by H3k14 acetylation (Fig. 6D).

## Discussion

Tox is an overexpressed transcription factor that regulates T-ALL leukemia cell growth and drives CD8^+^ T cell exhaustion. Our novel TOX ligand **KI-TOX-A3** modulates TOX protein-protein interactions, including KU70/80 and KAT7, while triggering T-ALL cytotoxicity and inducing CD8^+^ T cell re-invigoration.

The **KI-TOX-A3** chemical probe revealed selective and binding activity in the low micromolar range, intracellularly and biophysically, in figure 2. We were unable to use purified TOX from mammalian or *E. Coli* expression systems for thermal shift assays [48], including DSF, nanoDSF, or cellular lysate thermal shift assays. As mentioned previously, TOX is a disordered protein, and such protein tends to resemble unfolded protein in thermal shift assays. While the CD spectrum for TOX in the presence of compound exhibited an increase of signal at close to 200-210 nm wavelength, indicating a conformational change, we have not fully investigated its binding site(s) for the molecule within the TOX protein. While the DBD is an unlikely site for binding based on our studies, the N- or C-terminal regions of TOX represent potential sites of binding for **KI-TOX-A3**. We aim to understand where this **KI-TOX-A3** could bind in future studies using covalent labeling approach (diazirine labeling)[49].

**KI-TOX-A3** induced cell-type selectivity in cell viability assays involving TOX-dependent cells, suggestive of a target-specific effect. Compound treatment also led to an S phase cell cycle arrest phenotype that resembles the shRNA knockdown of *Tox* in previous studies[1]. We also observed proteasome-dependent reductions in TOX protein levels in the presence of **KI-TOX-A3** after 12 hours. Interestingly, preliminary interactomics experiments using pulldown assays coupled to mass spectrometry identified chaperones, including HSP90AB1, as TOX interactors that are stripped in the presence of compound. These factors may serve to stabilize the conformationally plastic transcription factor, and their removal leads to misfolding followed by degradation. The mRNA level of TOX is modulated slightly by 4 to 6 hours.

We next sought to evaluate the potential of the compounds to impact CD8^+^ T cell exhaustion based on the role that TOX plays in this process[9]. **KI-TOX-A3** and two additional candidates from our SMM assays appeared to exhibit a T cell re-invigoration effect as judged by downregulation of co-expressed inhibitory receptors (TIM-3 and LAG-3) and upregulation of the cytokines TNF-α and IFN-γ. Note that all three candidates showed significant upregulation of the *pdcd1* and *Nr4a1* gene in the mRNA level (Fig. S6C and D), which are further suggestive of acute T cell stimulation and activation[50]. Previous research indicates that the *Tox* knockout is correlated to PD-1 and NR4A1 downregulation in the mice T cells[9] and human CAR-T[42]. Although genetic alteration could be different from transient inhibition, more direct molecular mechanism studies on **KI-TOX-A3** regulation of PD-1 and NR4A1 are needed. We also explored the compounds in co-culture killing assay[44] with the T cell engager BiTE and concluded that the TOX modulators could efficiently engage T cell killing activity, even though the T cell was exhausted and originally incompetent to kill cancer cells.

With *Tox* gene knockout, the accessibility of more than 4,000 chromatin sites were altered -and epigenetic regulation of key genes was demonstrated [9]. The precise role of TOX in regulating genesis is unclear. To critically think about the role of TOX in T cell exhaustion, one objective is to understand how TOX regulates T cells through epigenetic regulations[9]. Currently, there is no solid evidence that TOX could drive transcription activity by itself. Since our compounds inhibit the TOX-KAT7 interaction, we assessed whether TOX impacts KAT7 protein function. In the paper, we probed H3K14 acetylation levels in T_ex_ cells and found that **KI-TOX-A3** could recover the acetylation of H3K14 globally to the level of non-exhausted T cells. This implies that KAT7 might have a critical maintenance function in T cell activation via H3K14 acetylation, and overexpressed TOX in exhausted T cells is an antagonist of KAT7, which will decrease the H3K14ac. We further performed this *in vitro* assay to clearly show that H3K14ac was decreased with TOX titration, while the TOX inhibitor **KI-TOX-A3** recovers the KAT7 function and increases the H3K14ac level. This implies that TOX can modulate gene expression via HAT recruitment.

## Conclusion

Tox is an intrinsic disordered transcription factor that regulates both cancer proliferation and T cell exhaustion. TOX has not been targeted as it is regarded as ‘undruggable’ protein. Currently, the mechanistic studies of this protein have been dependent on gene perturbation studies, which has its limitation in many transient mechanism studies. With the discovery of this novel and potent TOX inhibitor **KI-TOX-A3**, we could understand more epigenetic regulations that TOX performs.

## Acknowledgement

B.W. and A.N.K. acknowledge support from the MIT Center for Precision Cancer Medicine as well as the National Cancer Institute (NCI), including center support grant P30-CA014051 and U54-CA231630. P.M.K.W. was supported by grants from the National Institute of Health (NIH), NCI support grant P30-CA045508 and NCI K22CA279501, the V Foundation for Cancer Research, the Betty Ajces Trust, and the Wasily Family Foundation. D.J.I. and A.H. were supported from the Mark Foundation for Cancer Research. This work was supported in part by the Koch Institute Support (core) Grant P30-CA014051 from the National Cancer Institute. A.C. acknowledge supports from US. Department of Defense, Defense Threat Reduction Agency (DTRA) HR00112120010, National Institute of General Medical Sciences (NIGMS) R01GM137606, and National Institute of Diabetes and Digestive and Kidney Diseases (NIDDK) U01DK137242.

We benefitted from the guidance of Stephanie Gaglione and Michael Birnbaum at MIT, including donation of PBMC samples and sharing protocols related to T cell culture and flow cytometry analyses. We received T-ALL cell lines from Professor David Langenau at MGH. Ben Leu provided technical support for SMM screens. We thank Yumeng Liu, protein specialist from University of Wisconsin-Madison, and Sunbin Deng, a postdoc from Harvard Medical school, for the professional advice related to protein expression and purification. We thank Adgeboyega Yomi Oyelere, professor at Georgia Institute of Technology, with all his professional guidance and suggestions in chemistry.

