## Supplementary material for "Small molecule modulators of TOX protein re-invigorate T cell activity": materials and method

Bocheng Wu[1,2], Heng Jui Chang[1,2], Prashant Singh[2], Alexander Hostetler[1,3,4], Yichen Xiang[1,3], Shenghao Guo[1,3], Fiona Yihan Wang[3], Julia J. Zhong[3], Becky S. Leifer[1,2], Richard P. Schiavoni[1], Nan Jiang[1,2,3], Amit Choudhary[2], Peter M. K. Westcott[5], Angela N. Koehler[1,2,3*]
[1]Koch Institute for Integrative Cancer Research, Massachusetts Institute of Technology, Cambridge MA 02139
[2]Broad Institute of Massachusetts Institute of Technology and Harvard University, Cambridge, MA 02142
[3]Department of Biological Engineering, Massachusetts Institute of Technology, Cambridge, MA 02139
[4]Department of Immunology and Microbiology, The Scripps Research Institute, La Jolla, CA 92037
[5]Cold Spring Harbor Laboratory, Cold Spring Harbor, NY 11724

*Corresponding Author

**Key word:** T cell exhaustion, T cell acute lymphoblastic leukemia, TOX, small molecule microarray, intrinsically disordered protein, protein-protein interaction, transcription factors, DNA-binding domain, transactivation domain, cytokine, KAT7, histone acetylation, epigenetics

**Materials:**

**Cells:**

| Cell name | Lab or Company | Part number |
|---|---|---|
| Jurkat | Langenau lab from MGH | N/A |
| Molt-4 | Langenau lab from MGH | N/A |
| HH | ATCC | CRL-2105 |
| Hut78 | ATCC | TIB-161 |
| K562 | ATCC | CRL-3344 |
| HBP-ALL | Langenau lab from MGH | N/A |
| Raji | Koch Preclinical core | N/A |
| Human Peripheral Blood $CD8^+$ T Cells | STEMCELL | 200-0164 |
| BL21 (DE3) | NEB | C2527H |
| BL21, C43 (Lemo21) | NEB | C2528 |
| Ramos | Irvine Lab | N/A |

**Plasmid:**

*Mammalian cell expressed TOX plasmid construction*

vectorbuilder VB230807-1552yxc.
pRP[Exp]-EGFP/Puro-EF1A>Avi/3xGGGGS
/hTOX[NM_014729.3]/3xGGGGS/3xFLAG

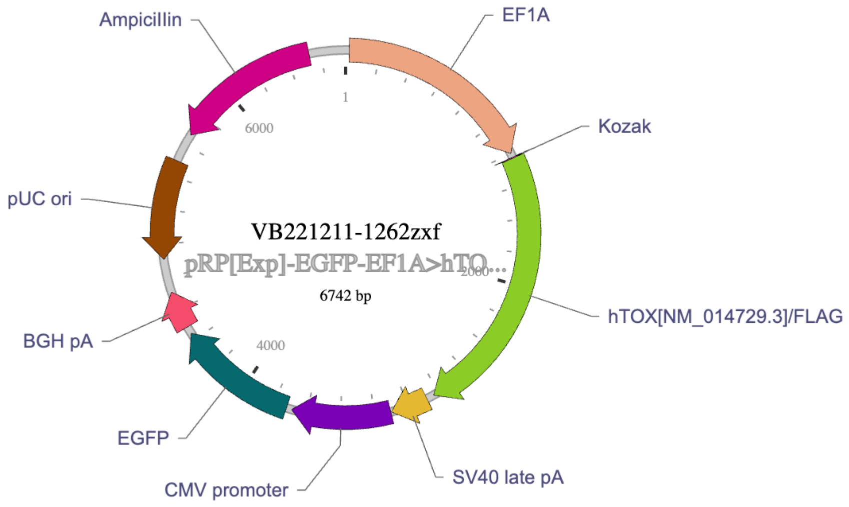


*E.Coli plasmid:*
Backbone Vector pSMT3 was cut from plasmid 191261 of Addgene. Inserted TOX DNA was made by Twist Bio.

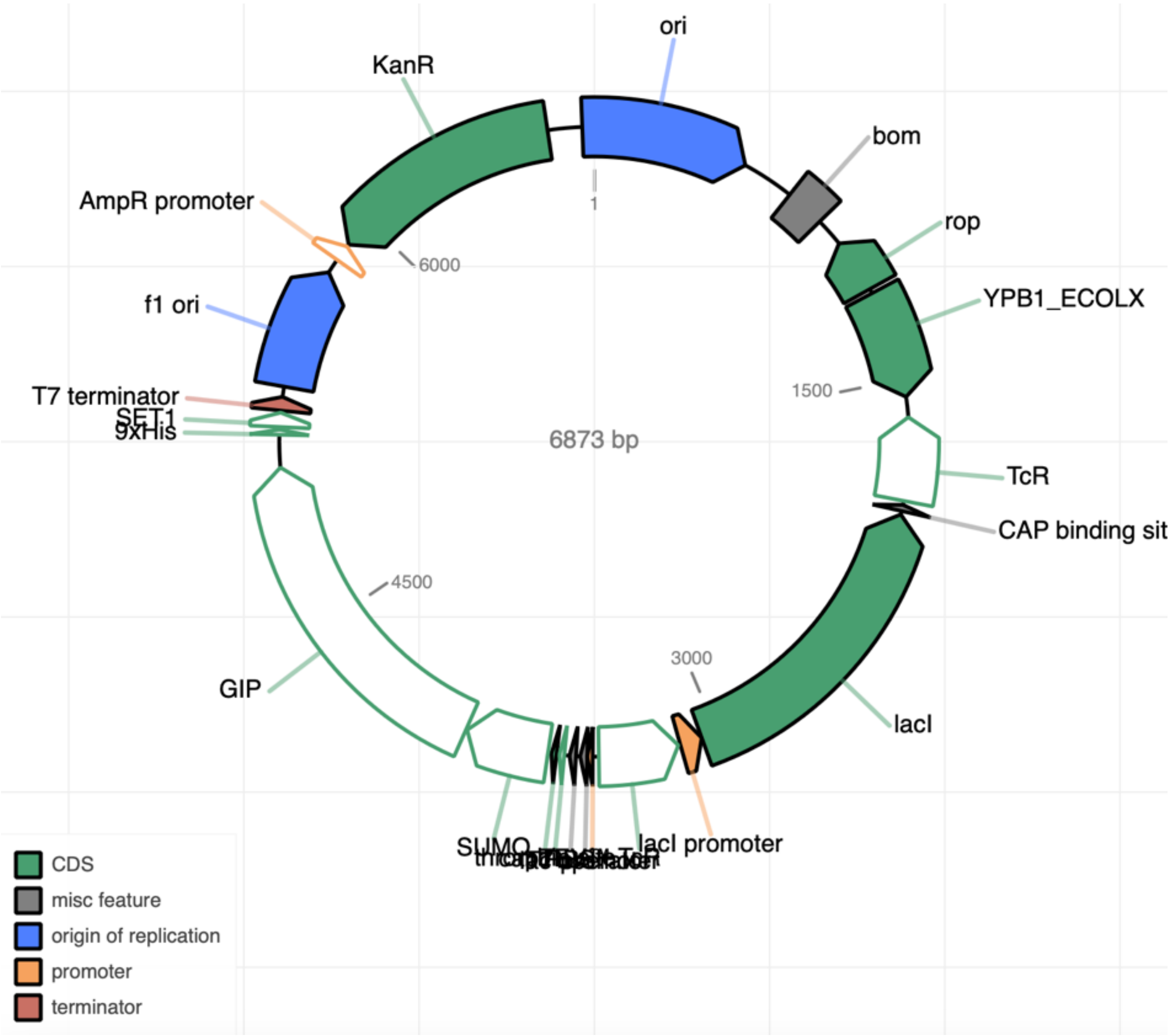


### Primers:

| | |
|---|---|
| TOX | FW: CGCTACCTTTGGCGAAGTCTCT |
| | RS: CTGGCTCTGATATGCTGCGAGTT |
| CDK9 | FW: CTGAAGAAGGTGCTGATGGAA |
| | RS: AGTTGACCACATTCTCGTGTT |
| Myc | FW: TCCTCGGATTCTCTGCTCTC |
| | RS: TCTTCCTCATCTTCTTGTTCCT |
| RUNX3 | FW: GGCAATGACGAGAACTACTCCG |
| | RS: GATGGTCAGGGTGAAACTCTTCC |
| NR4A1 | FW: GGACAACGCTTCATGCCAGCAT |
| | RS: CCTTGTTAGCCAGGCAGATGTAC |
| PD-1 | FW: AAGGCGCAGATCAAAGAGAGCC |
| | RS: CAACCACCAGGGTTTGGAACTG |
| GAPDH | FW: GTTCAGGAAGAGTGACACCA |
| | RS: TTCTCCGCATCTCCATTCTC |
| TNF-α | FW: CTCTTCTGCCTGCTGCACTTTG |
| | RS: ATGGGCTACAGGCTTGTCACTC |

| | |
|---|---|
| LAG3 | Hs.PT.58.28328449 (IDT PrimeTime® qPCR Primers) |
| TIM3 | Hs.PT.58.916495 (IDT PrimeTime® qPCR Primers) |
| CTLA-4 | Hs.PT.58.3907580 (IDT PrimeTime® qPCR Primers) |
| Sanger sequencing primers | FW: 5'-GCGGATCCATGGGTTTAAATGACATATTCG-3' |
| | RS: 5'-GCCTCGAGTTAGCTGCCACTG-3' |
| Duplex oligo of TOX EMSA DNA probe | 5'-CGGCGCCCCCGCCCGC-3' |

### Protein/Peptide

| **Protein/peptide** | **Source of expression** | **Tag** | **Company** | **Part** |
|---|---|---|---|---|
| Ku70/80, human | Baculovirus-Insect Cells | His | Sino Biological | CT018-H07B |
| KAT7, human | Sf9 cells | FLAG | Active Motif | 31489 |
| TOX, human | HEK293T | FLAG | Origene | TP303792 |
| Sumo protease | *E. coli derived* | His | Trialtus bioscience | N/A |
| BirA enzyme | N/A | N/A | Avidity LLC | N/A |
| XhoI | N/A | N/A | NEB | R0146S |
| BamHI | N/A | N/A | NEB | R3136S |
| IL-2 | *E. coli-derived* | N/A | R&D system | 202-IL-050/CF |
| 3xFLAG peptide | *N/A* | N/A | LifeTein | LT8153 |
| IRDye® 680RD Goat anti-Rabbit IgG Secondary Antibody | *N/A* | N/A | Licor | 926-68071 |
| IRDye® 680RD Goat anti-Mouse IgG Secondary Antibody | *N/A* | N/A | Licor | 926-68070 |
| IRDye® 680RD Streptavidin protein | *N/A* | N/A | Licor | 926-68079 |

### ELISA binding assay

Nunc® Immobilizer™ Amino Plates and Modules (Thermo Scientific); Chemiluminescent SuperSignal™ ELISA Femto Substrate (Thermofisher); chemicals (Chemibridge); Ethanolamine (Thermofisher). Superblock Blocking buffer (Thermofisher). mTOX recombinant protein (Origene).

**SPR:** Cytiva S sensor SA chip. Biacore™ T200 SPR system.

**Antibodies:**

| Antibody Target | Vendor | Part number |
|---|---|---|
| TOX1/2 | CST | E6G5O |

| | | |
|---|---|---|
| TOX1 | Abcam | ab237009 |
| RUNX3/AML2 | CST | D6E2 |
| H3 | CST | 9715 |
| Acetyl-Histone H3 (Lys14) | CST | D4B9 |
| Nur77 | CST | D63C5 |
| GAPDH | CST | 14C10 |
| Anti-mouse HRP | CST | 7076 |
| Anti-rabbit HRP | CST | 7074 |
| HSP90AB1 | Proteintech | 11405-1-AP |
| DYKDDDDK-tag Antibody, pAb, Rabbit | Genscript | A00170 |
| KAT7/MYST2 | Thermofisher (Invitrogen) | MA5-48711 |
| PE PD1 | Biolegend | 367403 |
| APC TIM-3 | Biolegend | 364803 |
| PE-cy7 LAG-3 | Biolegend | 125225 |
| PE-cy7 IFN-y | Biolegend | 502527 |
| Alexa Fluor® 594 TOX | Biolegend | 682604 |
| Zombie Violet™ Fixable Viability | Biolegend | 423113 |

CST=Cellsignaling

**PLA:** The Duolink® Proximity ligation assay was purchased from Sigma (DUO92101). Olympus FV1200 Laser Scanning Confocal Microscope. 35 mm Dish (No. 1.5 Coverslip 20 mm Glass Diameter, MATTEK, P35G-1.5-20-C-HA). Cell fixation paraformaldehyde (Thermofisher).

**Methods:**

### Mammalian Cell Culture:

Cancer cells: Jurkat cell, Molt4, HBP-ALL, K562, Raji, HH cell lines were incubated with RPMI 1640 media with 10% FBS under 37°C and 5% $CO_2$. The Hut78 cells were incubated with IMDM and 20% FBS in the same incubation condition. HEK293-T cells were plated in T-75 culture flask (Corning) in growth media (DMEM supplemented with 1% penicillin-streptomycin solution and 10% FBS) at a density of 1 x $10^6$ cells per flask.
CD8$^+$ T cells: ImmunoCult™-XF T Cell Expansion Medium with 50unit/ml IL-2.

### *Mammalian TOX expression :*

A plasmid with pRP contained both promoter EF1A for the full length human TOX (Avi tag and 3xGGGGS at N-terminal and 3xGGGGS and FLAG tag at C-terminal) and CMV promoter for EGFP as the transfection indicator. We extracted the plasmid and transfected HEK293-S Gn- cells using PEI-MAX (Polysciences, 49553- 93-7) and plasmid at the ratio 3:1 where the DNA is 20ug/5million cells. In day two of the transfection, we added peptone primatone as the nutrition to the cells. The transfection lasted for 96 hours, and the cells were collected. Then, the 50 million cells were washed by 1X PBS and lysed using 5ml IP-lysis buffer (Thermofisher). The cell lysate was vortexed for 3-4 times (15s/time) in 15 minutes (the rest of the time in ice). Then, Anti-FLAG® M2 Magnetic Beads were incubated to the lysate for 2h in cold room with rotation. Then, the beads were washed one time by regular washing buffer (TRIS 20mM, NaCl=150mM), then one time with high salt washing buffer (TRIS 20mM, NaCl=1000mM), and one time again with regular wash buffer. The FLAG protein was eluted by 1xFLAG peptide (0.75mg/ml). The peptide in the eluted proteins was removed by 10k centrifugal filter (Sigma) 4 times to exclude the peptide. The yield of mTOX is around 10ug per 10million cells.

#### *E. Coli* recombinant protein expression

*1. Plasmid vector, cloning, and transformation:*

The backbone vector restriction sites of the pSMT-3 were recognized and cut by BamHI and XhoI. The protocol follows the suggested restriction digest protocol of NEB. Briefly, the 0.5μg of plasmid was dissolved in the ultrapure water to make a solution of 10uL. Then, 2uL of 10x NEB buffer was added. Then 1uL of each restriction enzyme was added to the mixture and fill the solution to 20uL with ultra-pure water. The solution was incubated at 37°C for 2h, followed by 85°C denature and cooldown to 12°C. The backbone has been cut and is ready to be purified. The gel purification was applied using the NEB gel purification kit. By determining the concentration of the backbone band using NanoDrop, we calculated a 1:4 mixture of insert: backbone amount. Then, after mixing insert and backbone, we added a 10x ligation buffer and 1ul of T4 ligase. The ligation was incubated in the 4-8°C refrigerator on the door side. On day 2, the NEB stable *E. coli* strain was defrosted under ice. Then, 50ng of the mixture was added to the cells. After an additional 10-minute incubation in ice, the cell/DNA mixture was placed in 42°C water bath for 45s. Then, the cells were recovered by growing with SOC outgrowth media at 37°C for 1h. The cells were diluted by 1:10, 1:100, and 1:1000 in LB broth. Then, three prepared kanamycin (30μg/ml) added LB agar plates were used to culture 50uL of each dilution. The cells were grown at 37°C overnight. We found that 1:100 was the optimum condition with good number of colonies. We picked 15 colonies, grew them, and extracted the DNAs for nanopore analysis. Colony #5 has clear His-SUMO-AVI-TOX DNA insert and no impurities. We chose this colony for protein expressions.

*2. Protein expression:*

The c43 were transformed with the chosen DNA and the cells were grown in the LB Agar plate that contains both kanamycin (30μg/ml) and chloramphenicol (30g/ml) to maintain the Lemo21 and TOX plasmid. The colony was picked and examined with Sanger sequence to double confirm the existing TOX plasmid.

The c43 were cultured and induced in multiple conditions: 18°C overnight, 30°C 5h, and 37°C 2hr and with 0mM, 0.25mM, 0.5mM, 1mM, 1.5mM, and 2mM of L-rhamnose. IPTG=0.4mM was added to the OD=0.8 in LB broth media. Each condition was incubated in the 10mL bacterial culture tube with 5mL of cells, and 1mL cells were collected and pelleted when the induction was ended. The pellets were cooked under 100°C with 4x Laemmli buffer and 2v/v% of BME. We selected the 37°C 2hr with L-rhamnose=0mM as we observed maximum amount of TOX expression in the Coomassie gel.

To proceed the large scale, we cultured the cells in 1L of LB broth media (Kan/Cam=30μg/ml) with 250RPM 37°C. Up to OD=0.8, 25ml of the cells were collected, mixed with 10% glycerol, and reserved in -80°C for future protein expression. The rest of the cells were mixed with IPTG powder to make a final concentration of 0.4mM. The cells continued to grow under 37°C for 2hr. The cells were centrifuged into pellets (4000 xg 10mins) and washed again by 1X PBS buffer. We gained approximately 4g of bacteria. We stored the pellets at -80°C until day 2.

The cells were ready to be lysed. We defrosted the pellets and lysed the cells with 20mL of Bug buster master mix buffer (Sigma) and 4 tablets of cOmplete EDTA-free Protease inhibitor (Sigma). The lysates were incubated on a head-to-end rotator at room temperature for 30 minutes (or 1h at cold room). The lysate turned from viscous and cloudy to clearer with less viscosity. The lysate was sonicated (15s on, 20s off, 60%

Amp) in ice bath for 5 cycles. The cells were centrifuged (21,000 xg 45mins). The supernatant was removed, and the pellet was collected. Then, an inclusion body wash buffer (100mM Tris, 2% Triton 100, 5mM DTT, 2M Urea) was used to wash the pellet by pipetting. The pellet was further incubated by the rotator at room temperature for 7mins. Then, the lysate was centrifuged again at 21,000 xg for 30 mins. The pellet was washed again for 2 repeats and at the end of the third wash the pellet should be white. Then, the last wash was applied with the IB wash buffer 2 (100mM Tris, 5mM DTT). Then, the denature eq. buffer (20mM sodium phosphate, 300mM sodium chloride, 6M guanidine•HCl, 10mM imidazole; pH 7.4) was used to equilibrate the NTA resin (25217, Thermofisher) and to dissolve the inclusion body. The inclusion body was dissolved in 4ml of the eq. buffer and mixed with 0.5ml of equilibrated resin. The mixture was incubated on the rotator for 15-20mins at room temperature and was filtered using the 10mL spin column. The resin was further washed by wash buffer A (10mM HEPES, 300mM NaCl, 2mM DTT, 20mM imidazole, pH=7) and wash buffer B (10mM HEPES, 1M NaCl, 2mM DTT, 30mM imidazole, pH=7) with 20 equivalent volume of resin. The spin column was dried by a quick spinning (1,000 xg, 1 minute). Then, the TOX elution was applied by incubating 1mL of elution buffer (10mM HEPES, 300mM NaCl, 2mM DTT, 500mM imidazole, pH=7) at room temperature for 5 minutes with shaking. Then, the column was spun for 1,000 xg 1 minute to obtain the first elution batch. We repeated the elution 2 more times and combined the elution.

*3. Dialysis:*

The dialysis buffer (20mM HEPES, 200mM NaCl, pH=8) was applied. Note that our protein has a PI value around 7.4 which the PBS (pH=7.4) dialysis will cause protein aggregation dramatically. We recommend using this buffer for protein storage. The protein was dialyzed overnight at 4°C and is ready to use.

*4. SUMO-tag removal:*

The dialyzed protein (500μg) will be incubated with 50μg SUMO Protease (Trialtus bioscience) in 4°C with head-to-end rotation overnight. In day 2, we pre-incubated and equilibrated 0.1g of NTA resin with 10mM imidazole contained TOX buffer. Then, the resin was co-incubated with the SUMO-removed protein mixture for 10 minutes at room temperature. Subsequently, we filtered out the resin by centrifuge column and the filtrate is the pure Avi-tagged TOX protein (250μg).

*5. Biotinylation:*

Different from other processes, the BirA enzyme lost its activity by 50% in 100mM NaCl or higher. Therefore, as an alternative buffer condition, we dialyzed the Avi-TOX in the BirA preferred buffer (10mM HEPES, 100mM potassium glutamate, pH=8.1). The protein (500μL, 0.5μg/ul) was mixed with 5μg BirA enzyme and 50μL of BirA buffer A, B, and C that were provided by Avidity (BirA500: BirA biotin-protein ligase standard reaction kit). The reaction was maintained on the rotator overnight in the cold room. Then, the biotinylated protein was desalted by the Zeba column (Thermofisher) to remove 90% of biotin. The column can perform buffer exchange by recharging the column with TOX buffer (20mM HEPES, 300mM NaCl) by spinning 4 times of 5ml of this buffer. Then loading our sample and centrifuge. The sample was spun at 1500 xg for 5 minutes and the filtrate was the crude biotinylated TOX protein. We still noticed 10% biotin in the protein solution and therefore a dialysis to TOX buffer was performed. The final yield of the

TOX protein was around 100μg (40%). This biotinylation was validated by western blot, with a IRDye® 680RD Streptavidin protein staining (Licor, P/N 926-68079).

### Small molecule microarray

*1. Screening Preparation:*

Printing of small molecule microarray slides follows the previous manufacturing methods (Clemons et al., 2010). The setting of the screening is as follows: we spitted 65,000 chemicals into 16 sets and therefore each set has around 4,000 chemicals. One slide would contain all 4,000 chemicals that are replicated coated, alone with 2,000 spots that belong to DMSO or control (rapamycin). Overall, there are ~10,000 chemicals coated in a single slide, which stand for 1 set of chemical libraries. Lysate preparation by harvesting HEK293-T cell at 90% confluency, washing, lysis with RIPA and concentration determination by BCA. The concentration of lysate protein reached nearly 20mg/ml concentration. The cell debris was removed after centrifuge and the supernatant were aliquoted to 1.25mg/tube. Each 1.25mg of lysate would be applied to one slide screening.

*2. SMM Screening:*

First, add 2.5mg lysate and 50μL cocktail into the 5ml of MIPP buffer. Also prepare 1.25mg lysate and 25μL cocktail to 2.5ml MIPP for background solution. Then, add 22μL TOX (5μg) into 5 ml of MIPP buffer (20 mM $NaH_2PO_4$, 1 mM $Na_3VO_4$, 5 mM NaF, 25 mM β-glycerophosphate, 2 mM EGTA, 2 mM EDTA, 1 mM DTT, 0.5% Triton X-100, pH 7.2, cOmplete™ protease inhibitor cocktail tablets), mix well. Place the slides into the 4-well plate and label well with the barcode. Mark background (The chip incubates with buffer that does not contain TOX). Incubate the slide with the prepared TOX solution (sample) or blank buffer solution for 1 h (negative control). Then, pour the buffer and wash the slides with 1x TBST for 2min 3 times. Subsequently, dilute antibody into the TBST solution by 1:2000 and mix well. Carefully add the prepared antibody to the slides (2.5ml per chip) and incubate for another 1h. Later, the slides were washed by TBST for 2mins and repeated 2 times. Then wash with TBS buffer for 2 mins once. Last, the slides were dipped into water and washed, followed by drying up with centrifuge. The signals on the slide are now ready to be detected. The slides were inserted to the machine GenePix 4000B fluorescence scanner (Molecular Devices) with barcode facing down. The slide images were analyzed using GenePix Pro software to produce raw data for statistical analysis.

### ELISA assay:

First, protein KAT7 (5μg) mixed with 10ml $Na_2CO_3$:$NaHCO_3$ (100mM, pH=9.6) and incubate to the Nunc Amino 96-well white plate (Thermofisher) for 2 hours at room temperature with shaking. Each well contains 100μL of the KAT7 buffer. Then, gently pipet out the solution and move to deactivation. To do so, 10mM ethanolamine in $Na_2CO_3$:$NaHCO_3$ buffer solution (100mM, pH=9.6) was added to the plate with 150μL per well. The incubation lasts for 30 mins to 60 mins at room temperature with shaking. The following step is blocking. Note that the blocking is very critical as the ethanolamine can cause many non-specific bindings to the TOX or other disordered protein and therefore a blocking step can minimize the side-effect. We blocked the plate with Superblock blocking buffer (Thermofisher) for 2hrs at room temperature (200μL/well). While waiting for blocking, we prepared the protein of interest (10μg TOX per plate) with 10 ml PBS (pH=7.8). Then aliquot 205μL into each well of a regular 96-well plate. Then, add around 2.1μL of your compound of interest to the wells, record the location and chemical name, mix well by pipetting. The incubation of chemical-TOX mixture took around 10-15 minutes before adding to the pre-coated ELISA plate. When blocking is done, wash the plate with PBS-T 2 times with 250μL/well and remember to up-down the pipet to completely wash away the blocking buffer. Then, use TBS to wash again. In each time of wash, let the plate stay 2-3 minutes on the shaker can clean up the wells better. After the complete removal the TBS, add the TOX-chemical mixture to this blocked plate (100μL/well and replicate) and incubate 45-60mins at room temperature with shaking. After incubation, wash the plate lightly with TBS-T and TBS gently (200µL/well). After wash, add primary antibody with 1:1000 dilution to the TBS buffer and mix with a 5x antibody diluent (Thermofisher) as the mild blocker. We incubated the primary antibody overnight at 4°C (1h at room temperature was also fine in another project). Then, on day 2, wash away the primary antibody with both TBS-T and TBS (200µL/well) and add the1:2000 diluted 2$^{nd}$ Rabbit antibody with HRP (Cell Signaling) in the same buffer condition as the primary antibody. Incubate 30-60mins at room temperature. Consequently, the plate was washed again with 300µL/well of volume of TBS-T for 2 times and 400µL TBS as the last time. The complete wash and complete removal of TBS is very critical as any left-out antibody can cause a significant error in the machine reading. After the wash, add chemiluminescent substrate (Thermofisher) at 100µL /well. The plate was read through the Tecan 200 reader without long incubation. In the read, we chose the Nunc Plate of Thermofisher (the choice of plate is very critical that a mistake could lead to read error).

#### Proximity Ligation Assay:

The PLA assay follows the manufacture's protocol in general, except that our experiment was done in Eppendorf tubes rather than microscopic or culture plate. In brief, Molt4 cells were treated with **KI-TOX-A3, KI-TOX-D22,** or **KI-TOX-P14** at 10 and 5 µM or 24h. The cells were collected, washed, and fixed with 3.7% formaldehyde. Then, the cells were washed and stained with anti-TOX (Rabbit) and anti-KAT7 (Mouse) antibodies overnight at 4°C with 1:100 dilution in the 1x antibody diluent. The cells were centrifuge and washed again and stained with anti-mouse minus (Sigma) and anti-Rabbit plus (Sigma) in antibody diluent again for 2hrs at room temperature with rotation. The cells were washed by PLA buffer A and incubated in DNA ligation assay for 30 minutes at 37°C. Last, the fluorescent signal would need to be installed and amplified via the polymerase and amplification buffer under 37°C for 100 minutes. After the amplification, the cells were washed by the PLA buffer B by following the manufacture protocol and the cells were resuspended by Duolink® PLA Mounting Medium with DAPI for 15minutes. The cells were then pipetted and seeded on the microscopic plate (No.1.5 of glass bottom dish, MATTEK). The plate was ready for imaging using the Olympus/Evident FV1200 confocal microscope.

#### Surface Plasmon Resonance:

Binding kinetics were performed in an Assay buffer containing 1X PBS, 0.05% Tween-20 using a multicycle kinetics protocol in a Biacore T-200 (Cytiva, product no 28975001) at 25°C. Running buffer was prepared by supplementing 2% DMSO (reagent grade, Sigma, Cat # 472301) to 1X Assay buffer.

Test compounds **KI-TOX-A3** and **P14** were diluted from 10 mM DMSO stocks to a 1 mM intermediate stock in DMSO and further diluted to varying concentrations (10 µM to 0.15 µM) with a 2% DMSO as final concentration in 1x assay buffer. 100 µL of each compound dilution were transferred to 384 deep well plates (Greiner bio-one Cat# 781270), secured with the microplate sealer (Cytiva, Cat# BR100577), and centrifuged at 800 RPM for 1 minute at room temperature.

The system was primed with a running buffer before immobilization and kinetics measurements. Series S sensor chip SA (Cytiva, Cat# BR100531) was used for immobilizing Biotnylated Tox on the chip using biotin-streptavidin capture chemistry. Sensor chip surface activated with 3 injections of 50mM sodium hydroxide in 1M Sodium chloride solution for 60 sec. 200 nM of biotinylated Tox was prepared in 1X assay buffer and target immobilization of 2150 RU was performed on channel 2, while channel 1 was kept as blank. Post immobilization, both the channels' surface was stabilized by injecting 1X assay buffer for 60 seconds. 15 startup cycles were performed at a flow rate of 40 µl/min to stabilize the baseline. Compounds were injected at a flow rate of 35 µl/min for an association time of 60 seconds and dissociation data was collected for another 90 seconds. A blank injection containing a running buffer was injected after every 7 cycles. The injection port was washed with 50% DMSO after every injection. Multiple Solvent correction

run was performed by a varying percentage of DMSO from 1.25-2.75% with a fixed gradient of 0.25% in the assay buffer, injecting at the beginning and after every 35 cycles of kinetics.
Data processing was performed using Biacore evaluation software 3.2. The solvent correction was applied, followed by kinetics evaluation through reference subtraction using channels 2-1. Sensorgrams were fit to a simple 1:1 kinetic binding model to obtain binding parameters such as association constant (ka), dissociation constant (kd), and binding affinities ($K_D$). The final sensorgram along with fitting curves were plotted using GraphPad Prism version 10.1.2.

### Isothermal Titration Calorimetry (ITC)

The His-SUMO-TOX protein expressed from E.Coli was dialyzed after expression with the working buffer (HEPES 20mM, NaCl 150mM, pH=8) and concentrated to 10μM. Then, the dialysis buffer will also be collected for background titration. Later, 5% DMSO solution of 100μM KI-TOX-A3 was prepared using this dialysis buffer. To eliminate buffer mismatch, the protein solution was also added with DMSO to final 5V/V%.

ITC experiments were conducted using a MicroCal PEAQ-ITC instrument (Malvern) at 25 °C. The protein solution (10 μM) was loaded into the cell, and the small molecule ligand (100 μM) was prepared in the same buffer and titrated in 2 μL injections at 150-second intervals. Heat changes per injection were recorded and integrated after baseline correction. Binding isotherms were analyzed using a one-site binding model with the Microcell PEAQ-ITC analysis software to determine the dissociation constant (Kd), enthalpy (ΔH), and stoichiometry (n).

### Circular Dichlorism (CD)

The 10μM His-sumo-TOX protein was dialyzed in CD buffer (NaF, 100mM) and Na3PO4 (10mM) pH=8) overnight. Then, 50μL of the dialyzed protein was further diluted into 250μL 2μM with fresh un-used CD buffer. As a blank control, 50μL of the dialyzed buffer was also diluted by fresh CD buffer to ensure the salt condition were identical. Then, the 10μL of DMSO or 10mM DMSO stock solution of **KI-TOX-A3** was diluted to the fresh CD buffer and make a 10v/v% DMSO or 1mM **KI-TOX-A3** solution 10v/v% DMSO. Then, 2.5μL of this pre-diluted DMSO solution were added to the 247.5μL blank sample and TOX protein sample to make the negative and positive control. Then, 2.5μL of pre-diluted **KI-TOX-A3** solution was added to either the prepared blank or the TOX protein sample, so that there is blank sample that only has **KI-TOX-A3** without protein, and a protein **KI-TOX-A3** mixture. All the samples now have only 0.1%DMSO, which can prevent the high voltage and noise that could be caused by DMSO. Then, the samples ran through CD spectroscopy (JASCO J-1500) with screening rate of 20nM/minute and the 0.05nm reading data interval at the range of 260nm to 200nm. The runs repeated 6 times. The averaged readout data of the 6 runs were normalized by the blanks. Then, the protein-only group and protein-compound mixture group were plot on the Prism.

### Immunoprecipitation:

2mM solution of compound **KI-TOX-A3-0237b** was prepared in DMSO. The **KI-TOX-A3-0237b** solution and DMSO solvent were applied for NHS-active magnetic beads (Cytiva) coating with 10v/v% DIPEA and covered by aluminum foil cover. The beads were incubated overnight with head-to-end rotation in 4°C. Then the bead was immobilized on the magnetic rack. The supernatant of **KI-TOX0-A3-0237b** solution was collected and sent to LC-MS for quantification estimation. A 10% loss indicated 90% loaded onto the beads. The coated beads (100μL) were washed by water (1ml) for one time. Then, the coated and uncoated beads (300μL) were incubated in the 10mM ethanolamine TBS solution for de-activation on both groups. The incubation was overnight, and the beads were washed by water (1ml x 3 times) and restored in isopropanol under -20°C. On the day of pulldown, the beads are washed by 1X PBS for 4 times at 500μL and then blocked by Superblock® for 1 hour prior to the pulldown. In the meantime, a fresh Molt4 lysate was obtained from Molt4 cell lysis. The IP-lysis buffer (Thermofisher) was applied along with 1x protease inhibitor cocktail (Pierce) and 5μM MG-132 (Thermofisher) in the cell lysis. In the pulldown, we distributed 250μL of 2mg/ml lysate in each low-binding Eppendorf tube and incubated the lysate with the ethanolamine-deactivated uncoated beads (50μL). This step is to exclude the non-specific binding proteins of the lysate. The incubation was performed in 4°C for 1h. Then, the lysates were spitted to 4 groups, including (1) 'DMSO group' 50μL de-activated uncoated beads; (2) 'Pull down group' 50μL of de-activated **KI-TOX-A3-0237b** coated beads; (3) 'Comp. A3_20μM' 50μL of de-activated **KI-TOX-A3-0237b** coated beads and 20 μM **KI-TOX-A3; (4)** 'Comp. A3_40μM' 50μL of de-activated **KI-TOX-A3-0237b** coated beads and 40 μM **KI-TOX-A3** for 45 minutes in the 4°C with head-to-end rotation. The beads were then immobilized on the magnetic rack and washed by wash buffer 1 (500μL x3) and wash buffer 2 (500μL). Then, the beads were incubated with elution buffer (30μL per tube) in 95°C for 10 minutes. The eluted protein lysates were collected into aliquots (10μL/low-bind tube) and run through western blot.

Wash buffer 1: Octyl β-D-glucopyranoside (8mg/ml), Tris-HCl (50mM), MG 132 (5μM), protease inhibitor (1x); Wash buffer 2: TBS-T 1x, MG132 5μM, Protease inhibitor (1x); Elution buffer: LDS Nupage buffer 1x, DTT 10mM.

## IP-MS:

*Sample preparation*

The magnetic Dynabead protein G beads (300μL, 50μL /sample, 6 samples – 3 DMSO groups and 3 A3-10μM groups) were washed by 'wash and binding buffer' in the kit for 3 times (500μL/time), and resuspended in 620μL wash & binding buffer with 10μL anti-TOX antibody (68ng/μL) in the low-binding tube. Then the beads were incubated in 4°C for overnight. In the day 2, the Molt4 cell lysates were obtained in 2mg/ml with protease inhibitor and MG-132 addition. Then, 3 samples were added with compound **KI-TOX-A3** at 10μM (1% DMSO) and 3 samples with 1% DMSO along. In the meantime, the protein G beads were washed and added into the lysate. The incubation took overnight, and the beads were washed by the wash buffer that was provided in the kit for 3times (500μL/wash). The antibody and PPI proteins of TOX beads were eluted by the elution buffer (3 times, 30μL/elution). Then, the eluted samples were sent to Proteomic core of Koch institute for digestion and peptide analysis.

*Digestion*

Digestions were performed with S-trap micro spin columns from Protifi per manufacturer's protocol. Volumes in the protocol were adjusted for samples volume, 10 mM DTT(final concentration) was used instead of TCEP and 20 mM iodoacetamide (final concentration) was used instead of MMTS. After adding the DTT sample tubes were placed on a heating block for 10 minutes at 95C. Proteins were then alkylated with iodoacetamide, after adding the iodoacetamide samples incubated at RT for 30 minutes in the dark.

*LC-MS/MS*

The tryptic peptides were separated by reverse phase HPLC (Thermo Ultimate 3000) using a Thermo PepMap RSLC C18 column(2um tip, 75umx50cm PN# ES903) over a 90 minute gradient before nano electrospray using a Orbitrap Exploris 480 mass spectrometer (Thermo). Solvent A was 0.1% formic acid in water and solvent B was 0.1% formic acid in acetonitrile. The gradient conditions were 1% B (0-10 min at 300nL/min) 1% B (10-15 min, 300 nL/min to 200 nL/min) 1-7% B (15-20 min, 200nL/min), 7-25% B (20-54.8 min, 200nL/min), 25-36 B (54.8-65 min, 200nL/min), 36-80% B (65-65.5 min, 200 nL/min), 80% B (65.5-70 min, 200nL/min), 80-1% B (70-70.1 min, 200nL/min), 1% B (70.1-90 min, 200nL/min).

The Thermo Orbitrap Exploris 480 mass spectrometer was operated in a data-dependent mode. The parameters for the full scan MS were: resolution of 120,000 across 375-1600 m/z and maximum IT 25 ms. The full MS scan was followed by MS/MS for as many precursor ions in a two second cycle with a NCE of 28, dynamic exclusion of 20 s and resolution of 30,000.

*Database Search*

Raw mass spectral data files (.raw) were searched using Sequest HT in Proteome Discoverer (Thermo) against a Human database(Uniprot) and a contaminates database(made in house) with the following search parameters: 10 ppm mass tolerance for precursor ions; 0.02 Da for fragment ion mass tolerance; 2 missed cleavages of trypsin; fixed modification were carbamidomethylation of cysteine, variable modifications were methionine oxidation, methionine loss at the N-terminus of the protein, acetylation of the N-terminus of the protein and also Met-loss plus acetylation of the protein N-terminus.

### EMSA:

TOX protein was aliquoted to10μL with multiple dilutions (12, 6, 3, 1.5, 0.75, 0.375, 0.1875, 0μM) per tube and stored on ice under the EMSA binding buffer (Tris 20mM, NaCl 150mM, 2mM MgCl2, pH=8). Then the DNA oligonucleotide was prepared and dissolved in DNA folding buffer (TRIS 10mM, NaCl 50mM, 1mM MgCl2, 1mM EDTA) to make 10μM. The DNA was self-annealed by heated to 97°C for 5 minutes and cooled to 25°C with a ramp of 0.1°C/sec. Then, the DNA stock solution was diluted to 200nM by the folding buffer. Then, the protein was mixed with the DNA and make 20 μL mixture. The binding took 30 minutes before we run the gel. For the competition EMSA, the compounds were dissolved into the protein and reached to 0.5%DMSO solution. The compounds were incubated in the protein for 30 minutes. The nucleotide (10μL) was mixed with 10μL protein-compound mixture. The final protein concentration is 3 μM, the DNA is 100nM, and the compounds are from 10 μM to 0.31 μM with 2-fold dilution. While the incubation was ongoing, we pre-ran the TBE gel with 0.5x TBE buffer (150V, 30minutes). When the incubation was finished, the samples were mixed with 6x loading dye and loaded to the TBE gel. While running the gel, the running temperature should be 4°C and the voltage=110V and running time extended to 65 minutes. Note that higher temperature and voltage could interrupt DNA-protein interaction.

### Western blot:

Western blot analysis was conducted as follows: Various concentrations of **KI-TOX-A3** in DMSO were added to the Jurkat, Molt4, HBP-ALL and CD8$^{+}$ T cells, ensuring a final DMSO level of 1%. After various hours of treatment, cells were washed with cold PBS and lysed with RIPA buffer containing phosphatase and protease inhibitors. The lysates were then centrifuged, and the supernatants were collected. Total protein concentration was determined using a BCA protein assay kit, and lysates were diluted accordingly to achieve equal protein concentration. Subsequently, 20-40 µg of each lysate was loaded onto TGX MIDI 4-20% gels and run at 150V for 65 minutes. The gel was transferred onto Turbo PDVF membranes, blocked with 5% BSA, and then incubated overnight with specific antibodies. The following day, membranes were washed with TBST, incubated with secondary antibodies, and bands were quantified using an Odyssey CLx Image system.

### qPCR:

The cells were collected, centrifuged (500 xg, 5 minutes) and washed by 1x PBS prior to the lysis. We collected the RNAs using the Invitrogen RNA extraction kit (PureLink RNA Mini kit, Thermofisher). The extraction protocol followed the manufacture procedure. The RNA levels were checked with the Nanodrop detection and normalized with RNAse free water. After RNA extraction, we applied cDNA reverse transcription directly with the Applied Biosystem High-capacity reverse transcription kit (Thermofisher). To prepare a cDNA master mix, we followed the manufacture recipe. Later, we mixed 10μL of master mix solution with 10 μL of extracted RNA. After brief pipetting, the mixture was run with the thermocycle that was suggested by the cDNA kit. After 2hrs of incubation, the cDNAs were prepared and is diluted to 200μL (10x dilution), which is the working concentration for qPCR.

To run the qPCR, we pre-mixed the 1μL/well primers, 3μL/well water, with 5μL/well 2x SYBR master mix to make a working solution with 8μL/ well volume. Then, the working solutions were dispensed into wells that are triplicated. Later, the cDNA samples were dispensed into the wells with 2μL/well. After pipetting the plate was centrifuge with sealing sheet. Then, the 384-plate was placed on the CFX-384 for thermocycles.

The data obtained through the CFX Manager™ software (Bio-Rad) was exported to Microsoft® Excel® for analysis. The Ct (cycle threshold) value for each fluorescence channel in both uninduced and induced control wells was determined as the mean Ct value across all corresponding control wells ($n \geq 10$). ΔCt was calculated as the disparity between the Ct value in the Cy5 channel (PSA) and the FAM channel (GAPDH), while ΔΔCt was computed as the difference between the ΔCt value of the assay well and that of the induced wells to standardize expression relative to the induced wells. The expression fold change for each well was defined as 2ˆ(-ΔΔCt). The average expression fold change for qPCR technical replicates ($n = 3$) was utilized to assess compound performance, with each target being experimentally replicated once.

### Cell Cytotoxicity:

Cells were seeded to the regular flat transparent 96-well plate with density around 8000 to 10,000 cells per 100μL in each well. The cells were treated with multiple concentrations of compounds, alone with DMSO control that the DMSO was limited to 1% in media. The cells were treated for 24h, 48h, 72h, and 96h. The cells proliferation was tested by CellTiterGlo (Promega) by adding 80μL of the CTG buffer with the cells and read through the luminescence in Tecan 200.

### Cell cycle analysis:

Cells were treated with **KI-TOX-A3** at 20, 10, and 5μM in Jurkat and T cells for 12h. The cells were then washed by 1x cold PBS and fixed with 3.7% paraformaldehyde in PBS for 15 minutes at room temperature. The cells were then treated with 0.2mg/ml RNAse I for 0.5hr at room temperature. Then, the cells were treated with Propidium Iodide (PI) staining at 0.1mg/ml concentration for another 1hr. The cells were then sent to flow cytometry to analyze the cell cycle.

### T-cell exhaustion and treatments

$CD8^+$ T cells were stimulated in day 1 with 20μL/1M cells of the ImmunoCult™ Human CD3/CD28/CD2 T Cell Activator (STEMCELL) for 2 days. At day 3, the media of the cells were changed by centrifuge and the cells were split to 1M/ml density with fresh media again. The cells were stimulated again with 20μL stimulator for another 2 days and media was changed again with another 20μL stimulators. This is called one cycle. The cells were grown in another 4 cycles to reach exhaustion. At the end of the $5^{th}$ cycle, the cells were treated by 20μM, 10 μM, and 5 μM of **KI-TOX-A3**, **KI-TOX-D22**, and **KI-TOX-P14** with 1% DMSO buffer solution. The cells were collected by dispensed 100μL in a 96-well V shape plate (Biomarker detection), 100μL in a U shape plate (future stimulation and detection of cytokine), 400μL in Eppendorf tubes (for Western blot), and 400μL for another sets of Eppendorf tube (for PCR) in after 24h and 48h treatment.

In the V-shape plate, the cells were centrifuge with 500xg 5 minutes, removed media and washed by 200μL FACS buffer, re-centrifuged with 1000 xg for another 5 minutes. The cells were ready for biomarker staining. The biomarker antibody master mix contains 1:100 dilution of antibodies: PE PD-1, APC Tim-3 and PE-Cy7 LAG-3 (Biolegend) in FACS buffer. The cells were merged with 50μL of the master mix solution for 15 minutes in refrigerator (4-8°C) and centrifuged by 1000 xg for 5 minutes. The staining buffers were removed, and the cells were washed once with 200μL FACS buffer and re-suspended with 110μL FACS for the Symphony HTS A3 flow cytometer to detect.

In the U shape plate, the cells were further treated with 1μL of 100x cell activation cocktail with brefeldin A (an ionomycin/PMA/ Brefeldin A mixture from Biolegend) to make a final concentration of 1x cocktail in 1% DMSO solution. The cells were stimulated for 4 more hours after collection. This step is to test the intracellular cytokine level due to the addition of Brefeldin A. The cells were transferred into a V shape plate, washed by PBS (100μL/well) and fixed by the fixation buffer (100μL/well, Biolegend) for 15 minutes at room temperature. The cells were further washed by permeability buffer (Biolegend) for two more times before staining. The antibody Alexa 591 TOX and PE-cy7 IFN- γ (Biolegend) were diluted into the permeable buffer with 1:100 dilution. The master mix (50μL/well) was mixed well with the cells and the staining could go as short as 30 minutes to overnight at 4°C. The cells were washed with FACS buffer and ran the flow cytometry in Symphony A3 HTS machine.

For the batch of PCR, the RNA preparation protocol followed the previous Jurkat mRNA study. For the Western blot, the protocol shared with the cancer cell procedure above.

For the co-culture killing assay, both $T_{ex}$ and non-exhausted T cells were incubated in the 96-well plate with 100μL/well and the cell density was 0.5million/ml. The cells were treated by 1% DMSO solution of TOX inhibitors at various concentrations (2.5, 5, and 10μM) for 24h. Then, the cells were washed by 1x PBS and resuspended into a fresh T cell media. The CD19-Ramos cells were added into the T cells at a 3:1 E:T ratio (30k T cells, 10k Ramos) along with aCD19 BiTEs (1μg/mL) and cocultured in 200 μL in a 96-well plate. After 12h treatment in 37°C (mix by pipetting after 6h), we tested the supernatants of each well with LDH assay (CytoTox 96, Promega). As the positive control, we lysed Ramos cells according to the manufacturer protocol. The supernatant of the Ramos-only well was used as a negative control.

### TOX-KAT7 functional assay

Titration:

We prepared a reaction buffer with 50 mM Tris-HCl pH 8.0, 150mM NaCl, and 2 mM $MgCl_2$. On one hand, 0 or 0.2 μg of Histone H3 was dissolved to 5 μl reaction buffer that contains 40 μM acetyl-CoA. The KAT7 will be dissolved with or without TOX protein and top up to final 15 μl reaction buffer. The KAT7 or TOX-KAT7 mixtures will be incubated for 30 mins, then this mixture will be added to the H3-acetyl coA mixture for 2 hr at room temperature to make 20 μl reaction buffer where the acetyl coA concentration is 20μM. The groups were denatured by BME contained Laemmli buffer with 95°C for 8 minutes. The denatured sample were ready for Western blot study.

TOX inhibition:

For TOXi inhibition, we first dissolved KI-TOX-A3 into 10mM DMSO solution and then we had 10x dilution and make 1mM DMSO solution. Then, we had 2-fold dilutions to 0.5, 0.25, 0.125, and

0.0625mM DMSO solution. Then, we had another 10x dilution of these dilutions by adding 10uL of them into the 90uL binding buffer and make 10% DMSO solutions. Then, to make 1% DMSO solutions to TOX, we added 1µL of the 10% DMSO solution of KI-TOX-A3 to 4uL TOX protein (0.7mg/ml), followed by adding 5uL of KAT7 (0.15ug/5uL). We incubated this mixture for 10 minutes, and then we added 9uL of acetyl CoA-contained H3 solution (same preparation as the titration solution) and another 1uL of the 10% DMSO solution of the KI-TOX-A3 to make 20uL sample per group. Every group has 1%DMSO solution of varies concentrations of KI-TOX-A3. After 1-hr incubation, the 20 µl of reaction products will be denatured by mixing with 20 µL Laemmli buffer (BME added) by heating to 95°C for 8 mins. 20 µl of the denatured buffer were run on a 12.5% SDS-PAGE and stained by KAT7, H3 and H3K14ac antibodies.

### Chemical characterization

NMR: The Dimethyl sulfoxide-d6 deuterated solvent was used. The chemicals ran through the Neo500, a three-channel Bruker Avance Neo spectrometer operating at 500.34 MHz.
Column purification: Chemicals were further purified through the Combi-flash system.
HPLC purification: Interchim HPLC Puriflash system with c18 reverse phase column. The buffer A is 0.1% formic acid in water, and buffer B is 0.1% formic acid in ACN. The method is: 2minutes of 10% B and 90% A, then 15 minutes gradient ramp to 90% B and then equilibrate to 10% B in 2 minutes. Last, the column was equilibrated to 100% B for another 2 minutes.
LC-MS: Agilent 6125B mass spectrometer attached to an Agilent 1260. A gradient method that ramps from 10% ACN to 100%ACN over the period of six minutes and then equilibrates 10% ACN for one minute. The column is a c18 reverse phase column.
QTOF HRMS: Agilent 6545 mass spectrometer coupled to an Agilent Infinity 1260 LC system. The buffer A is 0.1% formic acid in water, and buffer B is 0.1% formic acid in ACN. The column is a c18 reverse phase column.

### Organic synthesis:

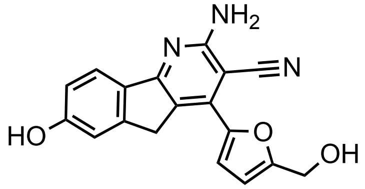


**KI-TOX-A3**

5-(hydroxymethyl)furan-2-carbaldehyde (151 mg, 1.2 equiv) was combined with 5-hydroxy-2,3-dihydro-1H-inden-1-one (148 mg, 1 equiv), ammonium acetate (350 mg, 4.5 equiv), and malonitrile (66 mg, 1 equiv). The mixture was dissolved in Toluene and ethanol (1ml/4ml) and heated to 90-100 ºC overnight. The reaction solution was dried via rotary evaporation, and the crude compound was redissolved in a DCM:methanol (1:1) solution. The solution was stirred with 0.5 g of Amberlyst 15 strongly acidic cation exchange resin. The resin was washed three times with a DCM:methanol (1:1) solution. The compound was eluted with 2M ammonia in methanol (30 mL), and the filtrate was dried. The filtrate was purified by column chromatography with DCM:Methanol (10:1) gradient as the eluent. The product was obtained as a yellow powder (yield: 63 mg, 0.0197 mmol, 19.7%). $^{1}$H NMR (500 MHz, DMSO) δ 7.68 (d, J = 8.3 Hz, 1H), 7.36 (d, J = 3.5 Hz, 1H), 7.03 (s, 1H), 6.88 (d, J = 8.3 Hz, 1H), 6.61 (d, J = 3.5 Hz, 1H), 4.56 (s, 2H), 4.31 (s, 1H), 3.99 (s, 2H). $^{13}$C NMR (126 MHz, DMSO) δ 160.4, 158.3, 148.1, 146.2, 125.8, 123.0, 120.0, 119.1, 115.6, 115.1, 112.3, 109.8, 108.0, 107.1, 79.2, 56.3, 46.1, 29.5. HPLC retention time: 3.8 minutes. Chemical Formula: $C_{18}H_{13}N_3O_3$. Exact Mass: 319.10. HRMS found: $[M+H]^+$ = 320.1034 m/z.

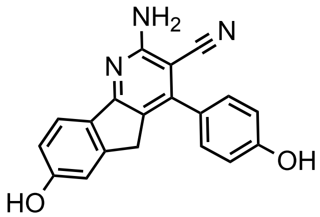

**KI-TOX-A3-0186**

4-hydroxybenzaldehyde (146 mg, 1.2 equiv) was combined with 5-hydroxy-2,3-dihydro-1H-inden-1-one (148 mg, 1 equiv), ammonium acetate (350 mg, 4.5 equiv), and malonitrile (66 mg, 1 equiv). The mixture was dissolved in ethanol:toluene (1:1, 5 mL) and heated to 90-100 ºC overnight. The reaction solution was dried via rotary evaporation, and the crude compound was redissolved in a DCM:methanol (1:1) solution. The solution was stirred with 0.5 g of Amberlyst 15 strongly acidic cation exchange resin. The resin was washed three times with a DCM:methanol (1:1) solution. The compound was eluted with 2M ammonia in methanol (30 mL), and the filtrate was dried. The filtrate was purified by column chromatography with DCM:Methanol (10:1) gradient as the eluent. Yellow powder with yield of 203mg (0.64 mmol, 64%). $^{1}$H NMR (500 MHz, DMSO) δ 7.68 (d, *J* = 8.3 Hz, 1H), 7.45 (d, *J* = 8.6 Hz, 2H), 6.96 (d, *J* = 1.8 Hz, 2H), 6.92 (d, *J* = 8.6 Hz, 2H), 6.87 (dd, *J* = 8.4, 2.2 Hz, 1H), 6.69 (s, 2H), 3.64 (s, 2H). $^{13}$C NMR (126 MHz, DMSO) δ 162.9, 162.3, 160.1, 158.7, 149.7, 149.0, 131.4, 130.6, 126.7, 123.0, 122.9, 118.7, 115.8, 115.6, 112.5, 84.9, 33.6. HPLC retention time: 3.8 minutes. Chemical Formula: C19H13N3O2. Exact Mass: 315.10. HRMS found $[M+H]^+$ = 316.1090 m/z.

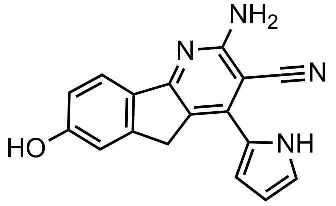

**KI-TOX-A3-0185**

1H-pyrrole-2-carbaldehyde (114 mg, 1.2 equiv) was combined with 5-hydroxy-2,3-dihydro-1H-inden-1-one (148 mg, 1 equiv), ammonium acetate (350 mg, 4.5 equiv), and malonitrile (66 mg, 1 equiv). The mixture was dissolved in ethanol:toluene (1:1, 5 mL) and heated to 90-100 ºC overnight. The reaction solution was dried via rotary evaporation, and the crude compound was redissolved in a DCM:methanol (1:1) solution. The solution was stirred with 0.5 g of Amberlyst 15 strongly acidic cation exchange resin. The resin was washed three times with a DCM:methanol (1:1) solution. The compound was eluted with 2M ammonia in methanol (30 mL), and the filtrate was dried. The filtrate was purified by column chromatography with DCM:Methanol (10:1) gradient as the eluent. Brown powder with yield of 23 mg (0.08 mmol, 8%). $^{1}$H NMR (500 MHz, DMSO) δ 7.66 (d, J = 8.3 Hz, 1H), 7.08 (q, J = 2.6 Hz, 2H), 7.00 (d, J = 1.7 Hz, 1H), 6.87 (dd, J = 8.3, 2.1 Hz, 1H), 6.74 (t, J = 3.6 Hz, 2H), 6.65 (s, 2H), 6.30 – 6.27 (m, 1H), 3.87 (s, 2H). $^{13}$C NMR (126 MHz, DMSO) δ 163.1, 162.5, 160.1, 148.7, 140.7, 131.4, 125.9, 122.9, 122.1, 121.0, 119.3, 115.5, 112.5, 112.3, 109.7, 82.4, 34.2. HPLC retention time: 3.9 minutes. Chemical Formula: $C_{17}H_{12}N_4O$. Exact Mass: 288.10. HRMS: $[M+H]^+$ = 289.1087 m/z.

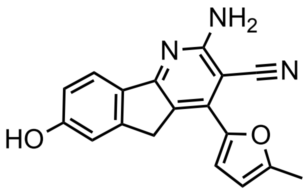


**KI-TOX-A3-0181**

5-methylfuran-2-carbaldehyde (132 mg, 1.2 equiv) was combined with 5-hydroxy-2,3-dihydro-1H-inden-1-one (148 mg, 1 equiv), ammonium acetate (350 mg, 4.5 equiv), and malonitrile (66 mg, 1 equiv). The mixture was dissolved in ethanol:toluene (1:1, 5 mL) and heated to 90-100 ºC overnight. The reaction solution was dried via rotary evaporation, and the crude compound was redissolved in a DCM:methanol (1:1) solution. The solution was stirred with 0.5 g of Amberlyst 15 strongly acidic cation exchange resin. The resin was washed three times with a DCM:methanol (1:1) solution. The compound was eluted with 2M ammonia in methanol (30 mL), and the filtrate was dried. The filtrate was purified by column chromatography with DCM:Methanol (10:1) gradient as the eluent. Yellow powder with yield of 52 mg (0.172 mmol, 17.2%). $^{1}$H NMR (500 MHz, DMSO) δ 10.02 (s, 1H), 7.67 (d, $J$ = 8.3 Hz, 1H), 7.32 (d, $J$ = 3.4 Hz, 1H), 6.88 (dd, $J$ = 8.3, 2.1 Hz, 1H), 6.73 (s, 2H), 6.46 – 6.41 (m, 1H), 3.96 (s, 2H), 2.45 (s, 3H). $^{13}$C NMR (126 MHz, DMSO) δ 163.8, 162.6, 160.4, 154.9, 148.9, 147.4, 135.8, 131.1, 122.9, 119.6, 119.1, 115.7, 115.6, 112.3, 109.3, 35.4, 14.0. HPLC retention time: 4.3 minutes. Chemical Formula: $C_{18}H_{13}N_3O_2$. Exact Mass: 303.10. HRMS found: $[M+H]^+$ = 304.1085 m/z.

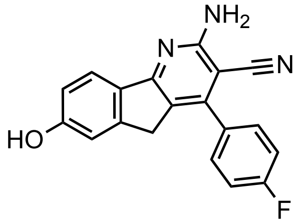


**KI-TOX-A3-0180**

4-fluorobenzaldehyde (148 mg, 1.2 equiv) was combined with 5-hydroxy-2,3-dihydro-1H-inden-1-one (148 mg, 1 equiv), ammonium acetate (350 mg, 4.5 equiv), and malonitrile (66 mg, 1 equiv). The mixture was dissolved in ethanol:toluene (1:1, 5 mL) and heated to 90-100 ºC overnight. The reaction solution was dried via rotary evaporation, and the crude compound was redissolved in a DCM:methanol (1:1) solution. The solution was stirred with 0.5 g of Amberlyst 15 strongly acidic cation exchange resin. The resin was washed three times with a DCM:methanol (1:1) solution. The compound was eluted with 2M ammonia in methanol (30 mL), and the filtrate was dried. The filtrate was purified by column chromatography with DCM:Methanol (10:1) gradient as the eluent. Yellow powder with yield of 95 mg (0.3 mmol, 30%) $^{1}$H NMR (500 MHz, DMSO) δ 7.72 – 7.64 (m, 2H), 7.44 – 7.35 (m, 2H), 6.97 (d, $J$ = 1.8 Hz, 1H), 6.92 – 6.83 (m, 1H), 6.81 (s, 1H), 5.76 (s, 0H), 3.62 (s, 2H). $^{13}$C NMR (126 MHz, DMSO) δ 163.9, 163.2, 162.2, 161.9, 160.3, 149.0, 148.5, 132.7, 131.3, 123.1, 118.3, 116.2, 115.7, 112.5, 84.8, 33.3. HPLC retention time: 4.4minutes. Chemical Formula: $C_{19}H_{12}FN_3O$. Exact Mass: 317.10. HRMS found: $[M+H]^+$ = 318.1043 m/z.

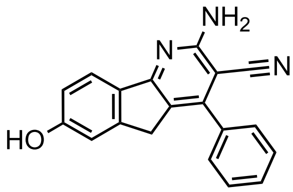

**KI-TOX-A3-0184**

Benzaldehyde (127 mg, 1.2 equiv) was combined with 5-hydroxy-2,3-dihydro-1H-inden-1-one (148 mg, 1 equiv), ammonium acetate (350 mg, 4.5 equiv), and malonitrile (66 mg, 1 equiv). The mixture was dissolved in ethanol:toluene (1:1, 5 mL) and heated to 90-100 ºC overnight. The reaction solution was dried via rotary evaporation, and the crude compound was redissolved in a DCM:methanol (1:1) solution. The solution was stirred with 0.5 g of Amberlyst 15 strongly acidic cation exchange resin. The resin was washed three times with a DCM:methanol (1:1) solution. The compound was eluted with 2M ammonia in methanol (30 mL), and the filtrate was dried. The filtrate was purified by column chromatography with DCM:Methanol (10:1) gradient as the eluent. Yellow powder with yield of 155 mg (0.52 mmol, 52%) $^{1}$H NMR (500 MHz, DMSO) δ 7.70 (d, J = 8.3 Hz, 1H), 7.63 – 7.49 (m, 7H), 6.96 (d, J = 1.8 Hz, 2H), 6.88 (dd, J = 8.3, 2.2 Hz, 1H), 6.79 (s, 2H), 3.62 (s, 2H), 2.09 (s, 2H).$^{13}$C NMR (126 MHz, DMSO) δ 163.2, 162.2, 160.2, 149.5, 149.0, 136.3, 131.3, 129.5, 129.5, 129.1, 129.0, 128.9, 123.1, 123.0, 118.3, 115.7, 112.5, 84.8, 33.4.HPLC retention time: 4.3minutes. Chemical Formula: $C_{19}H_{13}N_{3}O$. Exact Mass: 299.11. HRMS found: $[M+H]^{+}$ =300.1130 m/z.

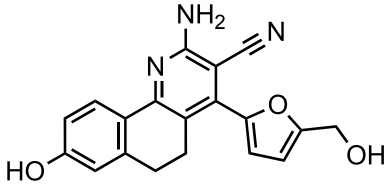

**KI-TOX-A3-0176**

5-(hydroxymethyl)furan-2-carbaldehyde (151 mg, 1.2 equiv) was combined with 6-hydroxy-3,4-dihydronaphthalen-1(2H)-one (176 mg, 1 equiv), ammonium acetate (350 mg, 4.5 equiv), and malonitrile (66 mg, 1 equiv). The mixture was dissolved in ethanol:toluene (1:1, 5 mL) and heated to 90-100 ºC overnight. The reaction solution was dried via rotary evaporation, and the crude compound was redissolved in a DCM:methanol (1:1) solution. The solution was stirred with 0.5 g of Amberlyst 15 strongly acidic cation exchange resin. The resin was washed three times with a DCM:methanol (1:1) solution. The compound was eluted with 2M ammonia in methanol (30 mL), and the filtrate was dried. The filtrate was purified by column chromatography with DCM:Methanol (10:1) gradient as the eluent. Dark yellow powder with yield of 122 mg (0.372 mmol, 37.2%) $^{1}$H NMR (500 MHz, DMSO) δ 9.92 (s, 1H), 8.00 (d, *J* = 8.5 Hz, 1H), 6.84 (d, *J* = 3.3 Hz, 1H), 6.75 (dd, *J* = 8.5, 2.4 Hz, 1H), 6.68 – 6.63 (m, 4H), 6.54 (d, *J* = 3.3 Hz, 1H), 5.38 (t, *J* = 5.8 Hz, 1H), 4.49 (d, *J* = 5.6 Hz, 2H), 2.81 – 2.66 (m, 6H). $^{13}$C NMR (126 MHz, DMSO) δ 160.1, 159.8, 157.5, 155.6, 147.1, 141.7, 140.5, 128.2, 125.2, 117.8, 117.3, 114.6, 114.6, 114.5, 109.1, 85.3, 56.2, 28.0, 24.6. HPLC retention time: 4.0 minutes. Chemical Formula: $C_{19}H_{15}N_{3}O_{3}$. Exact Mass: 333.11. HRMS found: $[M+H]^{+}$ = 334.1192 m/z.

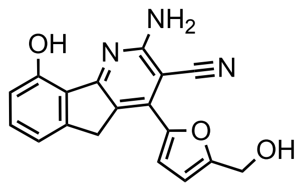

**KI-TOX-A3-0177**

5-(hydroxymethyl)furan-2-carbaldehyde (151 mg, 1.2 equiv) was combined with 7-hydroxy-2,3-dihydro-1H-inden-1-one (148 mg, 1 equiv), ammonium acetate (350 mg, 4.5 equiv), and malonitrile (66 mg, 1 equiv). The mixture was dissolved in ethanol and heated to 90-100 ºC overnight. The reaction solution was dried via rotary evaporation, and the crude compound was redissolved in a DCM:methanol (1:1) solution. The solution was stirred with 0.5 g of Amberlyst 15 strongly acidic cation exchange resin. The resin was washed three times with a DCM:methanol (1:1) solution. The compound was eluted with 2M ammonia in methanol (30 mL), and the filtrate was dried. The filtrate was purified by column chromatography with DCM:Methanol (10:1) gradient as the eluent. Yellow powder with yield of 33 mg (0.103 mmol, 10.3%) $^{1}$H NMR (500 MHz, DMSO) δ 7.44 (d, $J$ = 3.5 Hz, 1H), 7.38 (t, $J$ = 7.8 Hz, 1H), 7.15 (d, $J$ = 7.4 Hz, 1H), 7.08 (s, 2H), 6.86 (d, $J$ = 8.1 Hz, 1H), 6.64 (d, $J$ = 3.5 Hz, 1H), 5.49 (s, 1H), 4.57 (s, 2H), 4.10 (s, 2H), 3.17 (s, 1H). $^{13}$C NMR (126 MHz, DMSO) δ 164.0, 161.9, 158.8, 155.0, 147.8, 147.3, 136.5, 132.4, 124.3, 119.2, 118.6, 116.9, 115.7, 113.7, 109.9, 79.6, 56.3, 36.0. HPLC retention time: 4.4 minutes. Chemical Formula: $C_{18}H_{13}N_3O_3$. Exact Mass: 319.10. HRMS found: $[M+H]^+$ = 320.1029 m/z.

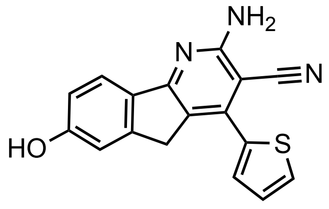

**KI-TOX-A3-0182**

Thiophene-2-carbaldehyde (151 mg, 1.2 equiv) was combined with 5-hydroxy-2,3-dihydro-1H-inden-1-one (148 mg, 1 equiv), ammonium acetate (350 mg, 4.5 equiv), and malonitrile (66 mg, 1 equiv). The mixture was dissolved in ethanol:toluene (1:1, 5 mL) and heated to 90-100 ºC overnight. The reaction solution was dried via rotary evaporation, and the crude compound was redissolved in a DCM:methanol (1:1) solution. The solution was stirred with 0.5 g of Amberlyst 15 strongly acidic cation exchange resin. The resin was washed three times with a DCM:methanol (1:1) solution. The compound was eluted with 2M ammonia in methanol (30 mL), and the filtrate was dried. The filtrate was purified by column chromatography with DCM:Methanol (10:1) gradient as the eluent. Yellow powder with yield of 15 mg (0.049 mmol, 4.9%). $^{1}$H NMR (500 MHz, DMSO) δ 7.44 (d, $J$ = 3.5 Hz, 1H), 7.38 (t, $J$ = 7.8 Hz, 1H), 7.15 (d, $J$ = 7.4 Hz, 1H), 7.08 (s, 2H), 6.86 (d, $J$ = 8.1 Hz, 1H), 6.64 (d, $J$ = 3.5 Hz, 1H), 5.49 (s, 1H), 4.57 (s, 2H), 4.10 (s, 2H), 3.17 (s, 1H). $^{13}$C NMR (126 MHz, DMSO) δ 164.0, 161.9, 158.8, 155.0, 147.8, 147.3, 136.5, 132.4, 124.3, 119.2, 118.6, 116.9, 115.7, 113.7, 109.9, 79.6, 56.3, 36.0. HPLC retention time: 4.3 minutes. Chemical Formula: $C_{17}H_{11}N_3OS$. Exact Mass: 305.06. HRMS: $[M+H]^+$ = 306.0697 m/z.

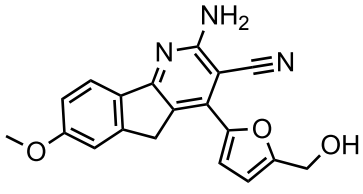

**KI-TOX-A3-0174**

5-(hydroxymethyl)furan-2-carbaldehyde (151 mg, 1.2 equiv) was combined with 5-Methoxy-2,3-dihydro-1H-inden-1-one (162 mg, 1 equiv), ammonium acetate (350 mg, 4.5 equiv), and malonitrile (66 mg, 1 equiv). The mixture was dissolved in ethanol:toluene (1:1, 5 mL) and heated to 90-100 ºC overnight. The reaction solution was dried via rotary evaporation, and the crude compound was redissolved in a DCM:methanol (1:1) solution. The solution was stirred with 0.5 g of Amberlyst 15 strongly acidic cation exchange resin. The resin was washed three times with a DCM:methanol (1:1)solution. The compound was eluted with 2M ammonia in methanol (30 mL), and the filtrate was dried. The filtrate was purified by column chromatography with DCM:Methanol (10:1) gradient as the eluent. Yellow powder with yield of 25 mg (0.075 mmol, 7.5%) $^{1}H$ NMR (500 MHz, DMSO) δ 7.77 (d, *J* = 8.4 Hz, 1H), 7.39 (d, *J* = 3.5 Hz, 1H), 7.26 (d, *J* = 2.0 Hz, 1H), 7.06 (dd, *J* = 8.5, 2.3 Hz, 2H), 6.83 (s, 2H), 6.63 (d, *J* = 3.5 Hz, 1H), 4.57 (s, 2H), 4.06 (s, 2H), 3.86 (s, 2H). $^{13}C$ NMR (126 MHz, DMSO) δ 163.5, 162.6, 161.9, 158.4, 148.9, 148.0, 136.1, 132.5, 122.8, 120.2, 115.2, 114.7, 110.6, 109.8, 56.3, 56.0, 35.7. HPLC retention time: 4.2 minutes. Chemical Formula: $C_{19}H_{15}N_3O_3$. Exact Mass: 333.11. HRMS: $[M+H]^+$ = 334.1194 m/z.

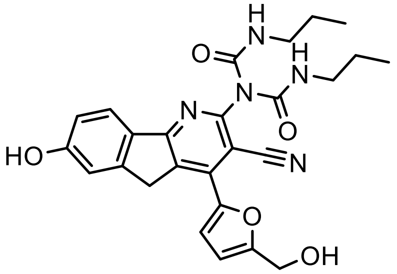

**KI-TOX-A3-0189**

Compound KI-TOX-A3 (320 mg, 1 mmol, 1 equiv) was first protected by Tert-Butyldimethylsilyl chloride (1.2 g, 8 mmol, 8 equiv) with DMF (5 mL) and imidazole (544 mg, 8 mmol, 8 equiv). The reaction was finished after 6 hr stirring according to the TLC. The solution was purified by Ethyl acetate:Hexane=3:7 in column chromatography and the protected intermediate (500 mg, 0.91 mmol, 91%) was a pale yellow solid. This intermediate (55 mg, 0.1 mmol) was then dissolved into DMF (1 mL) and mixed with isocyanopropane (20 mg, 0.23 mmol, 2.3 equiv) and followed with piperidine addition (20 mg, 0.23 mmol, 2.3 equiv). The reaction turn dark yellow overnight and was quenched by adding sodium bicarbonate solution, followed by DCM extraction. The organic layer was collected and dried by sodium sulfate. The compound was purified with chromatography by gradient DCM:Methanol solution. The final product was a yellow paste (8.3 mg, 0.0169 mmol, 16.9%). $^{1}H$ NMR (500 MHz, DMSO) δ 8.18 (s, 1H), 7.92 – 7.57 (m, 2H), 7.07 – 6.98 (m, 2H), 6.70 (t, J = 5.5 Hz, 2H), 6.63 (d, J = 3.5 Hz, 1H), 5.46 (t, J = 5.7 Hz, 2H), 5.09 (d, J = 5.8 Hz, 2H), 4.53 (d, J = 5.1 Hz, 2H), 3.04 (q, J = 6.4 Hz, 3H), 1.54 (dq, J = 14.9, 7.4 Hz, 4H), 1.39 (dh, J = 21.5, 7.3 Hz, 4H), 1.24 (s, 2H), 0.82 (q, J = 7.3 Hz, 6H). 13C NMR (126 MHz, DMSO) δ 177.9, 170.1, 162.8, 151.4,

146.8, 139.3, 122.2, 119.9, 109.9, 94.1, 83.8, 72.1, 62.9, 56.4, 48.3, 42.4, 23.6, 22.0, 11.7. HPLC retention time: 4.5 minutes. Chemical Formula: $C_{26}H_{27}N_5O_5$. Exact Mass: 489.20. HRMS: $[M+H]^+$ = 490.2095 m/z.

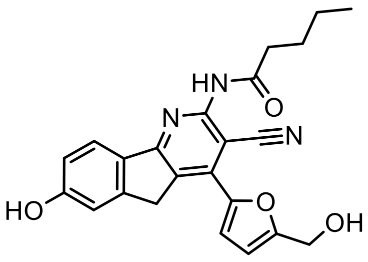


**KI-TOX-A3-0190**

Compound **KI-TOX-A3** (32 mg, 0.1 mmol, 1 equiv) dissolved into DMF (1 mL) and HATU (76 mg, 0.2 mmol, 2 equiv) and stirred. Then pentanoic acid (20 mg, 0.19 mmol, 1.9 equiv) was mixed with Hunig's base (0.1 mL) and DMF (0.5 mL) and stirred in another flask for 5 minutes. Then, the mixture was added dropwise to the **KI-TOX-A3** mixture, and the solution turned bright yellow. The reaction finished after overnight stirring, and the solution was washed by sodium bicarbonate (50 mL) and extracted by DCM (30 mL*2). The organic layer was collected and dried by sodium sulfate. The compound was purified with chromatography by gradient DCM:Methanol solution. The final product was a dark yellow solid (10 mg, 0.024 mmol, 24%). $^1$H NMR (500 MHz, DMSO) δ 10.70 (s, 1H), 7.79 (d, $J$ = 8.3 Hz, 1H), 7.49 (d, $J$ = 3.5 Hz, 1H), 7.10 (s, 1H), 6.94 (dd, $J$ = 8.4, 2.1 Hz, 2H), 6.67 (d, $J$ = 3.5 Hz, 1H), 5.52 (t, $J$ = 6.0 Hz, 1H), 4.59 (d, $J$ = 5.9 Hz, 2H), 4.20 (s, 1H), 1.63 (dt, $J$ = 15.0, 7.4 Hz, 4H), 1.40 (dq, $J$ = 14.6, 7.4 Hz, 4H), 0.92 (t, $J$ = 7.4 Hz, 2H). $^{13}$C NMR (126 MHz, DMSO) δ 172.3, 163.5, 160.9, 159.2, 154.8, 148.9, 136.0, 130.3, 128.5, 123.3, 116.3, 116.1, 112.4, 110.1, 56.3, 35.6, 27.5, 22.2, 14.3. HPLC retention time: 4.1 minutes. MS: $[M+H]^+$ = 404.1615 m/z.

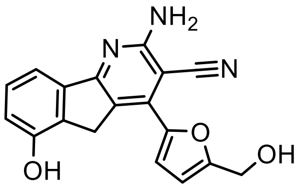


**KI-TOX-A3-0179**

5-(hydroxymethyl)furan-2-carbaldehyde (151 mg, 1.2 equiv) was combined with 4-hydroxy-2,3-dihydro-1H-inden-1-one (148 mg, 1 equiv), ammonium acetate (350 mg, 4.5 equiv), and malonitrile (66 mg, 1 equiv). The mixture was dissolved in ethanol and heated to 90-100 ºC overnight. The reaction solution was dried via rotary evaporation, and the crude compound was redissolved in a DCM:methanol (1:1) solution. The solution was stirred with 0.5 g of Amberlyst 15 strongly acidic cation exchange resin. The resin was washed three times with a DCM:methanol (1:1) solution. The compound was eluted with 2M ammonia in methanol (30 mL), and the filtrate was dried. The filtrate was purified by column chromatography with DCM:Methanol (10:1) gradient as the eluent. Yellow powder with yield of 23 mg (0.072 mmol, 7.2%). $^1$H NMR (500 MHz, DMSO) δ 7.30 (td, $J$ = 7.8, 2.6 Hz, 2H), 7.16 (d, $J$ = 3.4 Hz, 1H), 6.91 (d, $J$ = 7.9 Hz, 1H), 6.85 (d, $J$ = 11.6 Hz, 2H), 6.64 (d, $J$ = 3.5 Hz, 1H), 6.06 (s, 1H), 5.45 – 5.39 (m, 2H), 4.55 (d, $J$ = 5.2 Hz, 2H), 3.90 (d, $J$ = 2.5 Hz, 2H). $^{13}$C NMR (126 MHz, DMSO) δ 190.3, 167.8, 163.8, 162.5, 158.5, 157.3, 154.3, 149.1, 144.1, 141.5, 139.7, 136.8, 132.0, 130.9, 129.2, 128.3, 120.7, 118.7,

113.6, 112.8, 109.5, 90.1, 56.3, 52.4, 32.9. HPLC retention time: 3.9 minutes. Chemical Formula: $C_{18}H_{13}N_3O_3$. Exact Mass: 319.10. HRMS: $[M+H]^+$ = 320.1033 m/z.

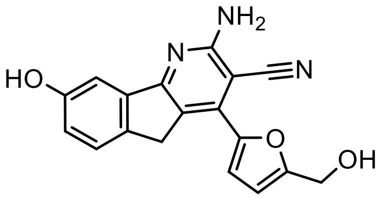

**KI-TOX-A3-0178**

5-(hydroxymethyl)furan-2-carbaldehyde (151 mg, 1.2 equiv) was combined with 6-hydroxy-2,3-dihydro-1H-inden-1-one (148 mg, 1 equivalent), ammonium acetate (350 mg, 4.5 equiv), and malonitrile (66 mg, 1 equiv). The mixture was dissolved in ethanol and heated to 90-100 ºC overnight. The reaction solution was dried via rotary evaporation, and the crude compound was redissolved in a DCM: methanol (1:1) solution. The solution was stirred with 0.5 g of Amberlyst 15 strongly acidic cation exchange resin. The resin was washed three times with a DCM: methanol (1:1) solution. The compound was eluted with 2M ammonia in methanol (30 mL), and the filtrate was dried. The filtrate was purified by column chromatography with DCM: methanol (10:1) gradient as the eluent. Yellow powder with yield of 50 mg (0.156 mmol, 15.6%) $^1$H NMR (500 MHz, DMSO) δ 7.46 (d, $J$ = 8.2 Hz, 1H), 7.39 (d, $J$ = 3.5 Hz, 1H), 7.25 (d, $J$ = 2.3 Hz, 1H), 6.92 (dd, $J$ = 8.1, 2.4 Hz, 1H), 6.87 (s, 2H), 6.62 (d, $J$ = 3.5 Hz, 1H), 4.56 (s, 2H), 3.95 (s, 2H). $^{13}$C NMR (126 MHz, DMSO) δ 163.6, 162.5, 158.5, 157.4, 147.9, 141.0, 136.9, 136.6, 126.4, 121.7, 118.8, 118.5, 115.4, 109.9, 107.5, 80.8, 56.3, 34.8. HPLC retention time: 3.8 minutes. Chemical Formula: $C_{18}H_{13}N_3O_3$. Exact Mass: 319.10. HRMS found: $[M+H]^+$ = 320.1028 m/z.

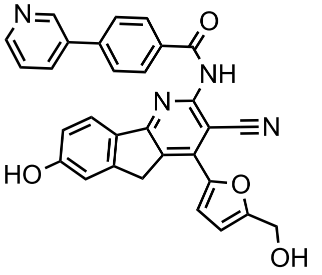

**KI-TOX-A3-0245**

KI-TOX-A3 (15mg, 0.047mmol) was mixed with 4-Pyridin-3-yl-benzoic acid (15mg, 0.075mmol, 1.6eq) in 2ml of DMF solvent. The HATU (50mg, 0.13mmol, 2.8eq.) was added to the solution and the system was purged with Argon for 5 minutes. Then, the DIPEA (10v/v%, 0.2ml) was added to the reaction. The stirring lasted over-night and the TLC indicates a completion of the reaction. The compound was worked up with water (100ml) and extracted by DCM (50ml x 3). After combination of organic layers and evaporation, the compound was purified by combi-flush column system with DCM: Methanol (20:1) and a pale-yellow compound was obtained (3.6mg, 0.0072mmol, 15.3%). $^1$H NMR (500 MHz, DMSO) δ 9.03 (d, $J$ = 2.4 Hz, 1H), 8.67 (dd, $J$ = 4.7, 1.6 Hz, 1H), 8.42 (s, 1H), 8.28 (d, $J$ = 8.3 Hz, 2H), 8.23 (dt, $J$ = 8.0, 2.0 Hz, 1H), 8.03 (d, $J$ = 8.2 Hz, 2H), 7.95 (d, $J$ = 8.2 Hz, 1H), 7.66 (d, $J$ = 2.1 Hz, 1H), 7.58 (dd, $J$ = 8.0, 4.8 Hz, 1H), 7.44 (dd, $J$ = 9.0, 2.7 Hz, 2H), 6.95 (s, 1H), 6.65 (d, $J$ = 3.5 Hz, 1H), 4.58 (s, 2H), 4.16 (s, 2H), 2.55 (s, 1H). $^{13}$C NMR (126 MHz, DMSO) δ 162.6, 158.7, 152.5, 150.0, 148.4, 148.0, 142.9, 136.9, 135.1, 131.1, 128.8, 127.9, 124.5, 122.5, 121.9, 120.9, 119.6, 115.7, 109.9, 99.2, 91.2, 56.4, 29.3. HPLC retention time: 4.5 minutes. Chemical Formula: $C_{30}H_{20}N_4O_4$. Exact Mass: 500.15. HRMS found: $[M+H]^+$ = 501.2678 m/z.

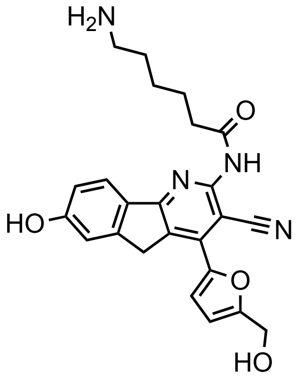

**KI-TOX-A3-0239**

KI-TOX-A3 (15mg, 0.047mmol) was mixed with Fmoc-epsilon-Acp-OH (25mg, 0.071mmol, 1.5eq) in 2ml of DMF solvent. The HATU (50mg, 0.13mmol, 2.8eq) was added to the solution and the system was purged with Argon for 5 minutes. Then, the DIPEA (10v/v%, 0.2ml) was added to the reaction. The stirring lasted overnight, and the TLC indicates a completion of the reaction. The compound was worked up with water (100ml) and extracted by DCM (50ml x 3). After combination of organic layers and evaporation, the compound was purified by Combi-flush column system with DCM: Methanol (20:1) and pale-yellow compound was obtained (10.2mg, 0.0156mmol, 33.2%). Then, this compound was dissolved into 1ml DMF again with stirring. By adding Piperidine (0.1ml) into the solution, the solution turned into bright yellow. After 1h stirring, the Fmoc protection group was removed based on the TLC spotting. Then, this solution was worked up by 100ml water and extracted by DCM again with 50ml x3. The organic layers were collected and evaporated. To completely remove the DMF, 50ml Toluene was again added to the mixture and evaporated by rotovap at 45C. Then, the compound was purified by HPLC with a gradience of water and acetonitrile (+0.1% formic acid). The compound was further frozen into -80C and dried by lyophilized overnight. The final product is yellow powder (1.9mg, 0.0044mmol, 9% yield) 1H NMR (500 MHz, MeOD) δ 7.81 (d, J = 8.4 Hz, 1H), 7.42 (d, J = 3.5 Hz, 1H), 7.06 (s, 1H), 6.91 (d, 2.3 Hz, 1H), 6.75 (d, J = 3.6 Hz, 1H), 5.28 (s, 2H), 4.61 (s, 2H), 4.01 (s, 1H), 2.93 – 2.86 (t, J=7.7 Hz, 2H), 2.48 (t, J = 7.3 Hz, 2H), 1.76 – 1.60 (m, 6H), 1.46 (t, *J* = 2.9 Hz, 2H). HPLC retention time: 3.75 minutes. Chemical Formula: $C_{24}H_{24}N_4O_4$. Exact Mass: 432.18. MS: $[M+H]^+$ = 433.1873 m/z.
