## Supplementary data for "Small molecule modulators of TOX protein re-invigorate T cell activity"

1. *ELISA screening of the hits*

We discovered 15 inhibitors to the protein-protein interaction of TOX-KU70/80 with the chemical structures shown in the figure S1A. **KI-TOX-A3** and **KI-TOX-H05** shared a similar tricyclocyano pharmacophore, though the H05 has milder inhibition activity than **KI-TOX-A3**. The compounds **KI-TOX-L09**, **KI-TOX-E16**, **KI-TOX-L06**, **KIT-TOX-B12**, **KI-TOX-I22** shared a the structural features of a five-member ring conjugated to another five or six-member ring and an amide bond. They all showed promising TOX PPI inhibition activities.

Note that, initially, **KI-TOX-C22** showed very strong inhibition of TOX binding to KU70/80 (IC50 ~ 40 nM) (Fig.S1B). Its derivatives **KI-TOX-P4, KI-TOX-N13**, **KI-TOX-05-P14** all showed relatively strong inhibition. To determine whether **KI-TOX-C22** was worth further pursuit, the purity of the molecule was examined by HPLC and we noticed that the compound showed another peak in the HPLC (Fig. S1C). Subsequently, a new batch of pure **KI-TOX-C22** was synthesized and we noted that the compound is not stable at room temperature and in light. After a few days exposure, the compound turned green from pale yellow in a DMSO solution. We hypothesize oxidation in the core 6-member ring with amine. The purified version of **KI-TOX-C22** did not result in significant TOX PPI inhibition, while the potency increased in the degraded version.

Figure S1B and figure 1G showed that the **KI-TOX-D22** is not as strong with respect to inhibition. Nonetheless, it is one of the most pure and stable chemicals in the library, and the TOX PPI inhibition is consistent and repeatable. As a result, we chose it as the mild PPI inhibitor in the T cell exhaustion study. I THINK THIS IS AT ODDS WITH THE TEXT ABOVE WHICH QUESTIONS PURITY AND STABILITY. I'M CONFUSED.

As shown in Figure 1G, KI-TOX-P14 demonstrates PPI modulation at 0.125 uM. To ensure if it binds to TOX, we probed a SPR analysis, which indicates kd=0.79 uM (Fig. S1D). At high concentrations, inhibition was adversely affected in both the PPI and PLA assay (Figure 2A). This chemical would need further investigation on its mechanism of action in TOX protein.

A

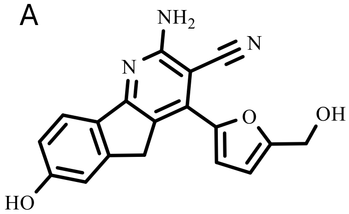

KI-TOX-A03
1351116-34-1

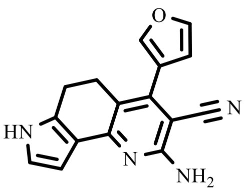

KI-TOX-H05
1351045-99-2

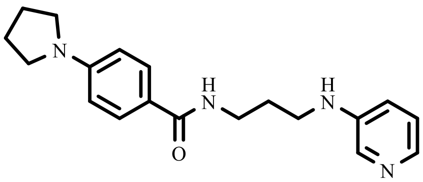

KI-TOX-P14
1783285-79-9

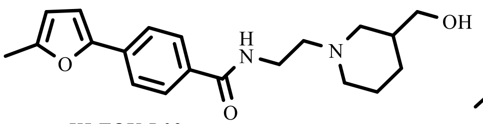

KI-TOX-L09
2179318-44-4

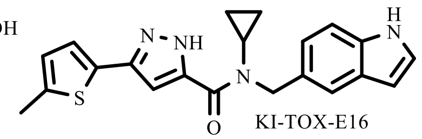

KI-TOX-E16
1467747-55-2

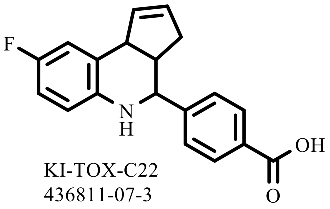

KI-TOX-C22
436811-07-3

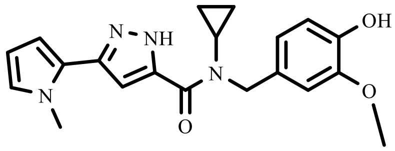

KI-TOX-L06
1470374-30-1

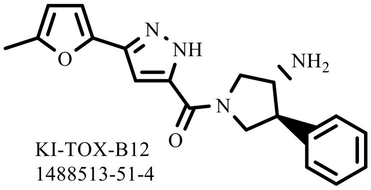

KI-TOX-B12
1488513-51-4

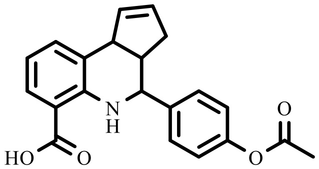

KI-TOX-05-P14
439294-98-1

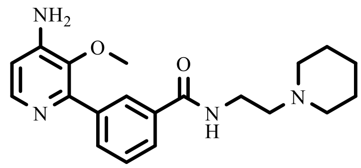

KI-TOX-D22
2178334-94-4

KI-TOX-B04
1069616-85-8

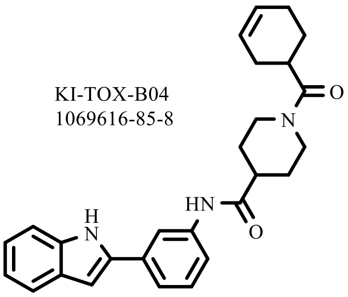

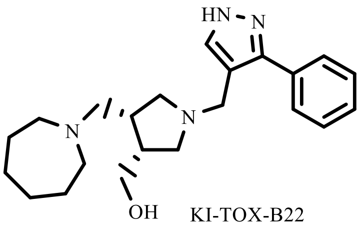

KI-TOX-B22
1422880-95-2

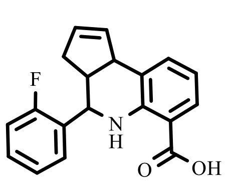

KI-TOX-P4
360790-06-3

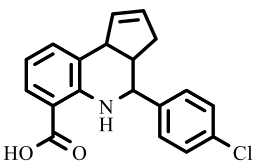

KI-TOX-N13
473265-37-1

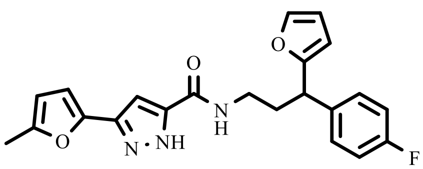

KI-TOX-I22
1486621-72-0

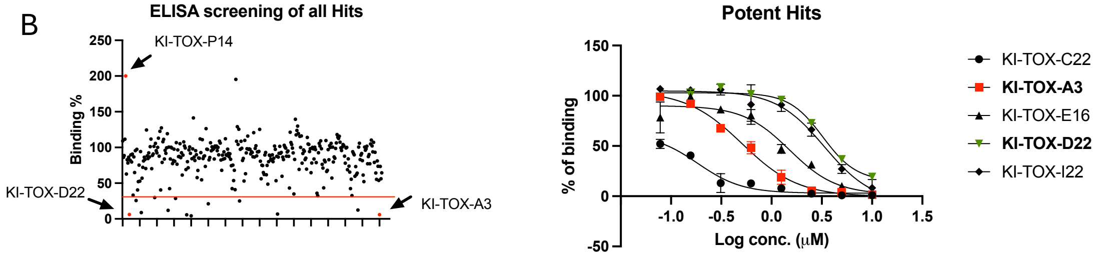

C.

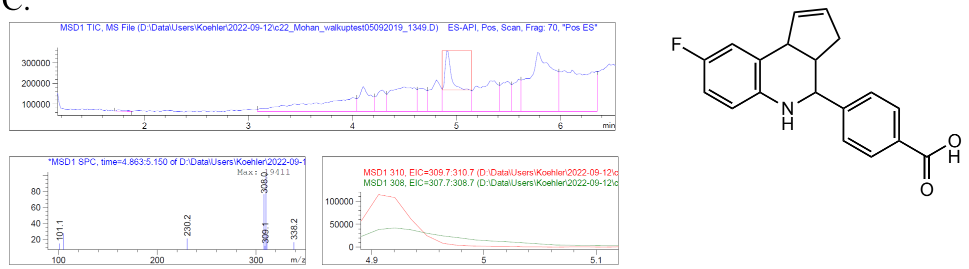

Exact Mass: 309.12

Possible potent form:

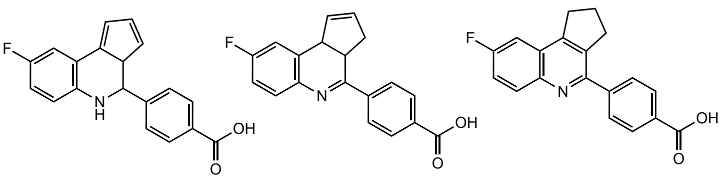

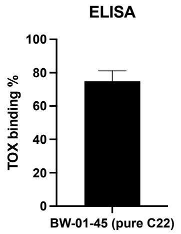

Exact Mass: 307.10

| Candidates | Cell cytotoxicity in μM | | | | | |
|---|---|---|---|---|---|---|
| | Molt4 (μM) | Jurkat | HBP-ALL | HH | Hut78 | Raji |
| KI-TOX-C22 | NI[b] | NI | NI | NI | NI | NI |

D. SPR of **KI-TOX-P14**

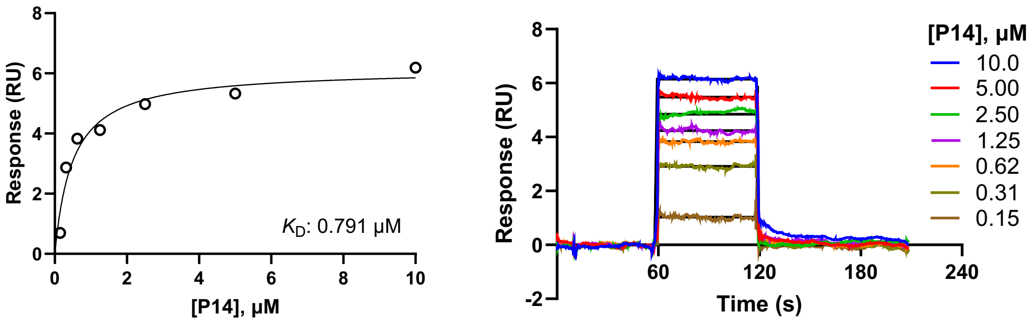

Figure S1. (A) the hits that showed inhibition in the ELISA assay. (B) The ELISA screening data. (C). KI-TOX-C22 is not as potent in its pure form. (D). SPR binding data for KI-TOX-P14.

2. *Synthesis of KI-TOX-A3 and derivatives.*

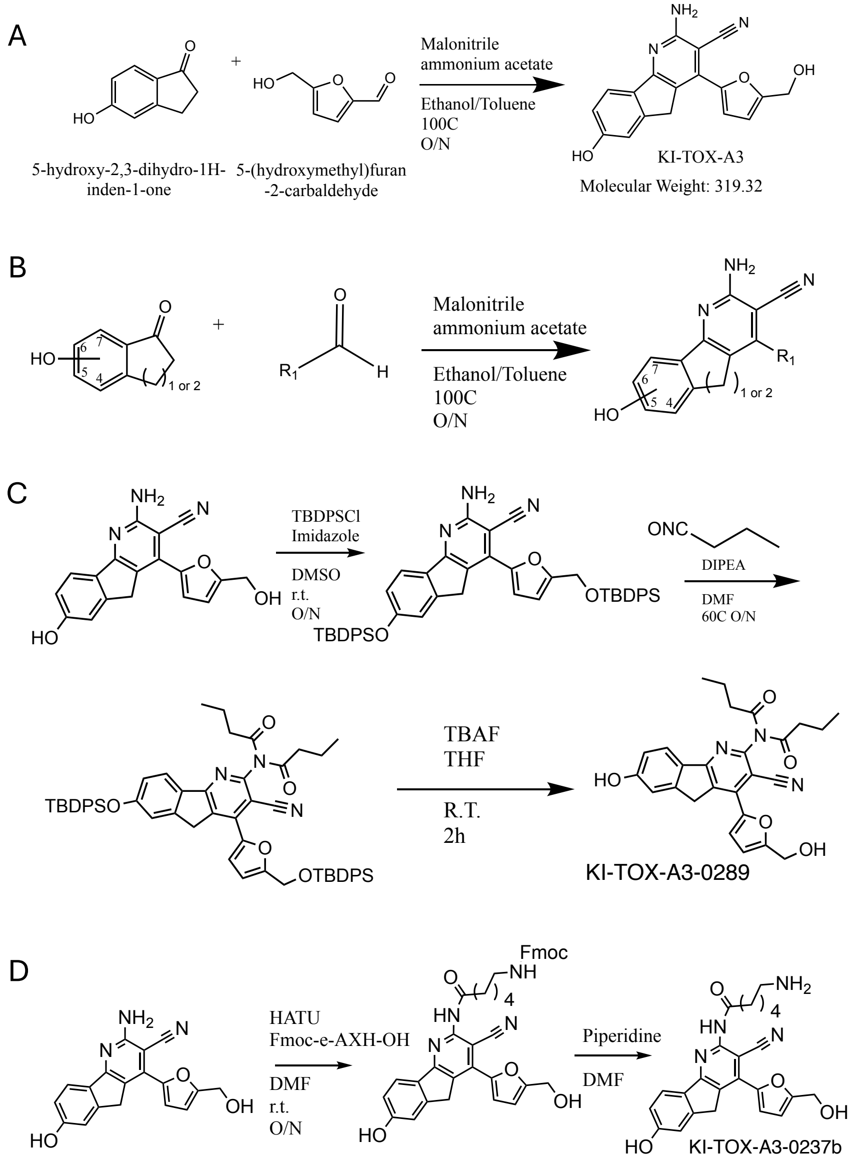

Figure S2. The synthesis methods of the derivatives of **KI-TOX-A3**. (A) The synthesis method of **KI-TOX-A3**; (B) The generalized scheme for **KI-TOX-A3** derivatives with relocation of hydroxyl group on phenol and change of size for the indanone ring. (C). Protection of the -OH groups on **KI-TOX-A3**, followed with isocyanate conjugation. (D) Coupling reaction to make chemical probe of **KI-TOX-0237b.**

Compounds provided herein were synthesized according to the methods disclosed in international patent application publication No. WO 2004/054505 A2 to Pharmacia Corporation/Reinhard[1]. Unlike the protocol described in the patent using dichloromethane or a mixture of toluene and benzene (66%:33%), the use of mixture of Toluene and ethanol (20%/80%) increased the yield significantly. Additionally, we found that an excess amount of ammonia acetate is essential (5-10 eq.) for the reaction. The product **KI-TOX-A3** partially crashes out from the solvent system but a simple filtration alone does not yield pure compound. An acidic resin ionic exchange step is necessary prior to silica gel column purification. After purification, **KI-TOX-A3** was diluted in DMSO or acetonitrile to 100μM and stored in -20°C.

The SAR study involved substitution of the indanone or the furan groups with the groups listed in Figure S3. We also explored expansion of the five-member ring in the indanone to a 6-member ring. As shown, we also modified the amine moiety of the parent compound to monitor impact on potency. Synthesis of modified compounds followed the same general reaction scheme as outlined for **KI-TOX-A3** synthesis, using different building blocks. In the case of **KI-TOX-A3-0189,** we utilized reaction of the propyl isocyanate with the amine group of the TBDPS-protected **KI-TOX-A3** and heat. The intermediate was deprotected using TBAF in THF solution. HATU catalysis was used to directly form an amide bond with **KI-TOX-A3** for selected compounds. For example, **KI-TOX-A3-0237b** was prepared after HATU coupling and deprotection of Fmoc by piperidine.

After probe synthesis, we tested the derivatives in the PPI assay and found that KI-TOX-A3-0174, 77, 82, and 89 showed calculable IC50s. We also noticed that the substitution of furan to benzyl group leads to significant reductions in cytotoxicity (**KI-TOX-A3-0180, KI-TOX-A3-0184, KI-TOX-A3-0186**) The substitution of furan to pyrrole and thiophene does not improve the potency of TOX inhibition or cancer cytotoxicity. Also, the positioning of the hydroxyl group of the phenol group of **KI-TOX-A3** is critical. $7^{th}$ position hydroxyl group (number noted on Fig. S2B) revealed TOX inhibition while $4^{th}$ position hydroxyl group would loss TOX inhibition. While the SAR conducted to date is preliminary and should be expanded to understand fine relationships, among all derivatives, **KI-TOX-A3-0189** revealed the most potent activity in TOX inhibition. This suggests that the amine is not associated with protein pocket interaction and is a suitable site to extend with conjugation reactions (e.g., exit vector). One of the applications enabled by conjugation is the development of a chemical pulldown probe. Therefore, we immobilized KI-TOX-A3 at this amine group with Fmoc-€-AXH-OH group extension and covalent coupling with and NHS- Mag Sepharose.

Looking forward to additional chemistry, this permissive amine will enable modification opportunities, including PROTAC designs. Degradation of TOX might be of interest for both therapeutic purposes related to T cell exhaustion, or molecular mechanistic studies of TOX in both T cells and leukemia cells.

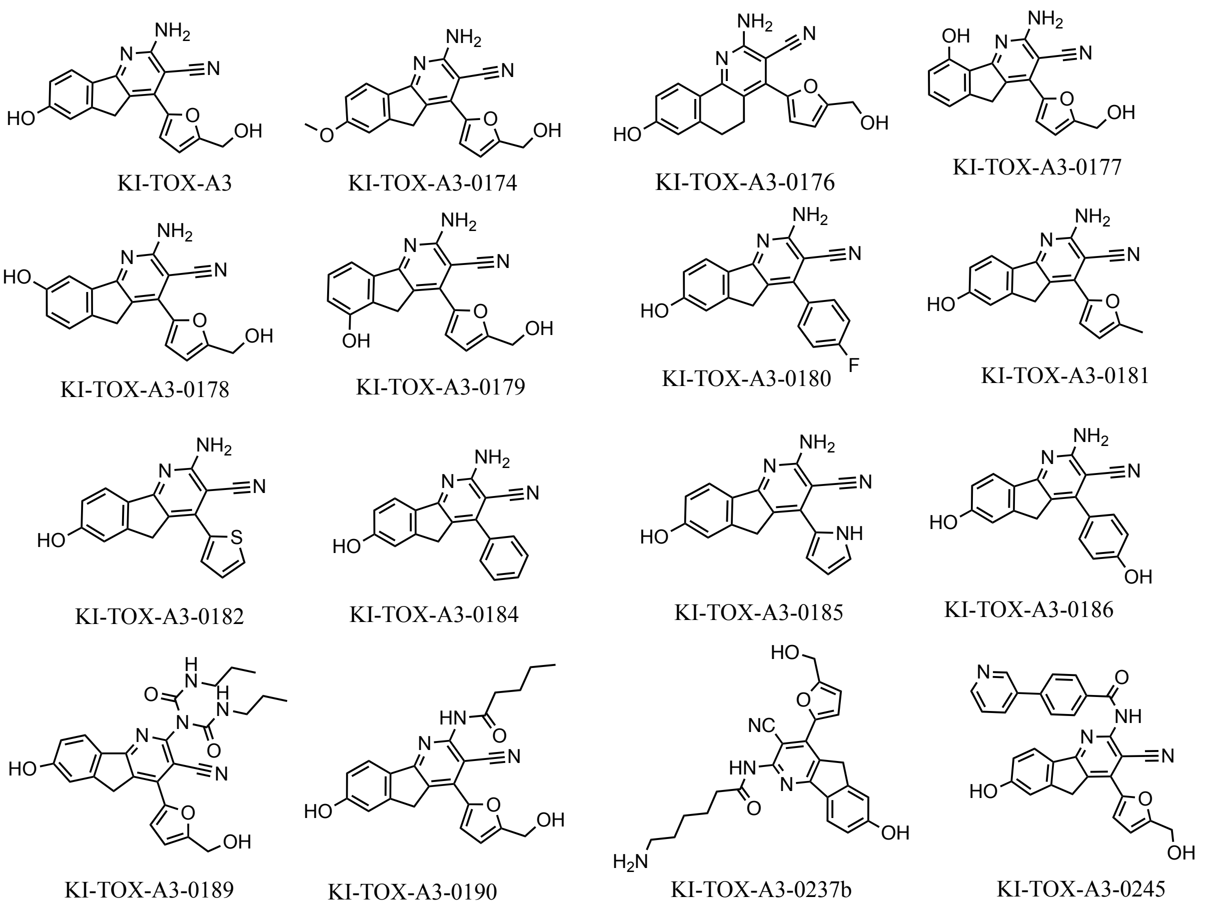

Figure S3. The list of derivates of **KI-TOX-A3** based on the Fig. S2.

*3. TOX protein purification:*

After SMM screening, the TOX protein was required for selected secondary assays. While ELISA assays require only low-microgram quantities of TOX, our biophysical studies required larger quantities of TOX protein to enable both assay development and the final studies.

Initially, we chose HEK293T mammalian expression for TOX as a FLAG-tagged protein. However, we noticed that the mammalian recombinant TOX protein had very low yield (5ug/5million HEK293T) even with relatively pure FLAG-TOX protein via affinity purification. We suspected the loss of TOX protein relates to aggregation on surfaces when we tried to remove the FLAG peptide by buffer exchange. TOX is a highly disordered protein and may stick to membranes or penetrate the MWCO filter. With consistently low yields across multiple attempts, we decided to pivot into *E. coli* expression protocols.

To enable bacterial expression, we constructed the protein sequence with the backbone of PSMT3 from Addgene (191261) and cloned an AVI-TOX sequence into the backbone. We chose this backbone because it contains a His-SUMO tag. SUMO tags are widely applied in disordered protein expression because the misfolding protein could easily be translocated to inclusion bodies, and the SUMO tag can partially increase the protein solubility.

Initially, we chose BL21 as our expression strain and unfortunately, the BL21 (DE3) strain is not compatible with our plasmid, as the protein leak is severe and IPTG induction does not grant large expression in the strain. As a result, we moved to the lemo21 c43 strain. This strain has a lemo21 plasmid which can prevent protein leaking by expressing T7 lysozyme as the natural inhibitor of T7 ribosome. In addition, the c43 lacks selected proteases, which could help prevent some potential truncations in the soluble TOX.

We explored a variety of expression conditions for c43 strain, including variations in temperature time, L-rhamnose concentrations (the sugar to induce Lemo21 plasmid expression), optimal OD600 and more. We found that there is no significant difference between 30°C and 37°C as the IPTG has successfully induced protein expression (Fig. S2A). In addition, in the uninduced group, there is no leak of TOX protein. Later, we tested the concentration of L-Rhamnose and surprisingly, the concentration 0 mM (No L-Rhamnose) gave the highest level of TOX induction (Fig. S2B). We hypothesize that the Lemo21 has a mild leakage of T7 lysozyme, and the leak already offers sufficient inhibition of T7 Ribosome activity. Lastly, we test the expression of TOX in TB and LB media. We also noticed that at the cell density corresponding to OD600 of 1.6, TOX protein levels are high (Fig. 2C).

Upon optimization of the protein expression, TOX solubility and purity were still issues. The TOX protein was typically truncated before cell lysis. More TOX appeared in inclusion bodies (IBs). Protein extraction and purification from the IBs helped to obtain pure His-SUMO-AVI-TOX protein (yield = 500ug/1L) (Fig. S2D). The cell lysis and protein extraction protocols are provided in the methods section.

After washing out guanidine and non-specific binders, TOX was refolded and eluted from 500 mM imidazole. We attempted to purify the protein further with size exclusion chromatography and noted that TOX is not compatible with Superdex 200 size-exclusion column, as the protein was trapped in the column with dramatic loss (80-90% loss). Therefore, we avoided using SEC purification in later runs. We next attempted a protocol involving dialysis instead of centrifuge filtration, SEC, or desalting column was applied, as we found all these methods caused high loss of protein levels. For dialysis, the buffer pH was critical – white precipitation was observed in PBS dialysis (pH = 7.4) while HEPES buffer with pH = 8.1 prevented protein precipitation. The TOX PI is 7.4. Therefore, a pH>8 buffer is ideal for the TOX in dialysis and storage.

Later, to cleave the SUMO tag, we incubated the SUMO protease (10% (w/w) of the TOX protein) with the His-SUMO-AVI-TOX and incubated at 4C overnight. Then, to purify the SUMO cleaved from TOX and excessive SUMO protease, we incubated 0.1g of equilibrated NTA resin to 500ug of the cleaved protein for 10 minutes. After centrifugation, pure AVI-tagged TOX protein was obtained. Then, to biotinylate the AVI-tagged TOX, we dialyzed the TOX protein with potassium glutamate salt (100 mM) and HEPES (10mM, pH = 8) to remove the NaCl, as the BirA enzyme was not compatible with an NaCl buffer. Then, we incubated the dialyzed protein with 10 ug BirA enzyme and the Buffer A, B, and C provided from the BirA biotinylation kit (Avidity). The protein was biotinylated as observed by western blot  (Fig.S3E)

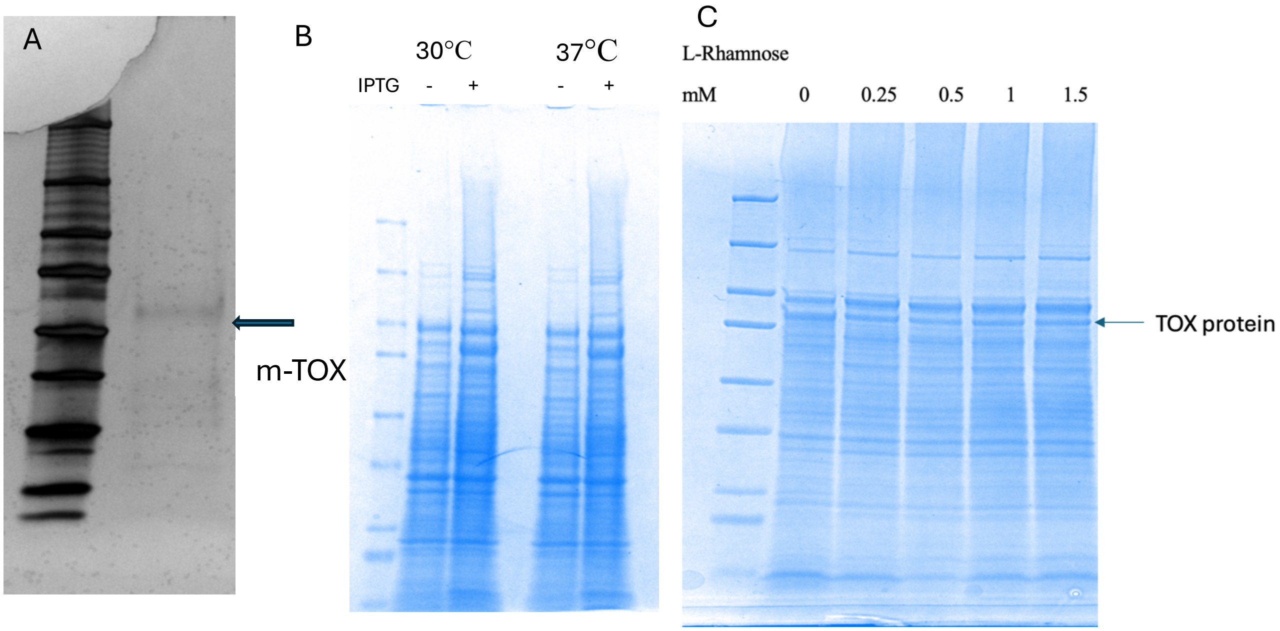

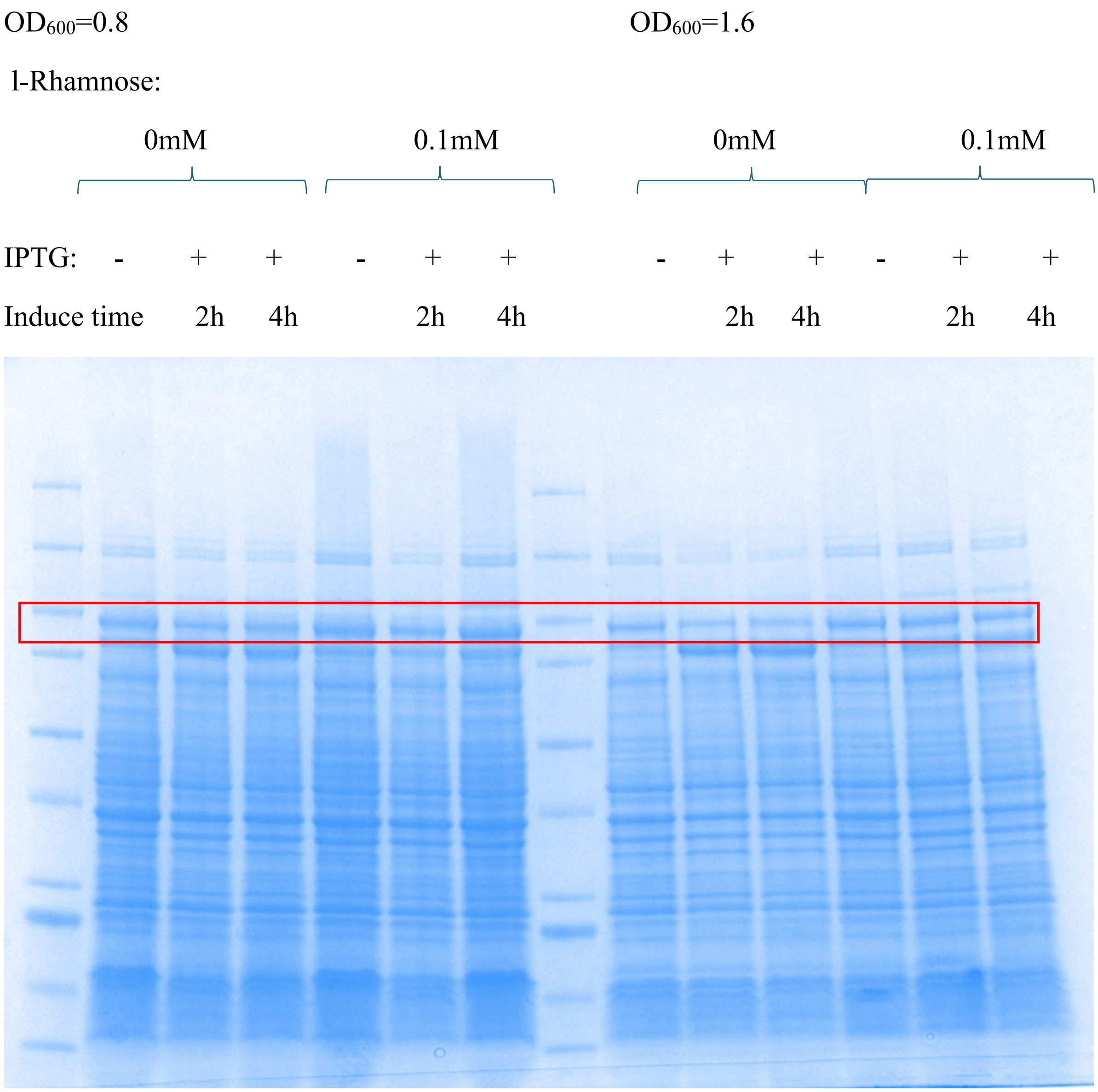

$OD_{600}$=0.8
$OD_{600}$=1.6
l-Rhamnose:
0mM
0.1mM
0mM
0.1mM
IPTG: - + + - + + - + + - + +
Induce time 2h 4h 2h 4h 2h 4h 2h 4h

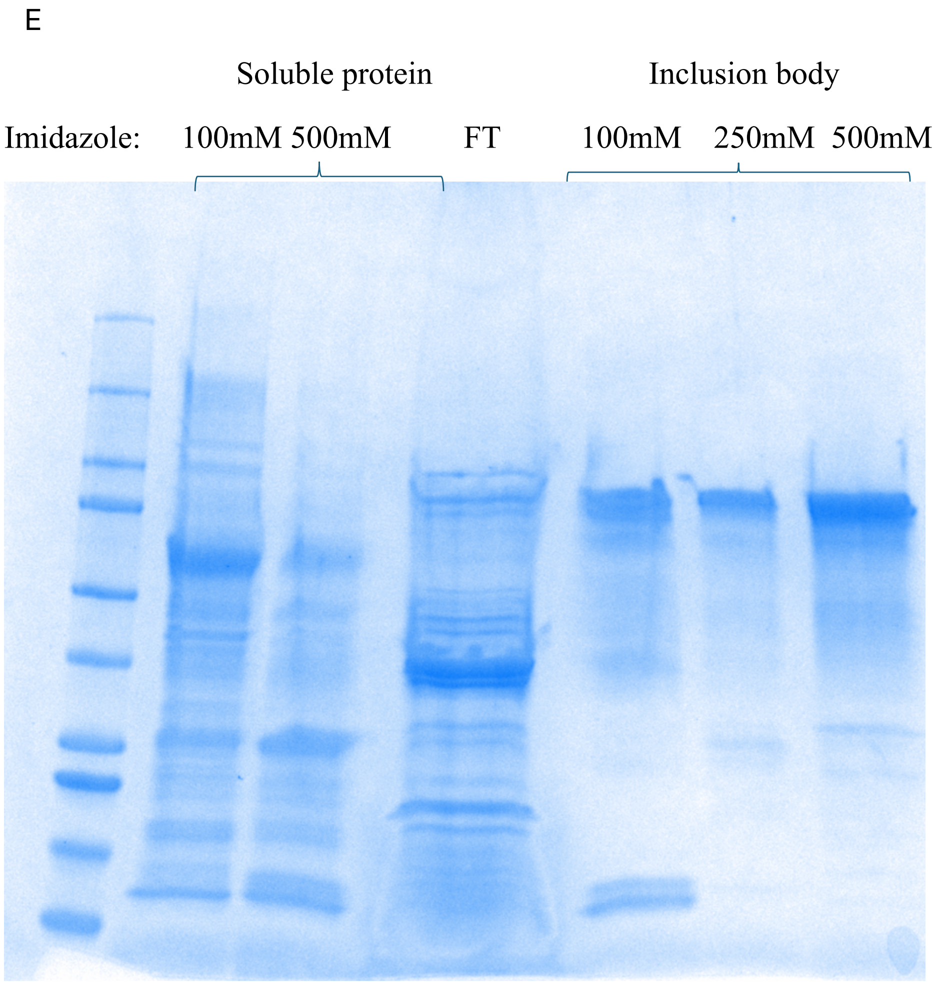

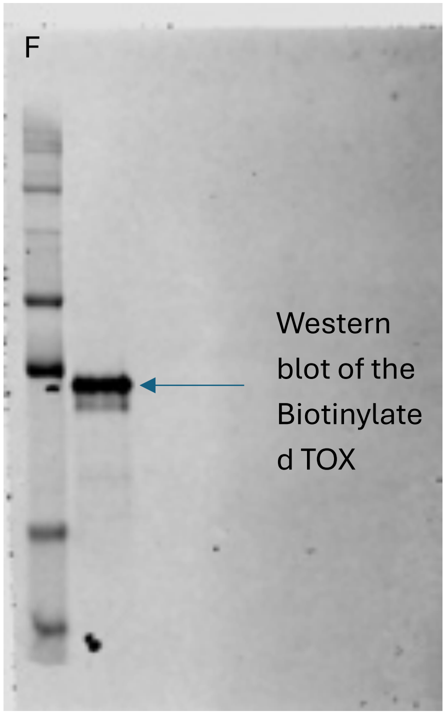

Figure S4. (A). mammalian TOX expression through HEK293-T as the host. (B-F). Protein expression using *E.Coli* strain (c43) under different conditions. (B) Temperature determination. (C). L-Rhamnose concentration determination; (D). The initial induction OD600 determination. (E) soluble versus inclusion body extraction; (F). The streptavidin western blot showed pure biotinylated TOX without SUMO tag.

4. KI-TOX-A3 can downregulate TOX protein in mammalian cells (Full gels)

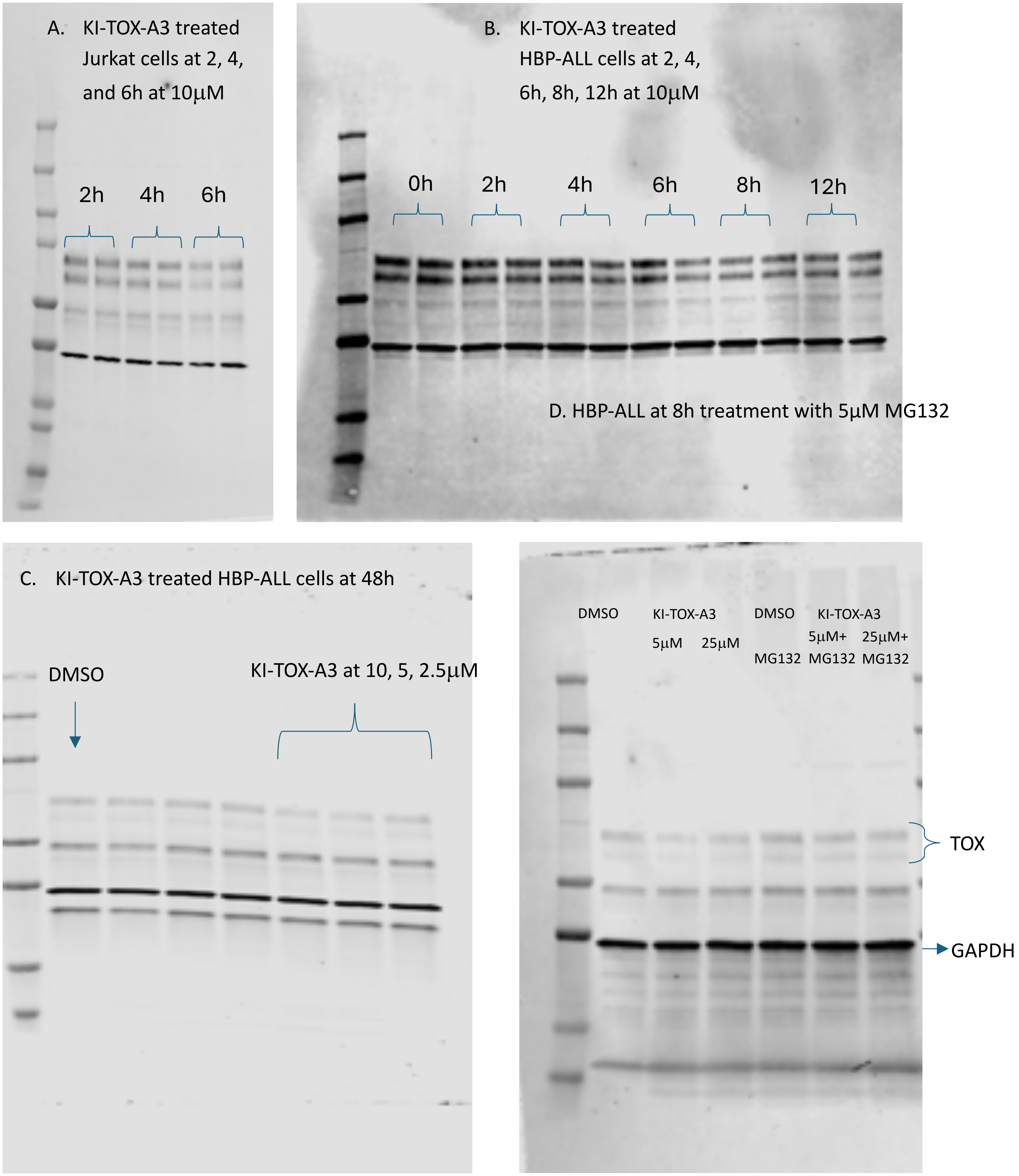

E. Molt4 at 12h treatment with 5µM MG132

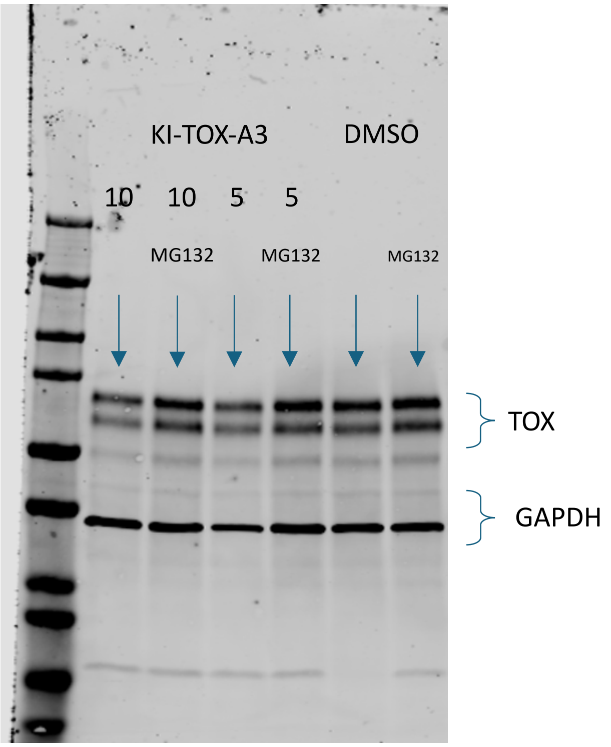

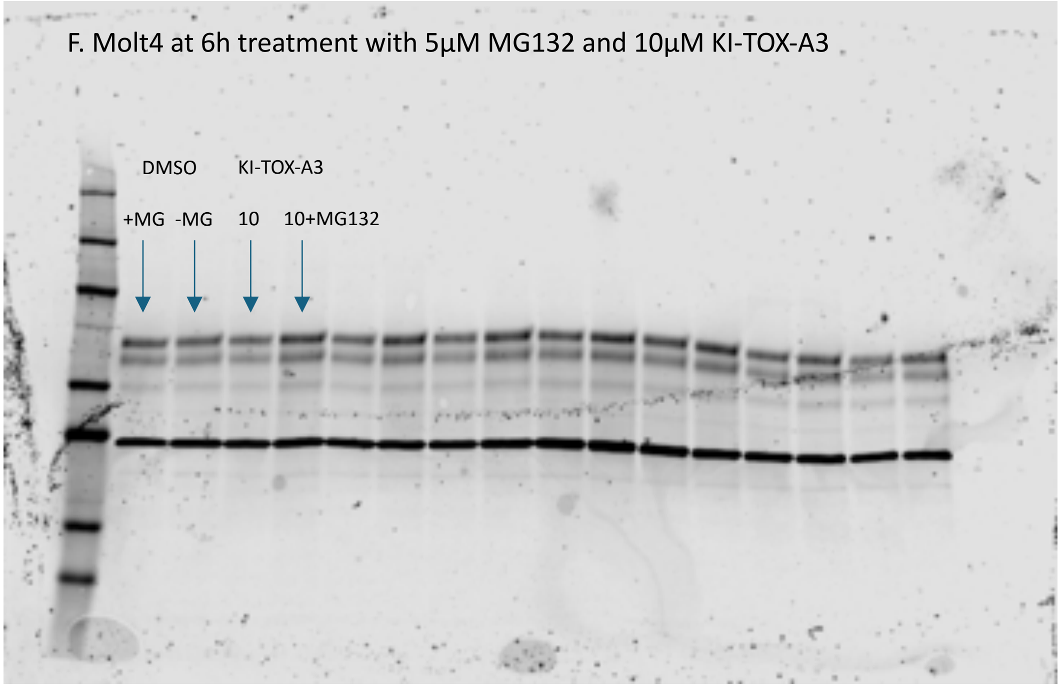

Figure S5. TOX level was downregulated by KI-TOX-A3. (A) **KI-TOX-A3** treated Jurkat cells at 2, 4, and 6h at 10uM; (B) **KI-TOX-A3** treated HBP-ALL cells at 2, 4, 6h, 8h, 12h at 10uM. (C). **KI-TOX-A3** treated HBP-ALL cells at 48h; (D). HBP-ALL at 8h treatment with 5µM MG132; (E). Molt4 at 12h treatment with 5µM MG132. (F). Molt4 at 6h treatment with 5µM MG132.

KI-TOX-A3 has shown mild downregulation of TOX protein levels in different cell lines, with dependence on both dose and time. TOX levels started to decrease as early as 4h and remain around 50% around from 8h to 48h. We also noticed that the figure S3D and S3E showed some reverse effect at 25µM in 8h treatment. Interestingly, MG132, a potent proteosome inhibitor, can rescue TOX levels, which suggests the proteasome plays some role in the alterations of TOX levels, rather than only modulating the transcription of TOX. The precise mechanism of this post-translational perturbation of TOX levels is under investigation.

*5. KI-TOX-A3 probes potential TOX PPIs in T-ALL cell in MS experiments*

| **Example proteins** | **Log(p-value) (A3 vs. DMSO)** | **Fold change (A3 vs. DMSO)** |
|---|---|---|
| HSP90AB1 | 2.05461654 | -5.1292265 |
| HSP90AA1 | 1.57043985 | -5.9136144 |
| HSPB1 | 1.23922025 | -4.8614066 |
| eEF2 | 1.13653742 | -3.7993851 |
| S100A7 | 4.28604107 | -5.7927094 |
| S100A12 | 1.21090216 | -3.8812444 |
| ERO1 | 4.92499678 | -5.2655412 |
| KU70 | 1.57316724 | 3.38368213 |
| AZU1 | 3.49383303 | -9.7131314 |
| CTSB | 2.4845755 | -4.2935313 |
| Histone H2A | 1.34492624 | -0.9957373 |
| Macro-H2A | 1.32915964 | -4.787635 |
| Histone H1.0 | 1.23617318 | -4.9560151 |

Table S1. Putative protein interactors of TOX from co-IP experiments +/- KI-TOX-A3. Note that the hits were identified after two independent repeated IP-MS experiments.

In addition to the heat shock proteins (HSPs) discussed in the manuscript, **KI-TOX-A3** also enabled us to develop hypotheses about other potential TOX PPIs in Molt4 cells (Table. S1). For example, we found Histones H2A and H1.0 with relatively significant difference between KI-TOX-A3 treated group and DMSO-treated Molt4 lysate based on the p-value and fold change significance. The histone binding could be direct or indirect. Further investigation should be proceeded for TOX role in this interaction. Furthermore elongation factor EFF2 might be another interested hit in this proteomic study. Elongation factors are responsible for aminoacyl-tRNA delivery to ribosome and peptidyl-tRNA translocation from site A to P in the ribosome[2]. The eEF2 is overexpressed in leukemia cells and indicated as a therapeutic target[3]. Later, we noted that members of the protein S100 series are pulled down with the anti-TOX bead and are eliminated with KI-TOX-A3 inhibition. The S100 family modulates cellular signals as intracellular Ca2+ sensors and as extracellular factors [4], and overexpression is associated with T-ALL[5]. Many other hits including ERO1, AZU1, CTSB are examples of hits playing unknown TOX partners in Molt4 or other leukemia cells. Further validation of the PPIs are needed.

### 6. TOXi can reverse the T cell exhaustion status.

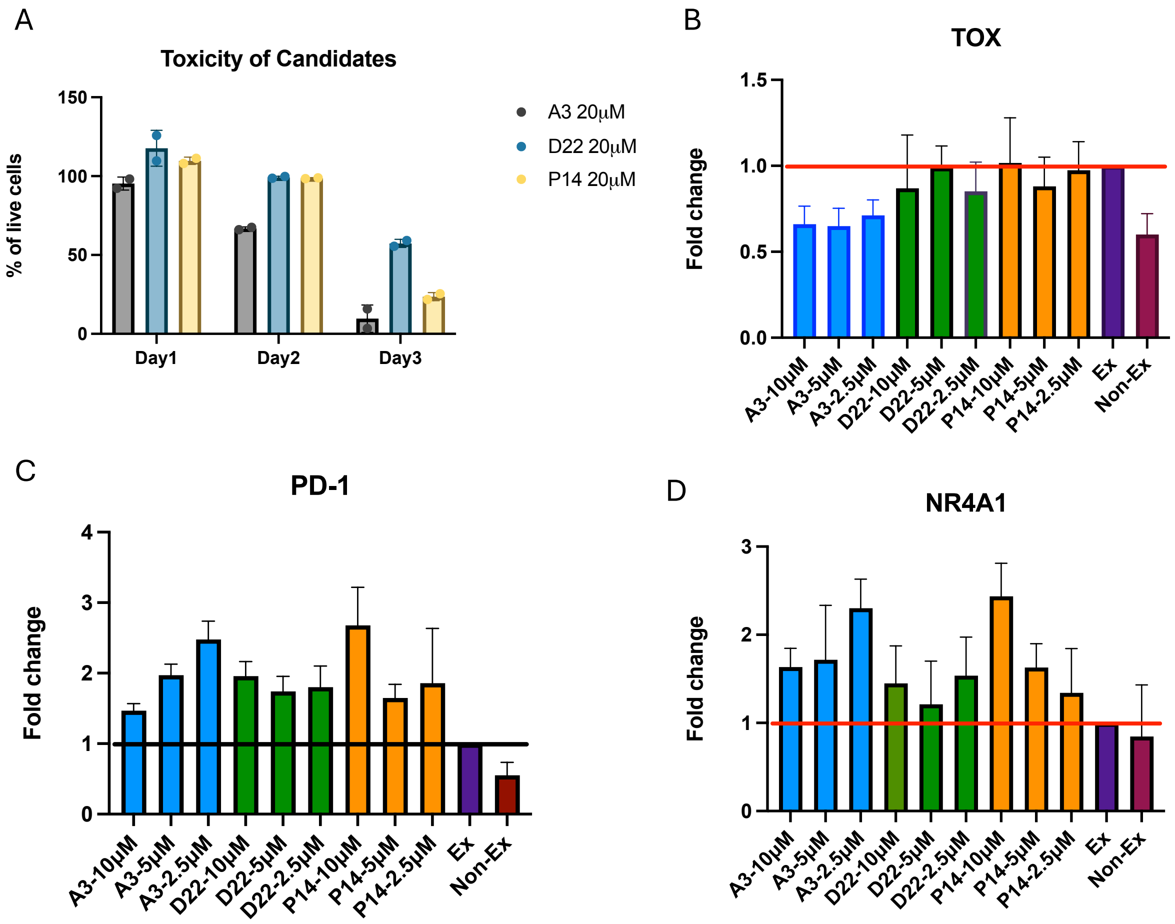

E

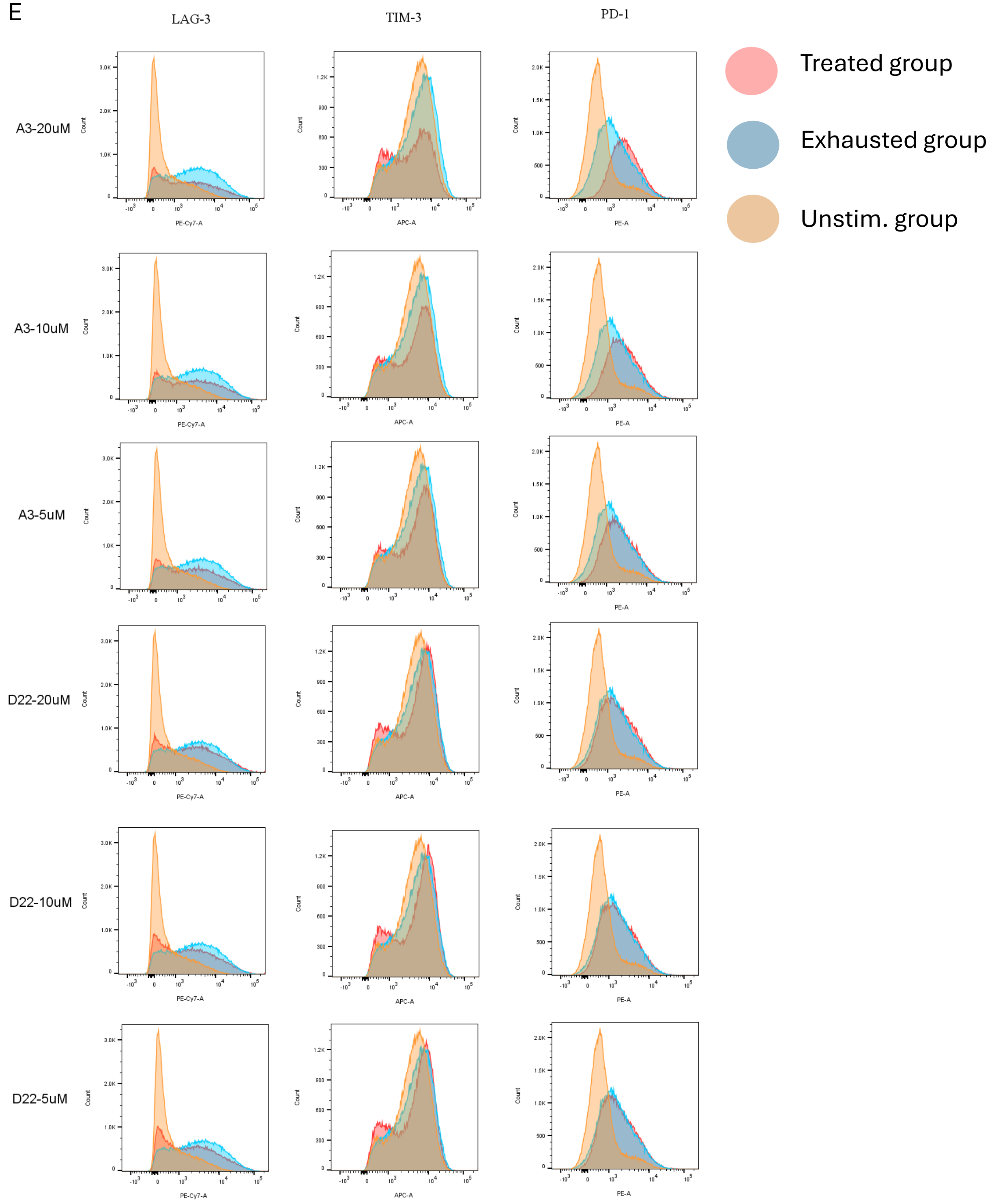

LAG-3
TIM-3
PD-1
Treated group
Exhausted group
Unstim. group
A3-20uM
A3-10uM
A3-5uM
D22-20uM
D22-10uM
D22-5uM
Count
PE-Cy7-A
APC-A
PE-A

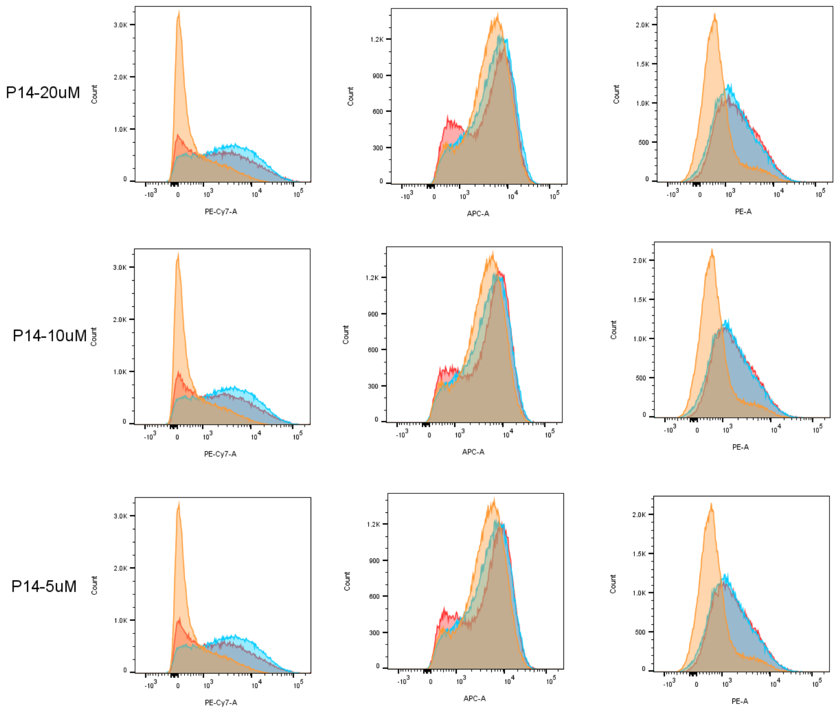

Figure S6. The TOX PPI inhibitors re-invigorate T cells from exhaustion. (A) Cytotoxicity at 20uM of the selected candidates. (B) mRNA level changes on *Tox*, (C) *pdcd-1*, and (D) *nr4a1*. (E) The biomarker PD-1, TIM-3, and LAG3 change with candidates at different concentrations (cycle $6^{th}$, 24h treatment).

Initially, we stimulated CD8+ T cells with Dynabead CD3/28 stimulator. Cells grew rapidly and expanded daily for the first ten days. However, the inhibitory receptors were not co-upregulated until day 20. Such a long process of cell culture reduced reproducibility, and we experienced issues with cell death. To resolve these technical concerns, we tried STEMCELL Technologies T cell activator, as this activator is soluble in liquid and stimulates T cells efficiently[6]. As expected, we have observed stronger exhaustion biomarker expression in the STEMCELL activator in the $3^{rd}$ cycle of the T cell stimulation. After cycle 5, we monitored a high-level expression of TOX and therefore treated cells with the selected TOX inhibitors. Fortunately, within 12 days, we found robust T cell re-activation in this method (Fig.4A).

To optimize the concentration of treatment, we tested **KI-TOX-A3, KI-TOX-D22,** and **KI-TOX-P14** in the cytotoxicity test. As shown in figure S6A, we found that all three candidates do not induce cell cytotoxicity at 20uM for day 1, but impact cell growth by days 2-3. Therefore, we chose to treat T cells at 20uM, 10uM, and 5μM for 24h.

We also probed the mRNA levels of the TOX and RUNX3 genes. We found that mRNA levels within the exhausted T cells is about 2-fold higher than the naïve T cells, while **KI-TOX-A3** could downregulate the mRNA level of TOX to the level of naïve T cells. This is expected and was observed in the T-ALL experiments. We hypothesized that this could be a TOX positive feedback loop after TOX degradation (Fig. 3). We also investigated the RUNX3 mRNA levels because TOX was known to be negatively regulated by the synergistic regulation of TOX and GATA3 in CTCL lymphoma, though the cellular and molecular mechanisms of action are not fully understood[7]. In this study, we do not notice any significant changes in RUNX3 and this implies that the TOX mechanism in CD8+ T cell is not identical to the CTCL cells.

Later, we treated T cells with the TOX inhibitors with various concentrations (20, 10, and 5μM). The figure S6C overlaid treated group, exhausted T cell group, and the non-exhausted T cell groups. We noticed that the compound **KI-TOX-A3** exhibited strong dose-dependent effect in inhibitory receptor expression reduction. The LAG-3 and TIM-3 Level were significantly downregulated in 20uM while 10uM and 5μM the TIM-$3^{low}$ population are less. On the other hand, **KI-TOX-A3** shifted PD-1 level dose-dependently as well.

The compound **KI-TOX-D22** and **KI-TOX-P14** candidates showed reduction of inhibitory receptors in 20, 10 and 5μM. However, both candidates do not show a strong pattern of dose-dependent effect, implying that 5μM might be saturated for TOX inhibition and increase the LAG-$3^{low}$TIM-$3^{low}$ population.

7. Compound Characterization:

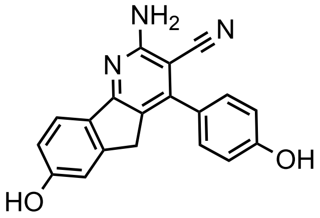

KI-TOX-A3-0186

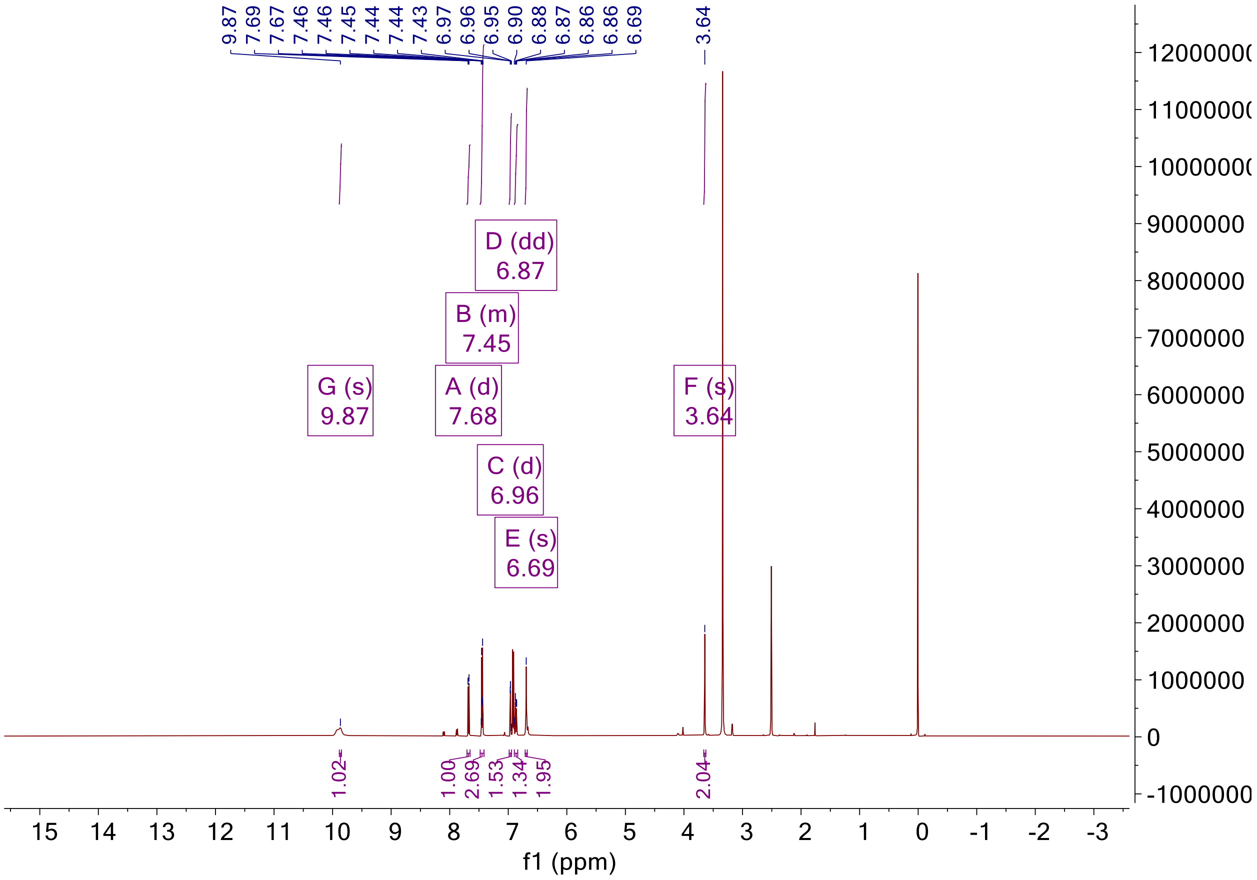

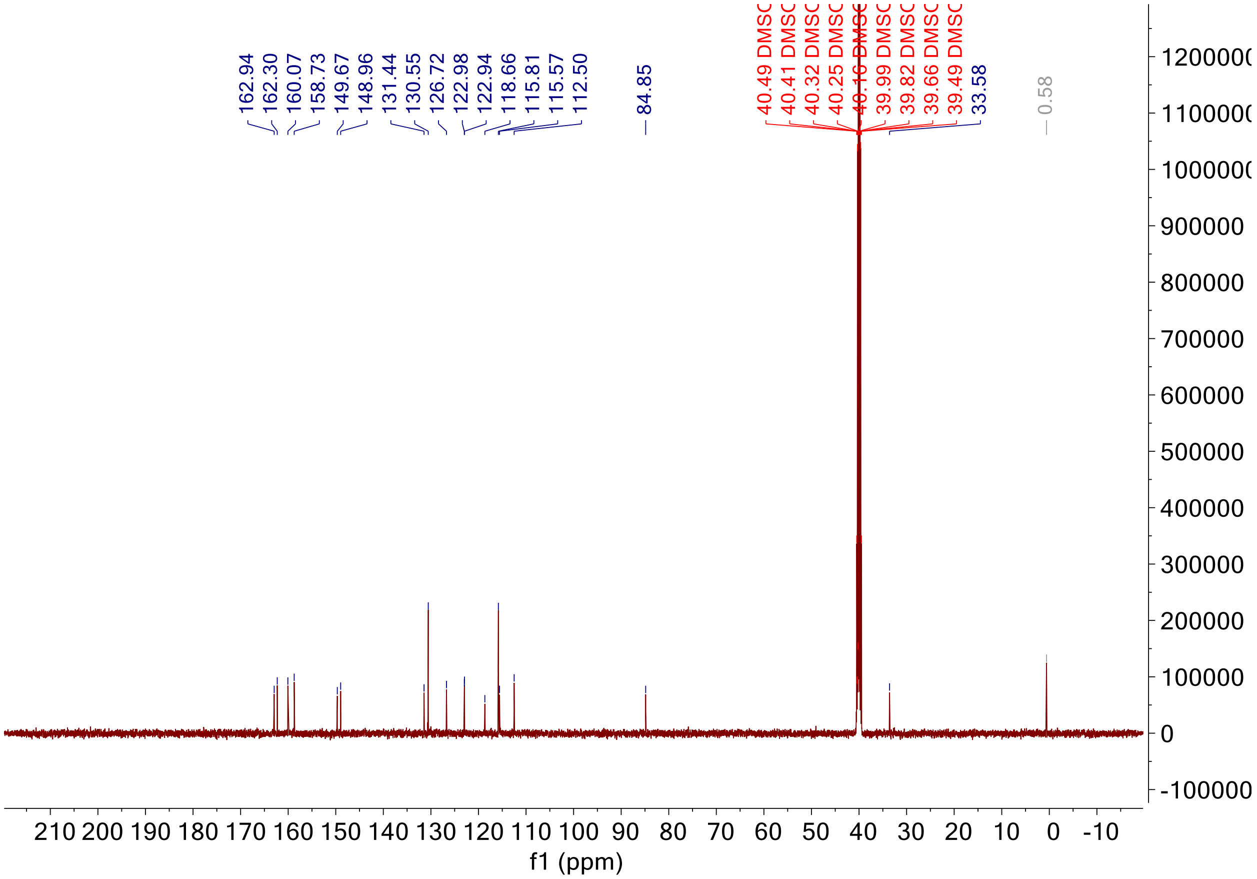

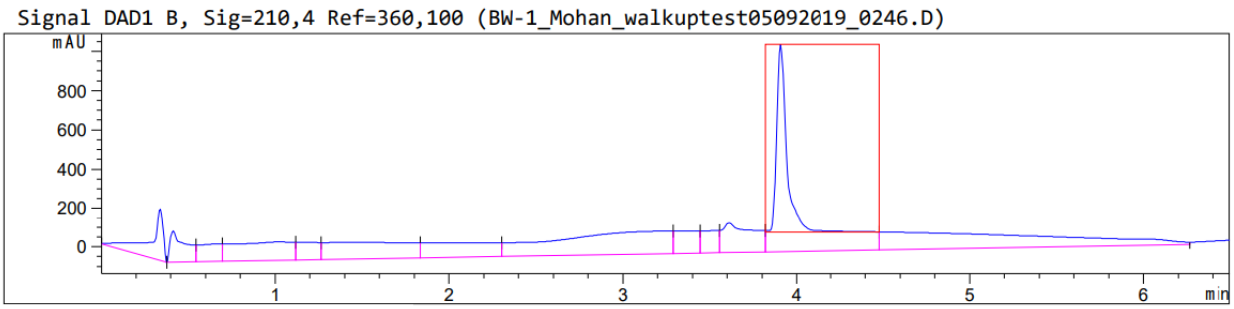

Peak :19 at 3.908 min Name : ?

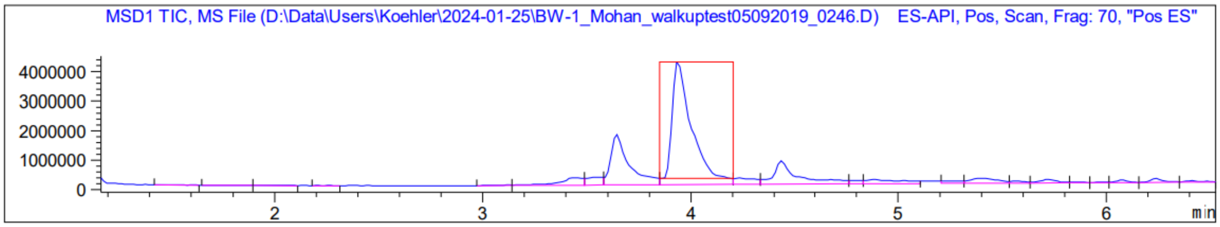
MSD1 TIC, MS File (D:\Data\Users\Koehler\2024-01-25\BW-1_Mohan_walkuptest05092019_0246.D) ES-API, Pos, Scan, Frag: 70, "Pos ES"
4000000
3000000
2000000
1000000
0
2
3
4
5
6
min

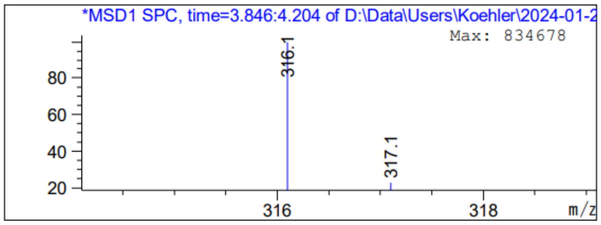
*MSD1 SPC, time=3.846:4.204 of D:\Data\Users\Koehler\2024-01-2
Max: 834678
316.1
317.1
80
60
40
20
316
318
m/z

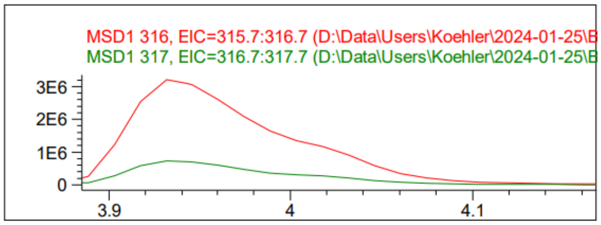
MSD1 316, EIC=315.7:316.7 (D:\Data\Users\Koehler\2024-01-25\B
MSD1 317, EIC=316.7:317.7 (D:\Data\Users\Koehler\2024-01-25\B
3E6
2E6
1E6
0
3.9
4
4.1

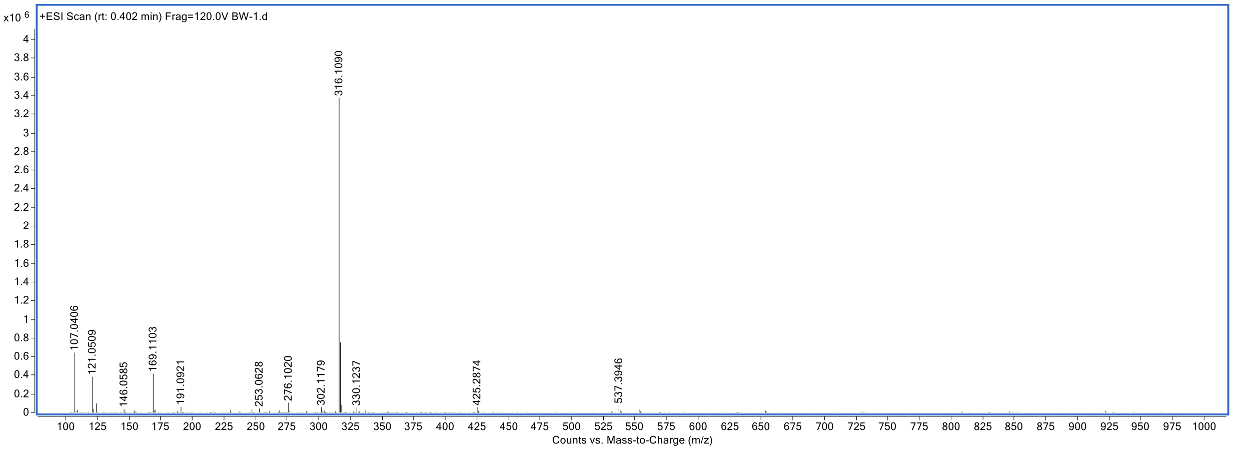
x10 6
+ESI Scan (rt: 0.402 min) Frag=120.0V BW-1.d
316.1090
107.0406
121.0509
146.0585
169.1103
191.0921
253.0628
276.1020
302.1179
330.1237
425.2874
537.3946
Counts vs. Mass-to-Charge (m/z)

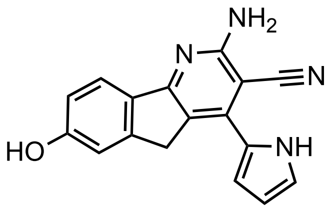

KI-TOX-A3-0185

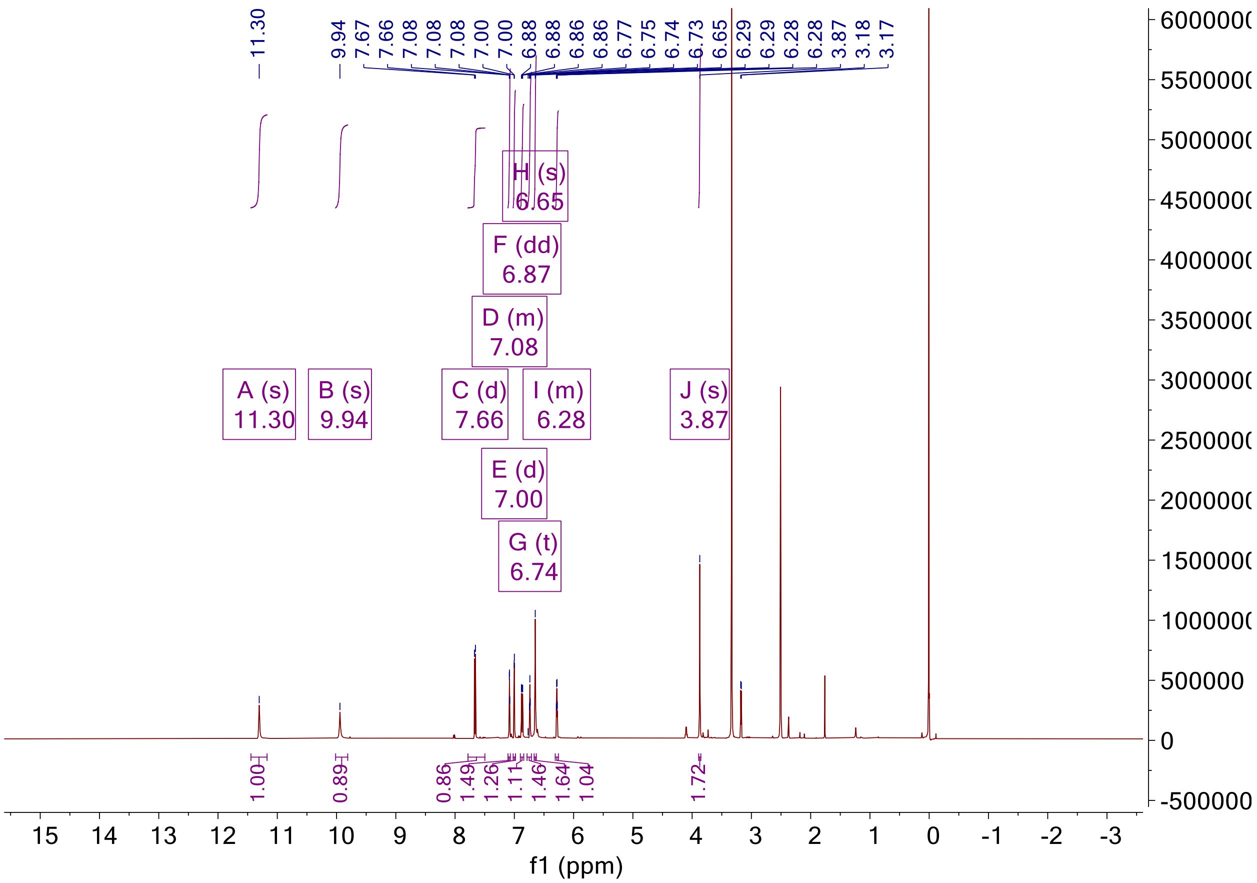

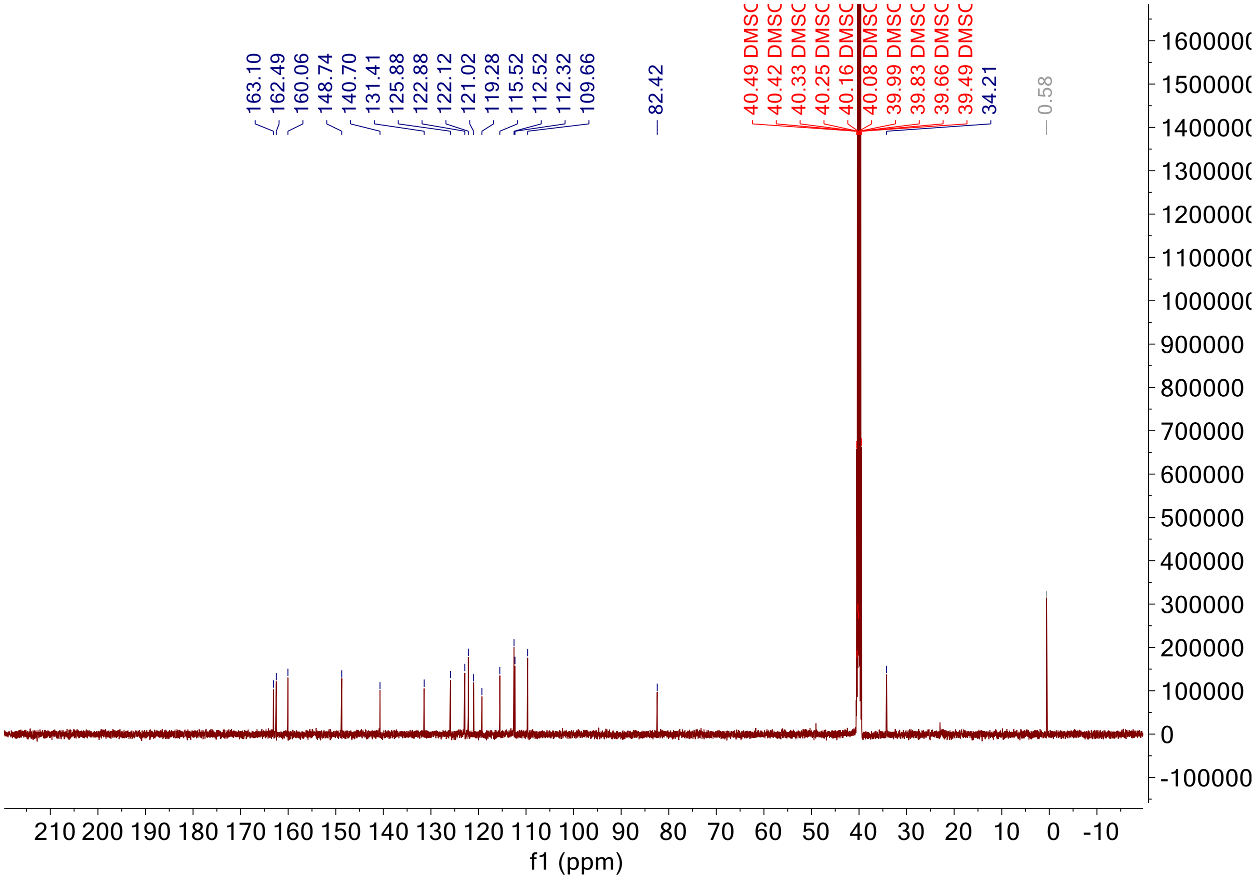

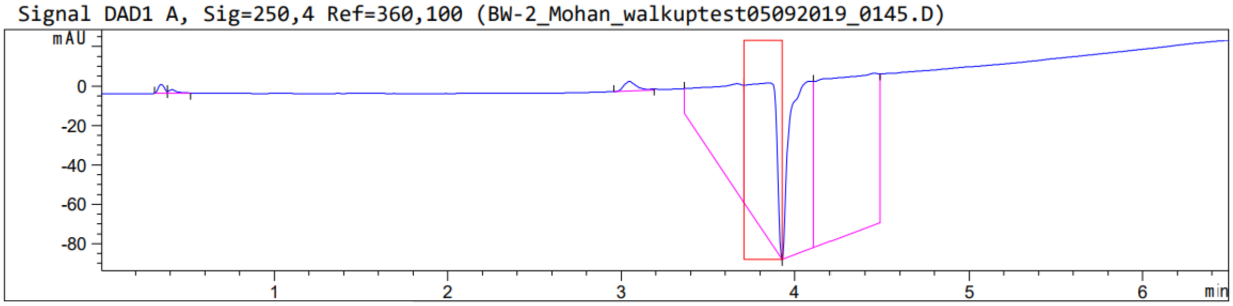

Peak :14 at 3.858 min Name : ?

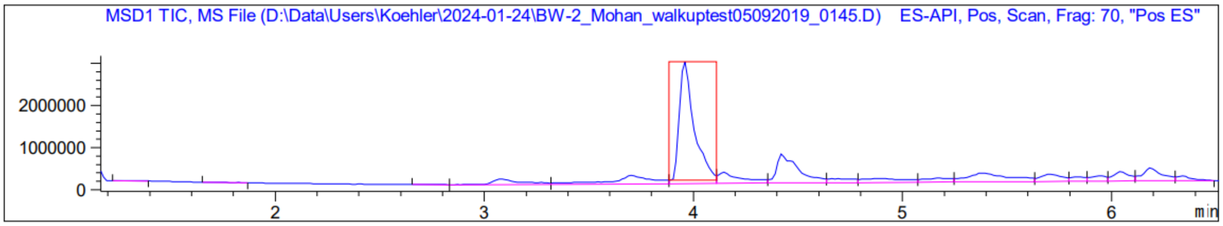
MSD1 TIC, MS File (D:\Data\Users\Koehler\2024-01-24\BW-2_Mohan_walkuptest05092019_0145.D) ES-API, Pos, Scan, Frag: 70, "Pos ES"
2000000
1000000
0
2
3
4
5
6
min

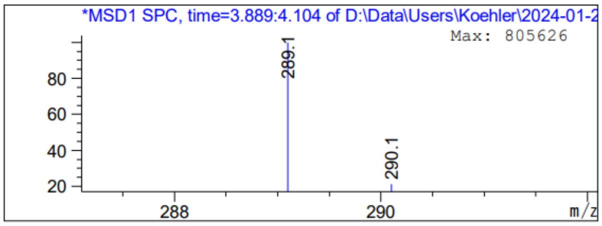
*MSD1 SPC, time=3.889:4.104 of D:\Data\Users\Koehler\2024-01-2
Max: 805626
289.1
290.1
80
60
40
20
288
290
m/z

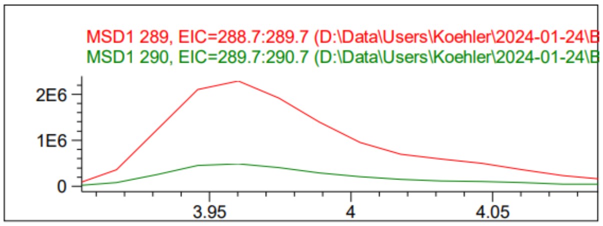
MSD1 289, EIC=288.7:289.7 (D:\Data\Users\Koehler\2024-01-24\B
MSD1 290, EIC=289.7:290.7 (D:\Data\Users\Koehler\2024-01-24\B
2E6
1E6
0
3.95
4
4.05

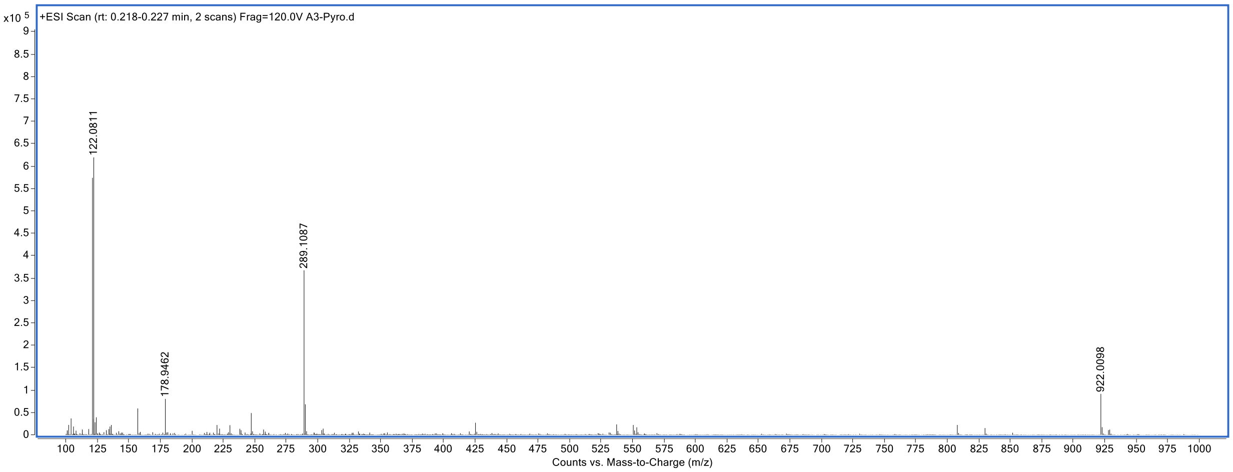
x10 5
+ESI Scan (rt: 0.218-0.227 min, 2 scans) Frag=120.0V A3-Pyro.d
122.0811
178.9462
289.1087
922.0098
Counts vs. Mass-to-Charge (m/z)

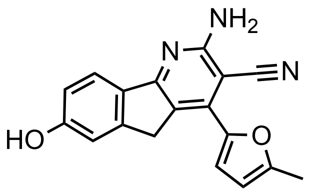

KI-TOX-A3-0181

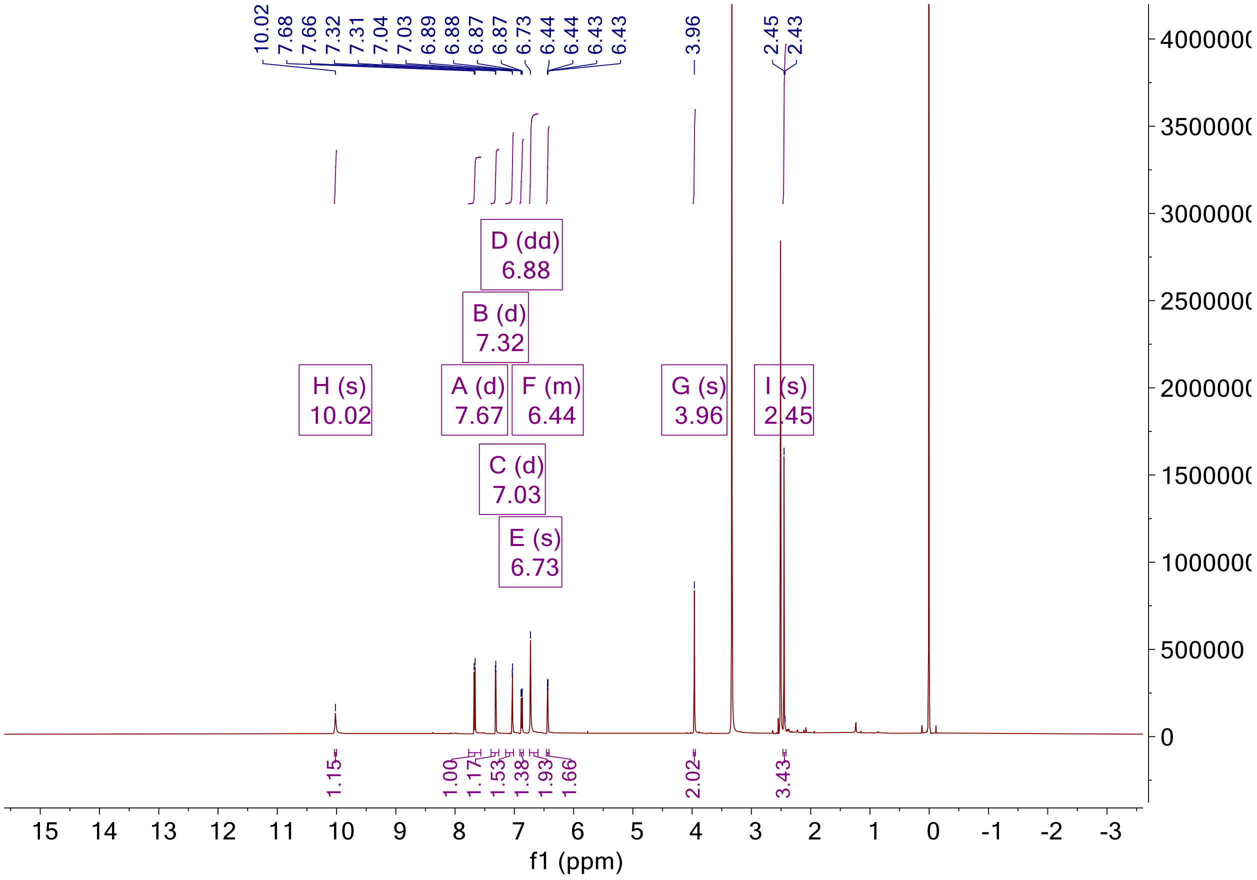

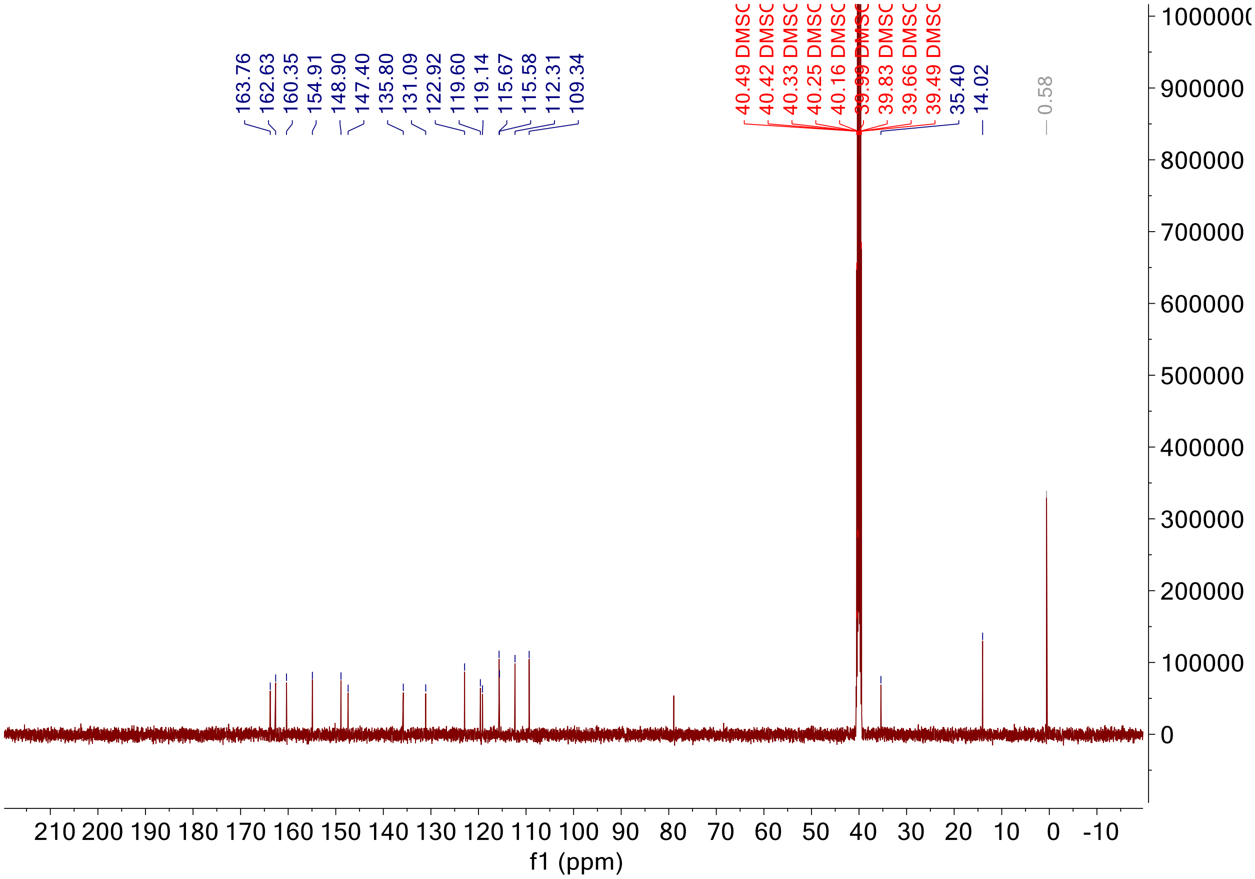

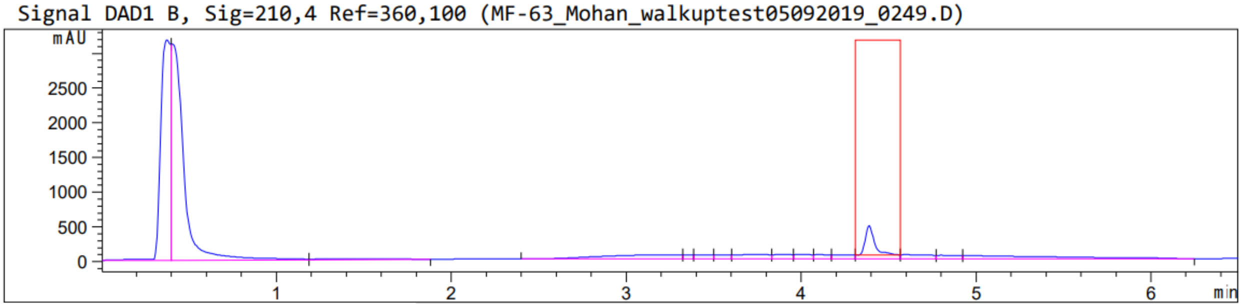

Peak :28 at 4.389 min   Name : ?

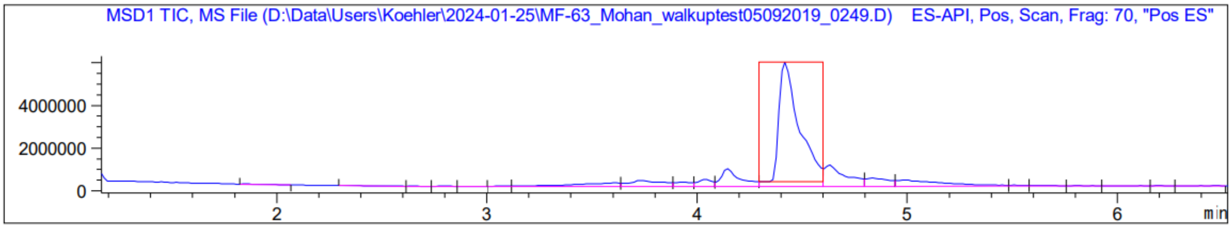
MSD1 TIC, MS File (D:\Data\Users\Koehler\2024-01-25\MF-63_Mohan_walkuptest05092019_0249.D) ES-API, Pos, Scan, Frag: 70, "Pos ES"
4000000
2000000
0
2
3
4
5
6
min

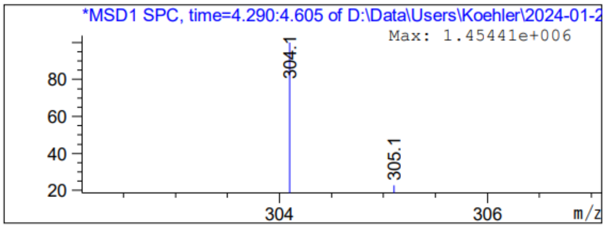
*MSD1 SPC, time=4.290:4.605 of D:\Data\Users\Koehler\2024-01-2
Max: 1.45441e+006
304.1
305.1
80
60
40
20
304
306
m/z

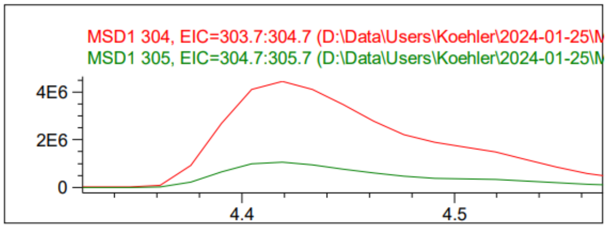
MSD1 304, EIC=303.7:304.7 (D:\Data\Users\Koehler\2024-01-25\M
MSD1 305, EIC=304.7:305.7 (D:\Data\Users\Koehler\2024-01-25\M
4E6
2E6
0
4.4
4.5

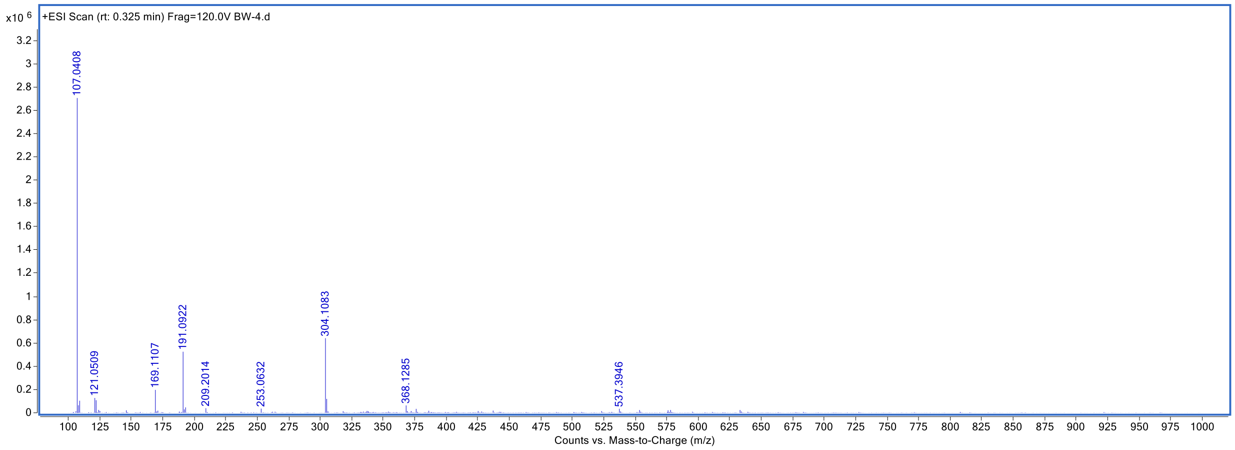
x10 6
+ESI Scan (rt: 0.325 min) Frag=120.0V BW-4.d
107.0408
121.0509
169.1107
191.0922
209.2014
253.0632
304.1083
368.1285
537.3946
Counts vs. Mass-to-Charge (m/z)

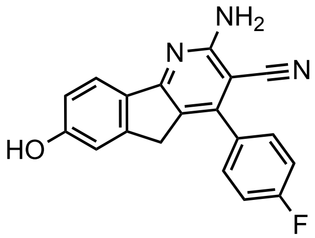

KI-TOX-A3-0180

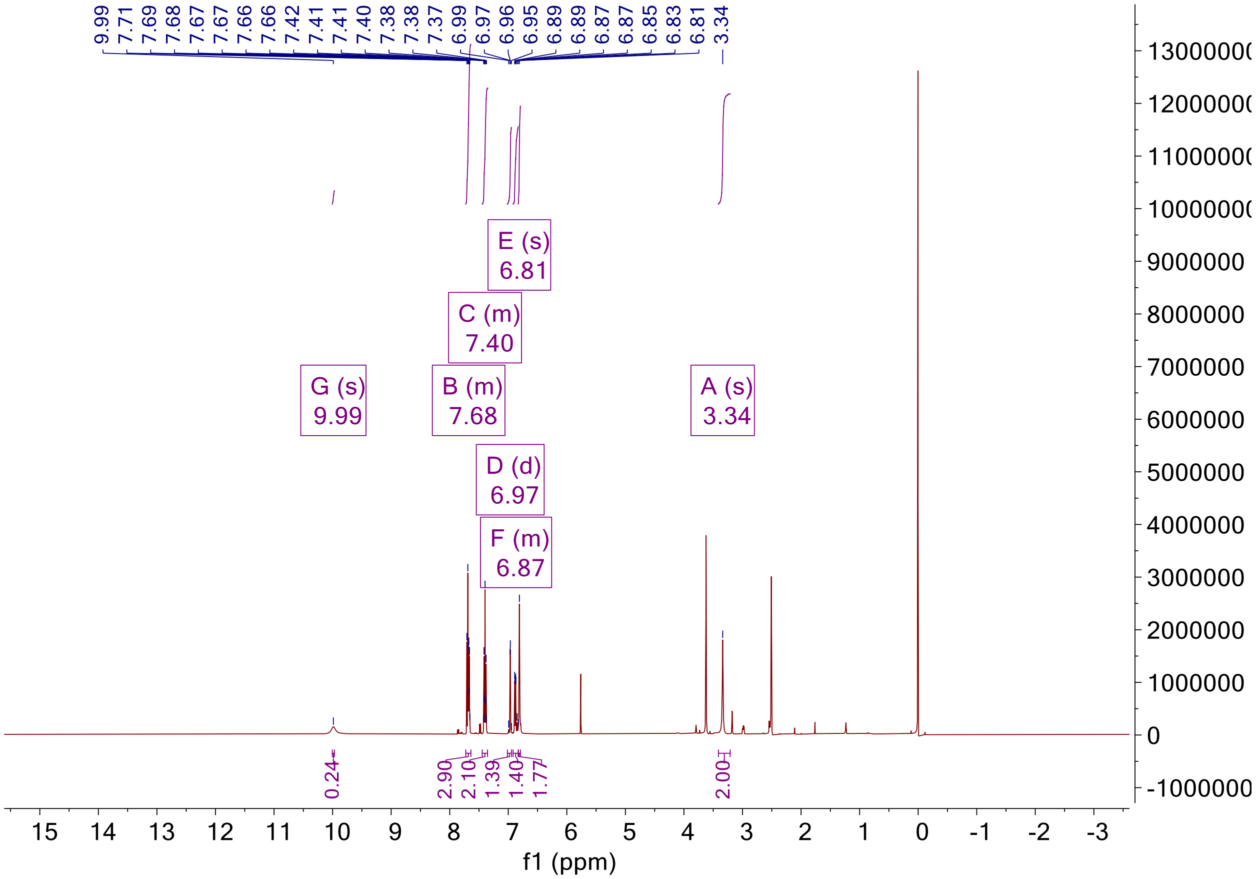

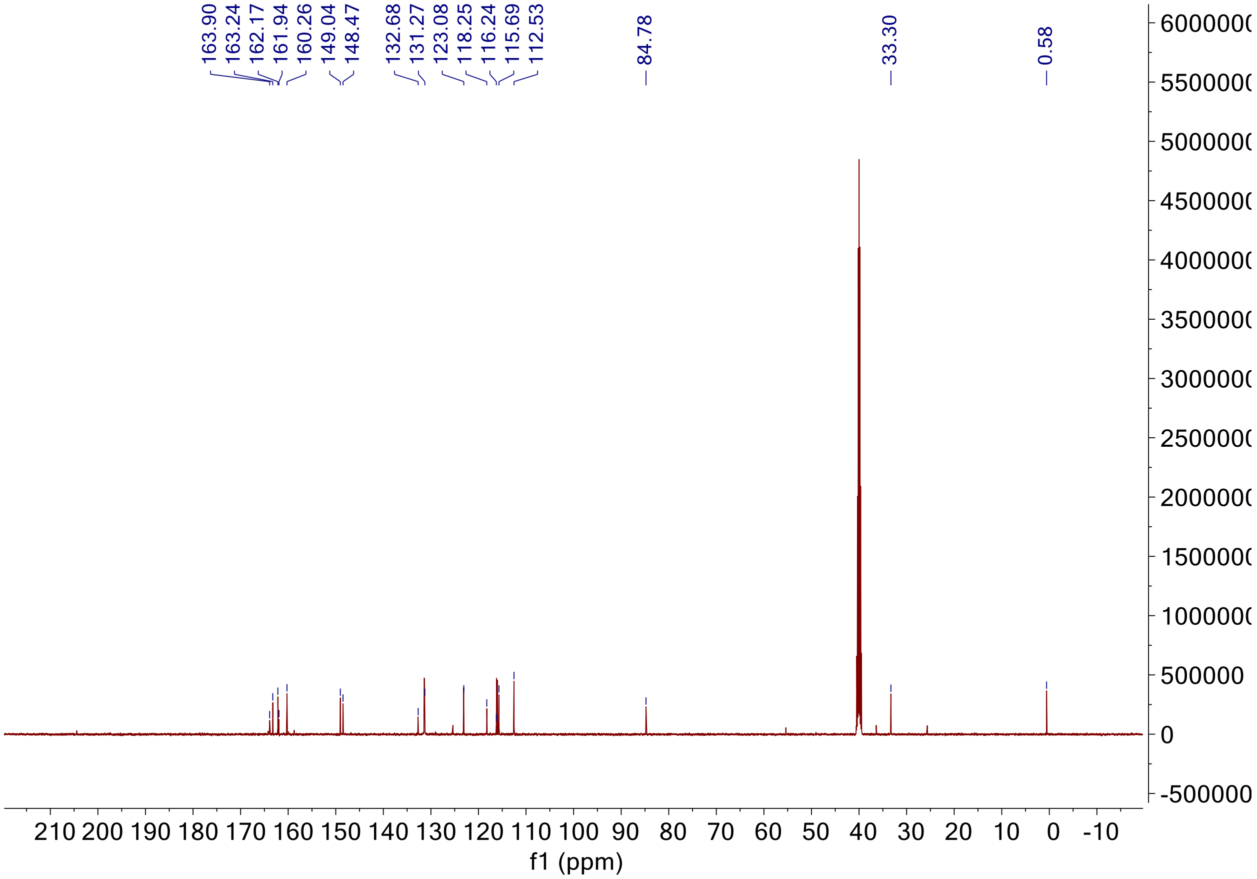

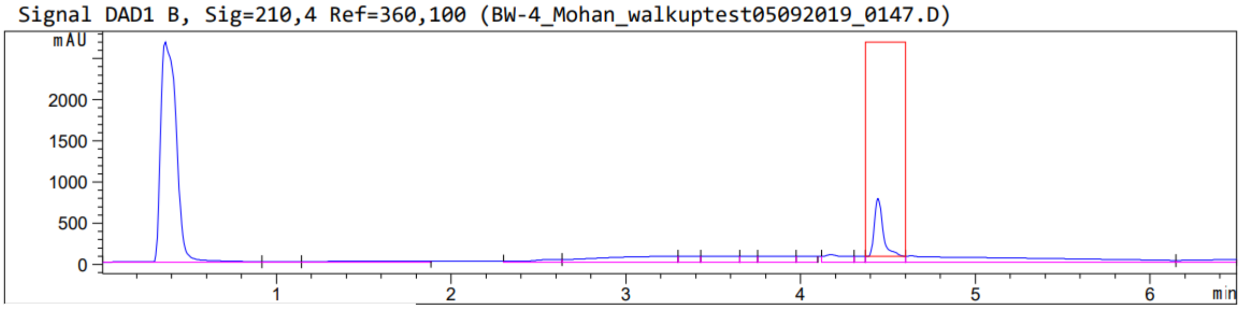

Peak :26 at 4.444 min Name : ?

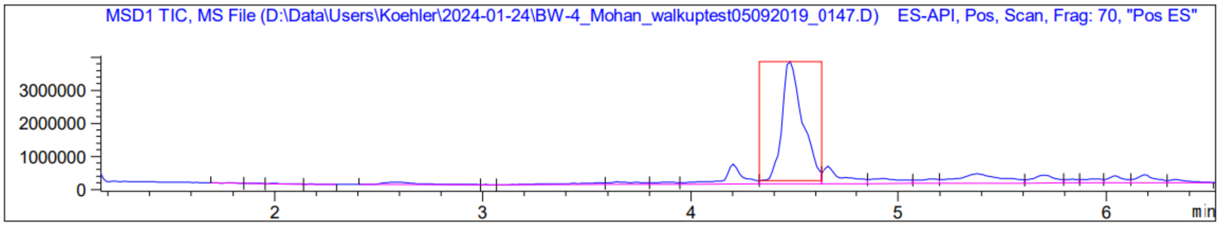
MSD1 TIC, MS File (D:\Data\Users\Koehler\2024-01-24\BW-4_Mohan_walkuptest05092019_0147.D) ES-API, Pos, Scan, Frag: 70, "Pos ES"
3000000
2000000
1000000
0
2
3
4
5
6
min

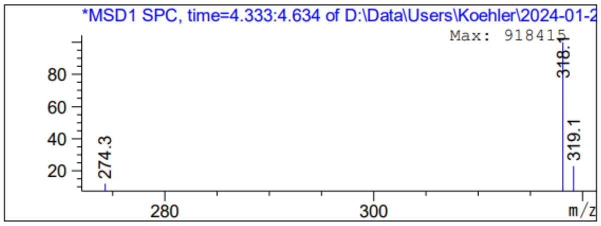
*MSD1 SPC, time=4.333:4.634 of D:\Data\Users\Koehler\2024-01-2
Max: 918415
80
60
40
20
274.3
318.1
319.1
280
300
m/z

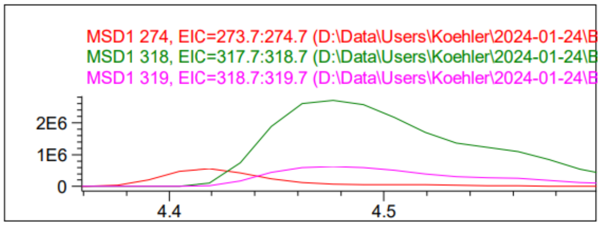
MSD1 274, EIC=273.7:274.7 (D:\Data\Users\Koehler\2024-01-24\B
MSD1 318, EIC=317.7:318.7 (D:\Data\Users\Koehler\2024-01-24\B
MSD1 319, EIC=318.7:319.7 (D:\Data\Users\Koehler\2024-01-24\B
2E6
1E6
0
4.4
4.5

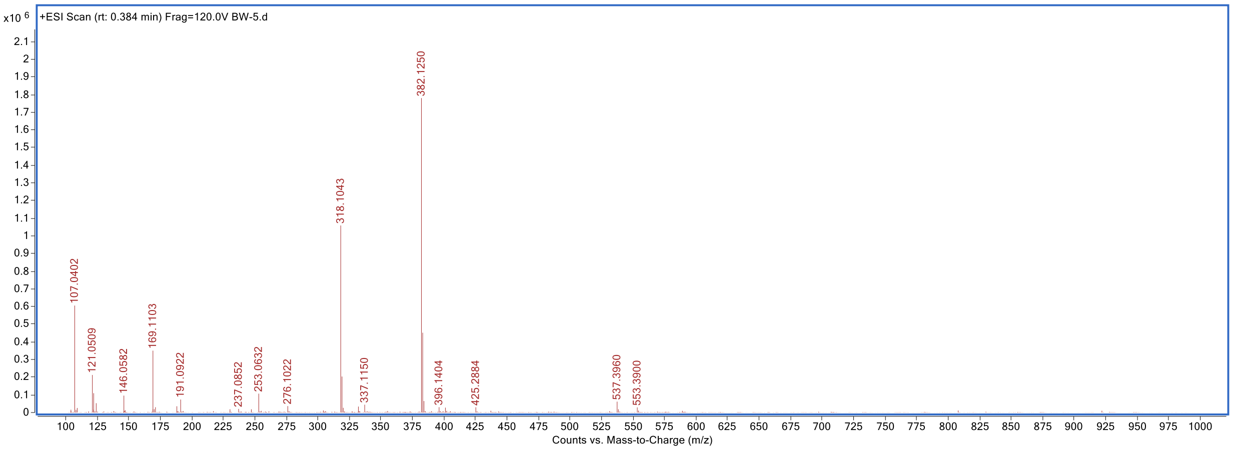
x10 6
+ESI Scan (rt: 0.384 min) Frag=120.0V BW-5.d
107.0402
121.0509
146.0582
169.1103
191.0922
237.0852
253.0632
276.1022
318.1043
337.1150
382.1250
396.1404
425.2884
537.3960
553.3900
Counts vs. Mass-to-Charge (m/z)

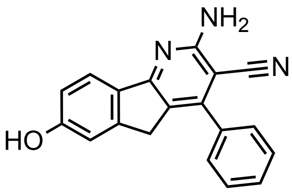

KI-TOX-A3-0184

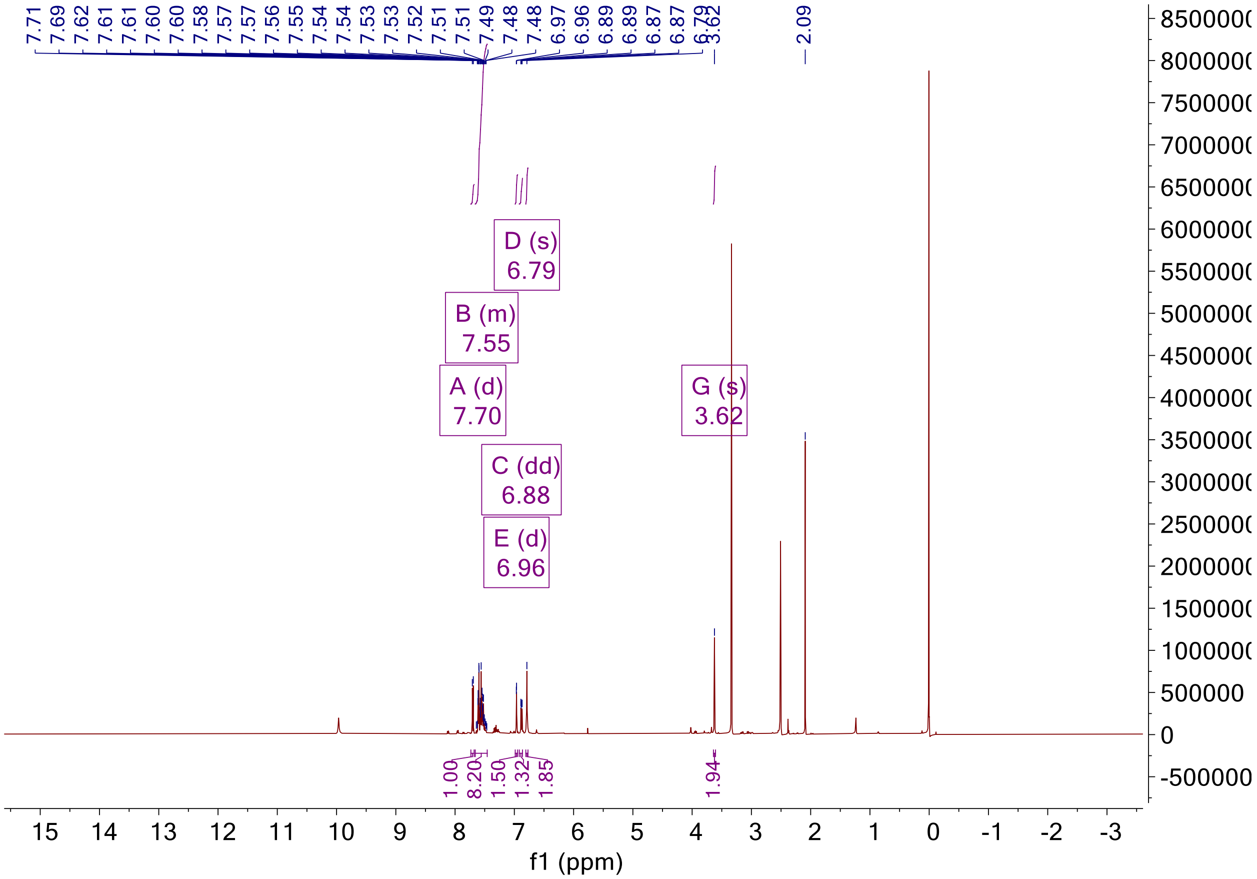

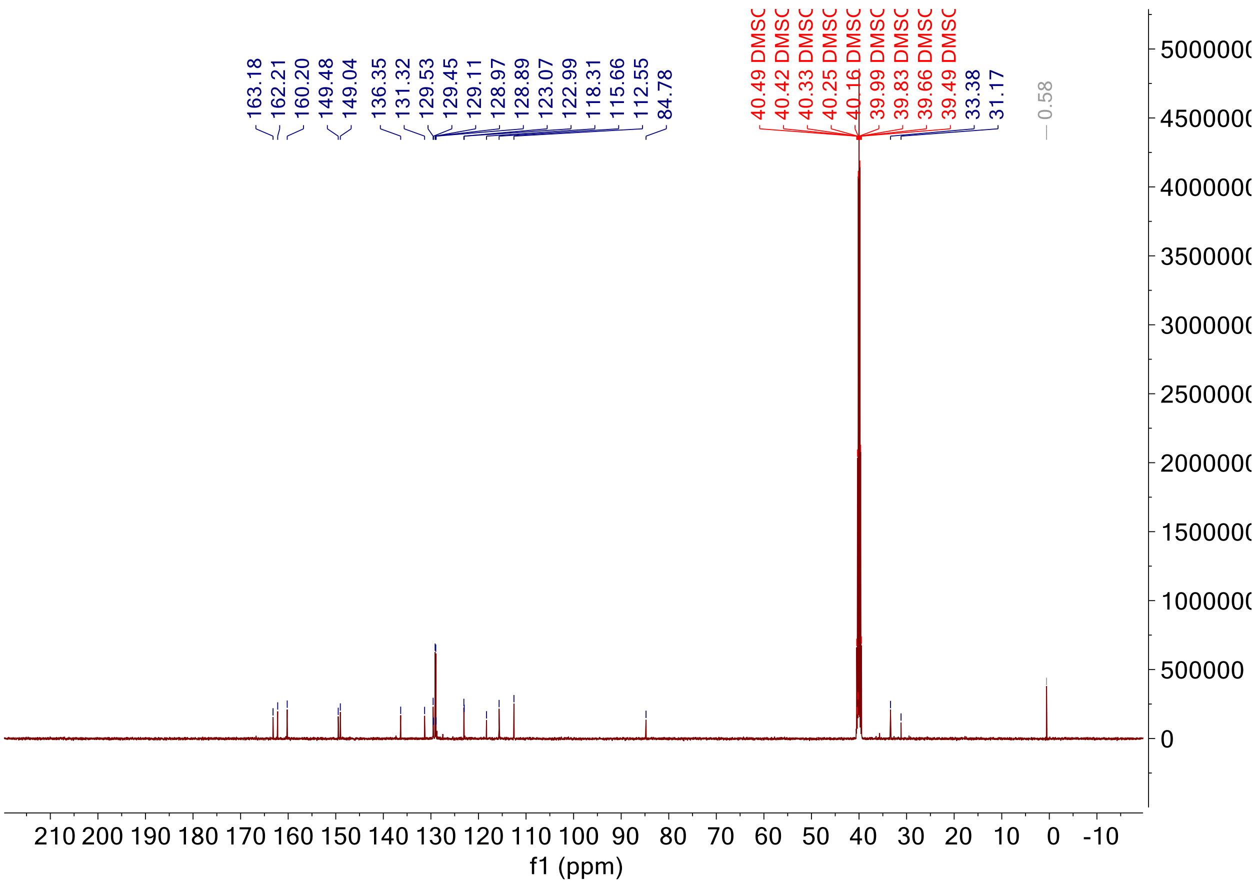

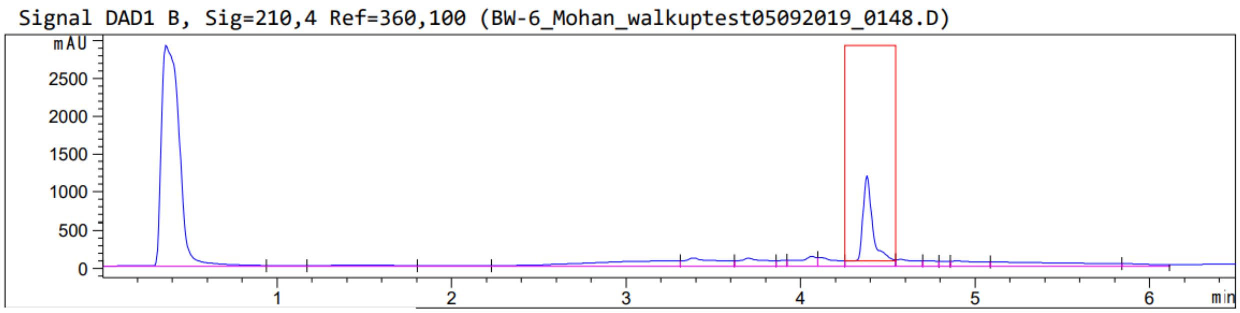

Peak :26 at 4.381 min Name : ?

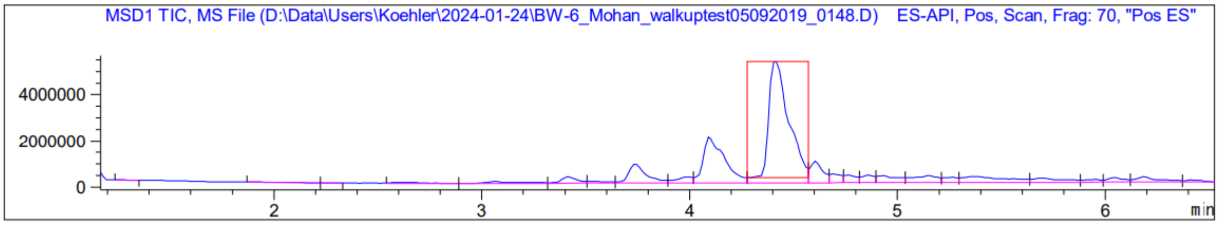
MSD1 TIC, MS File (D:\Data\Users\Koehler\2024-01-24\BW-6_Mohan_walkuptest05092019_0148.D) ES-API, Pos, Scan, Frag: 70, "Pos ES"
4000000
2000000
0
2
3
4
5
6
min

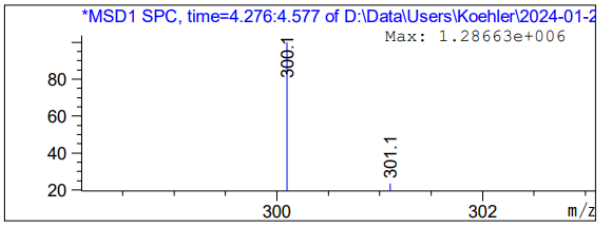
*MSD1 SPC, time=4.276:4.577 of D:\Data\Users\Koehler\2024-01-2
Max: 1.28663e+006
300.1
301.1
80
60
40
20
300
302
m/z

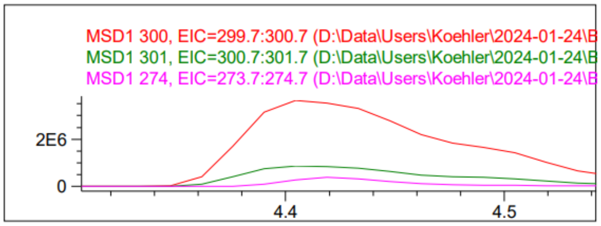
MSD1 300, EIC=299.7:300.7 (D:\Data\Users\Koehler\2024-01-24\B
MSD1 301, EIC=300.7:301.7 (D:\Data\Users\Koehler\2024-01-24\B
MSD1 274, EIC=273.7:274.7 (D:\Data\Users\Koehler\2024-01-24\B
2E6
0
4.4
4.5

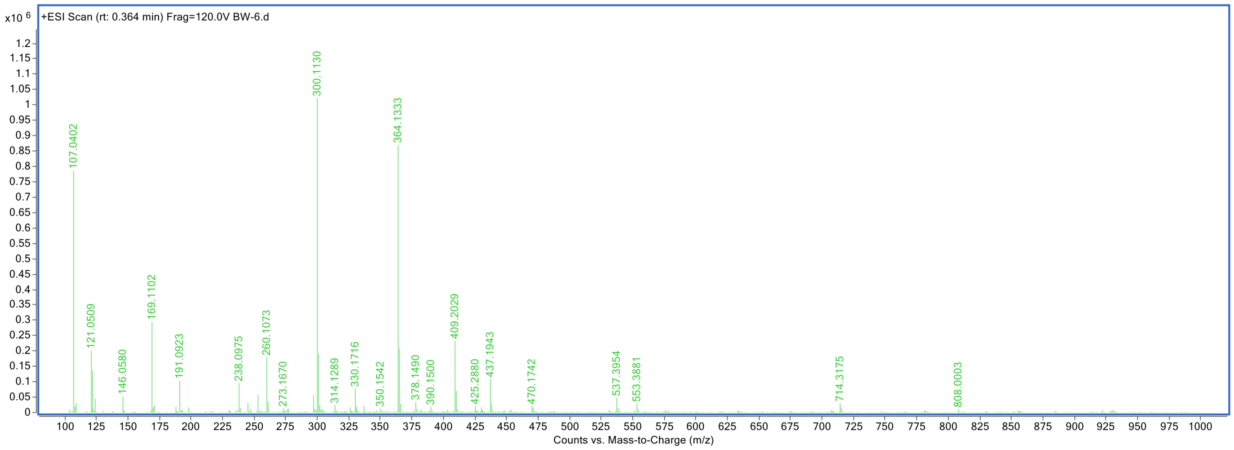
x10 6
+ESI Scan (rt: 0.364 min) Frag=120.0V BW-6.d
107.0402
121.0509
146.0580
169.1102
191.0923
238.0975
260.1073
273.1670
300.1130
314.1289
330.1716
350.1542
364.1333
378.1490
390.1500
409.2029
425.2880
437.1943
470.1742
537.3954
553.3881
714.3175
808.0003
Counts vs. Mass-to-Charge (m/z)

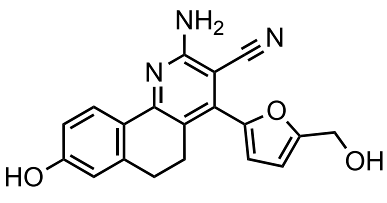

KI-TOX-A3-0176

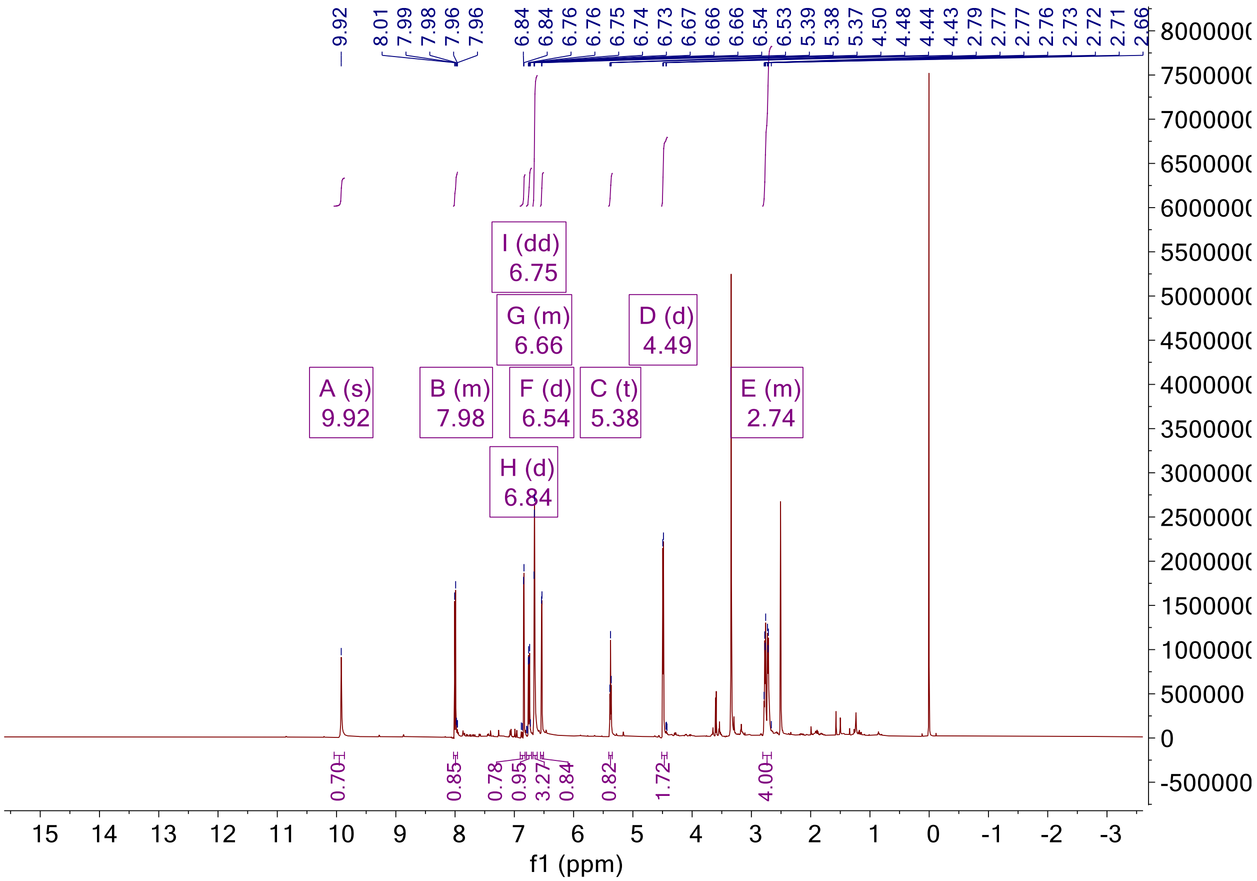

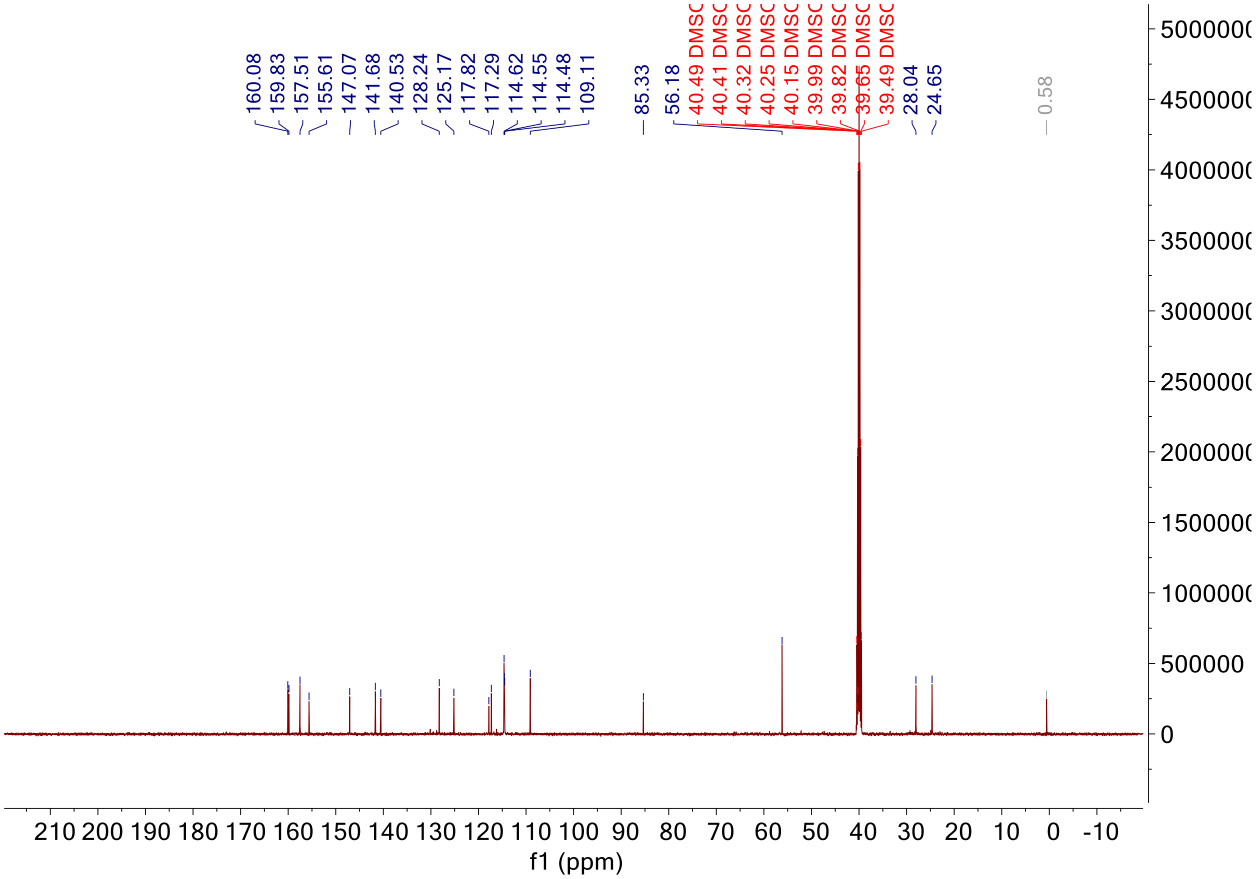

Signal DAD1 A, Sig=250,4 Ref=360,100 (BW-8_Mohan_walkuptest05092019_0150.D)

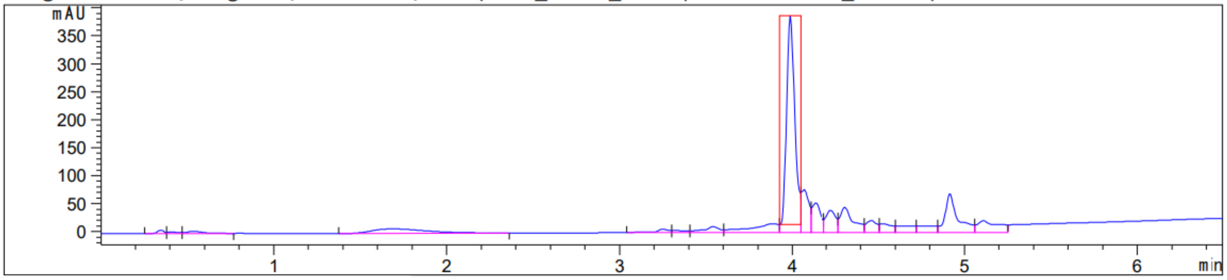

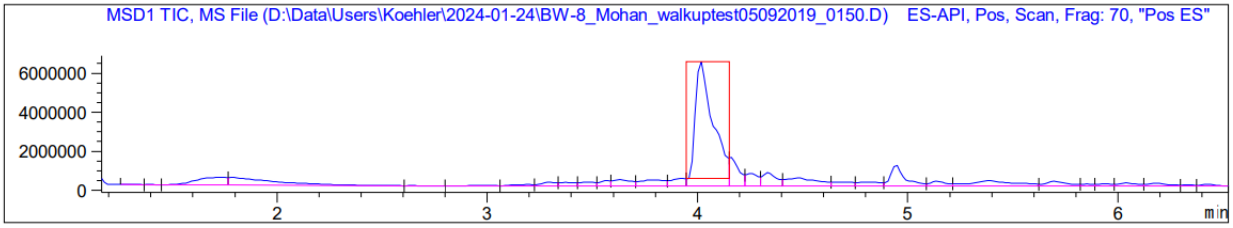
MSD1 TIC, MS File (D:\Data\Users\Koehler\2024-01-24\BW-8_Mohan_walkuptest05092019_0150.D) ES-API, Pos, Scan, Frag: 70, "Pos ES"
6000000
4000000
2000000
0
2
3
4
5
6
min

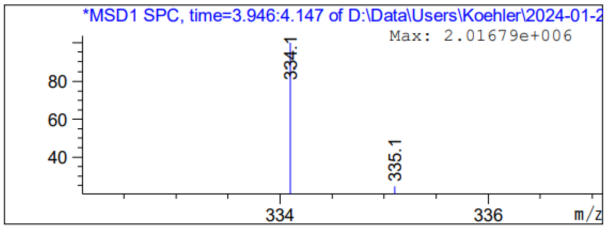
*MSD1 SPC, time=3.946:4.147 of D:\Data\Users\Koehler\2024-01-2
Max: 2.01679e+006
80
60
40
334.1
335.1
334
336
m/z

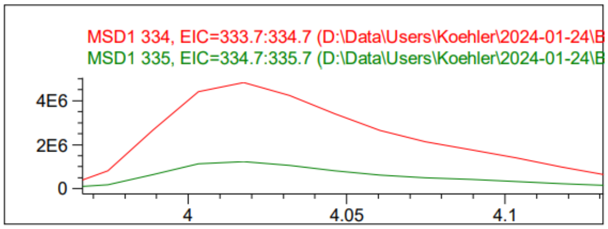
MSD1 334, EIC=333.7:334.7 (D:\Data\Users\Koehler\2024-01-24\B
MSD1 335, EIC=334.7:335.7 (D:\Data\Users\Koehler\2024-01-24\B
4E6
2E6
0
4
4.05
4.1

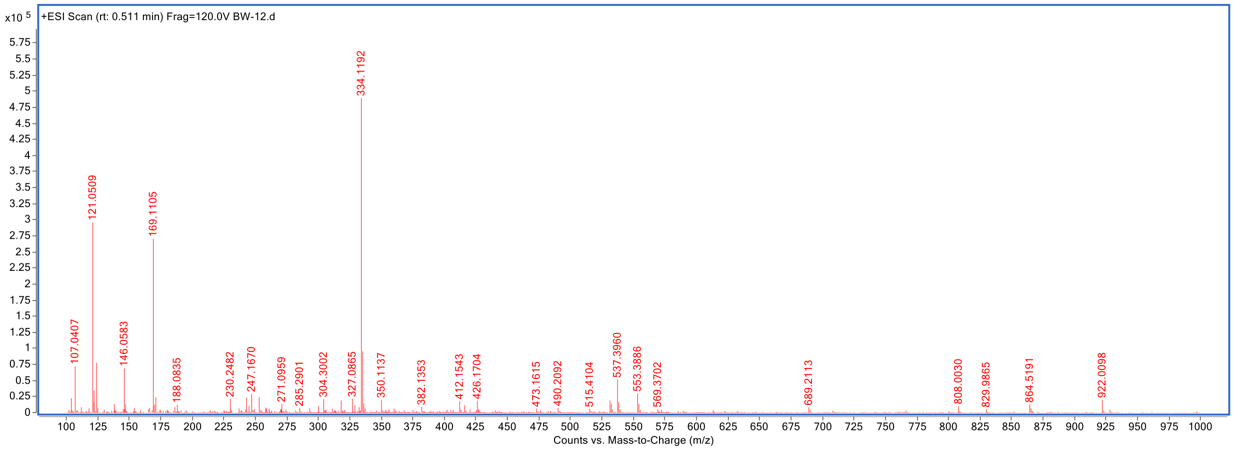
x10 5
+ESI Scan (rt: 0.511 min) Frag=120.0V BW-12.d
334.1192
121.0509
169.1105
107.0407
146.0583
188.0835
230.2482
247.1670
271.0959
285.2901
304.3002
327.0865
350.1137
382.1353
412.1543
426.1704
473.1615
490.2092
515.4104
537.3960
553.3886
569.3702
689.2113
808.0030
829.9865
864.5191
922.0098
Counts vs. Mass-to-Charge (m/z)

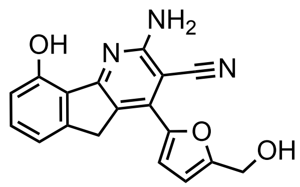

KI-TOX-A3-0177

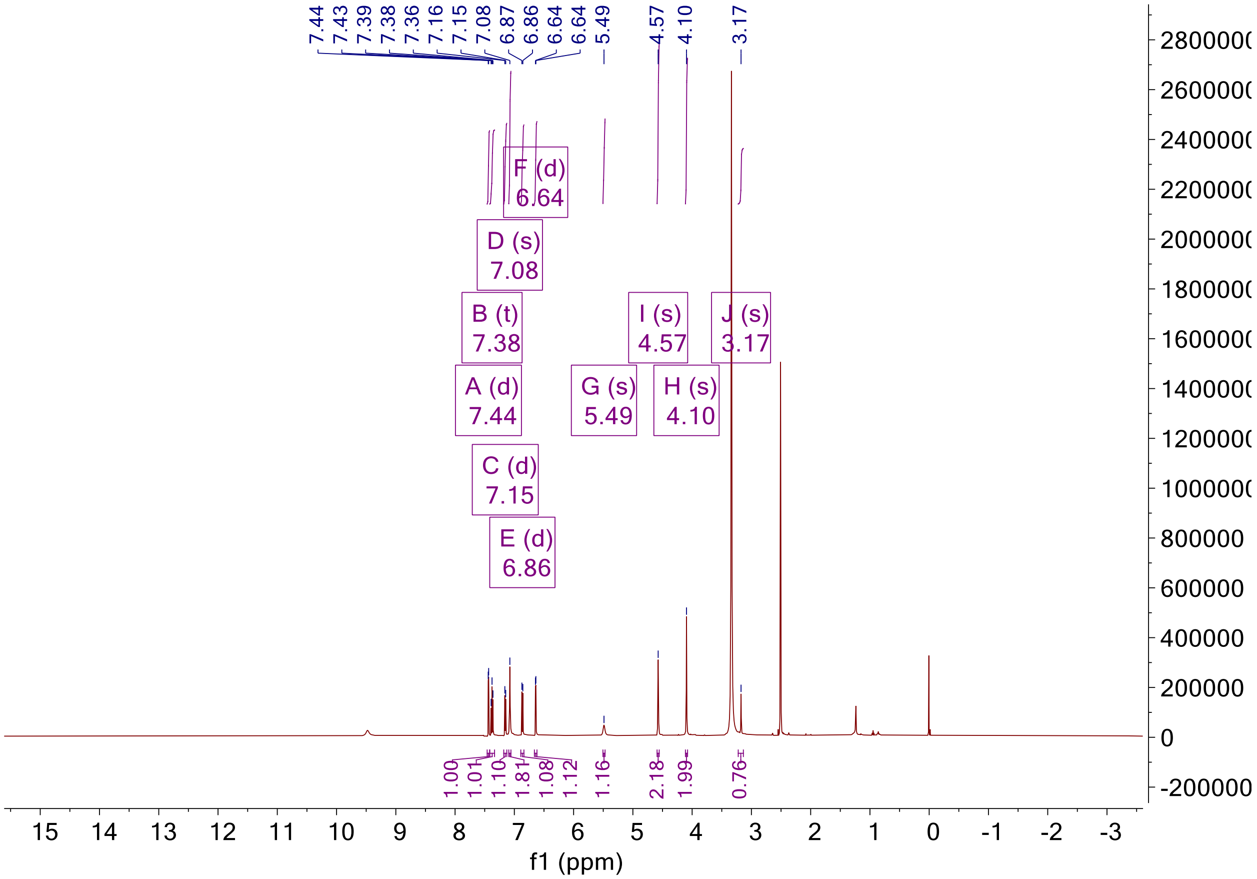

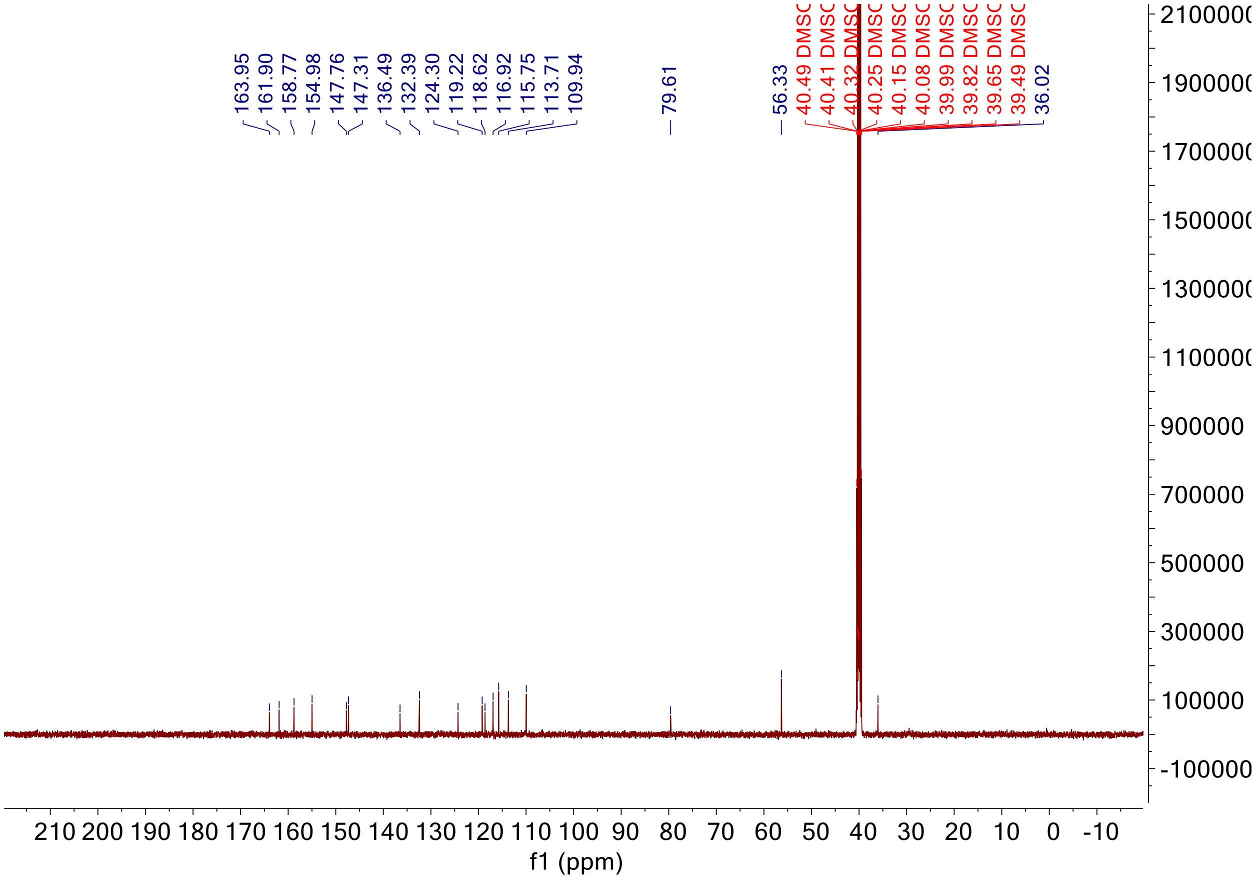

Signal DAD1 G, Sig=280,4 Ref=360,100 (BW-10_Mohan_walkuptest05092019_0152.D)

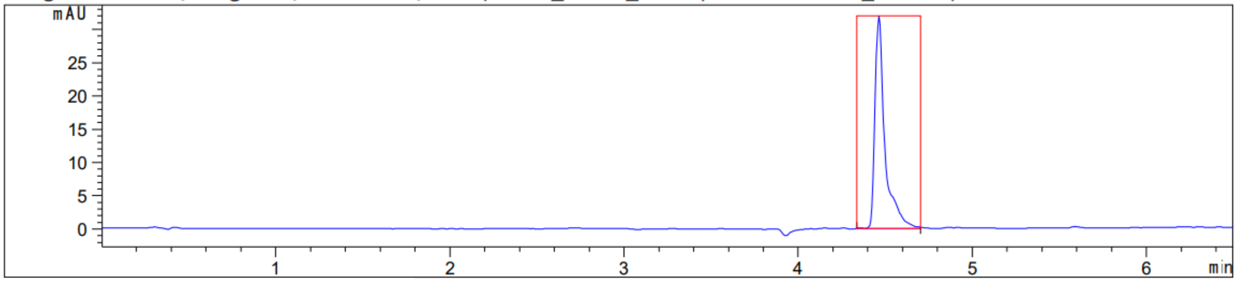

Peak :27 at 4.463 min Name : ?

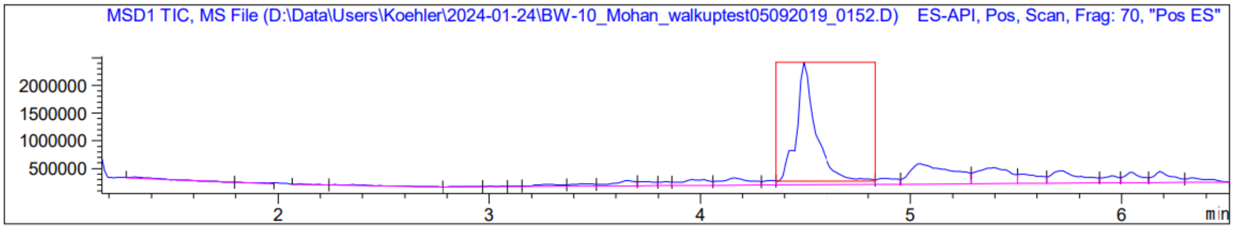
MSD1 TIC, MS File (D:\Data\Users\Koehler\2024-01-24\BW-10_Mohan_walkuptest05092019_0152.D) ES-API, Pos, Scan, Frag: 70, "Pos ES"
2000000
1500000
1000000
500000
2
3
4
5
6
min

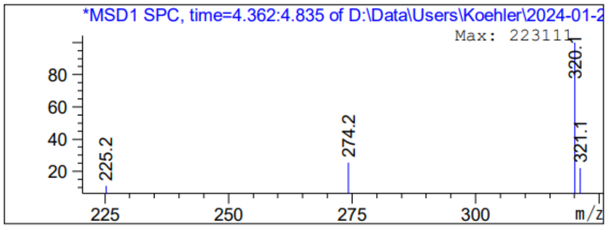
*MSD1 SPC, time=4.362:4.835 of D:\Data\Users\Koehler\2024-01-2
Max: 223111
80
60
40
20
225.2
274.2
320.1
321.1
225
250
275
300
m/z

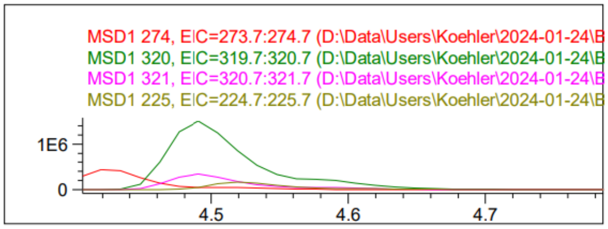
MSD1 274, EIC=273.7:274.7 (D:\Data\Users\Koehler\2024-01-24\B
MSD1 320, EIC=319.7:320.7 (D:\Data\Users\Koehler\2024-01-24\B
MSD1 321, EIC=320.7:321.7 (D:\Data\Users\Koehler\2024-01-24\B
MSD1 225, EIC=224.7:225.7 (D:\Data\Users\Koehler\2024-01-24\B
1E6
0
4.5
4.6
4.7

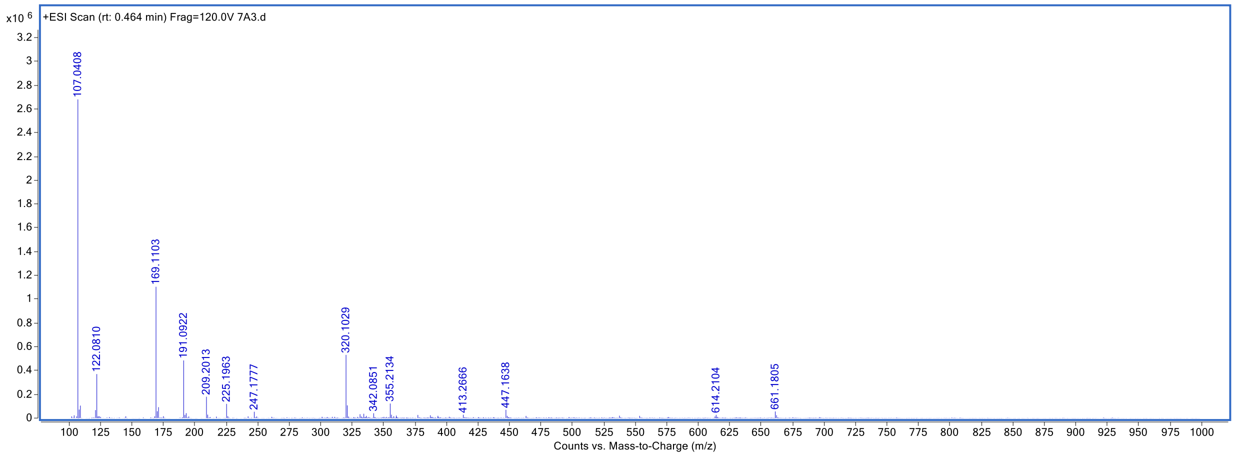
x10 6
+ESI Scan (rt: 0.464 min) Frag=120.0V 7A3.d
107.0408
122.0810
169.1103
191.0922
209.2013
225.1963
247.1777
320.1029
342.0851
355.2134
413.2666
447.1638
614.2104
661.1805
Counts vs. Mass-to-Charge (m/z)

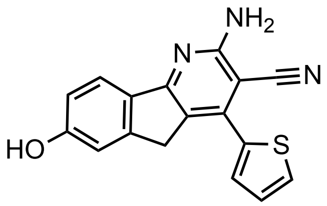

KI-TOX-A3-0182

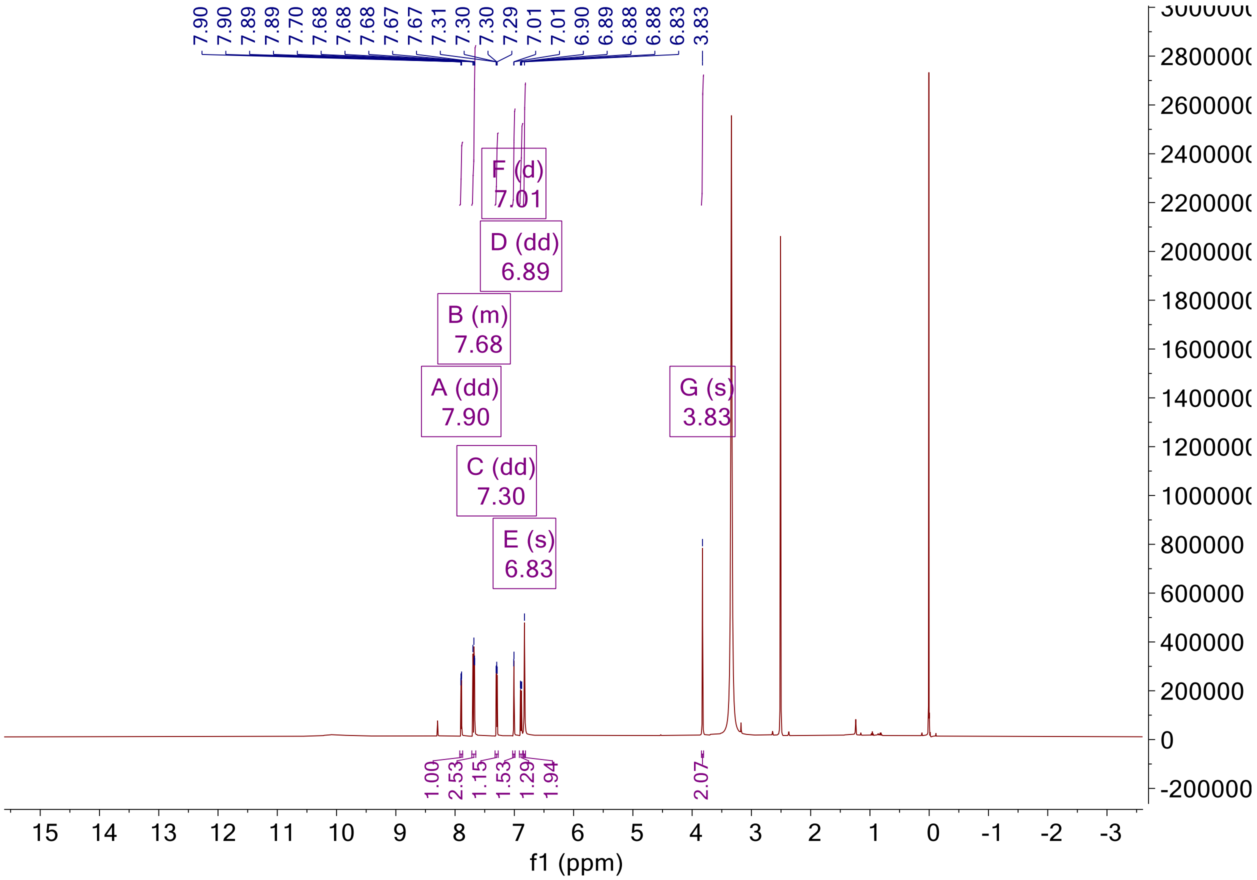

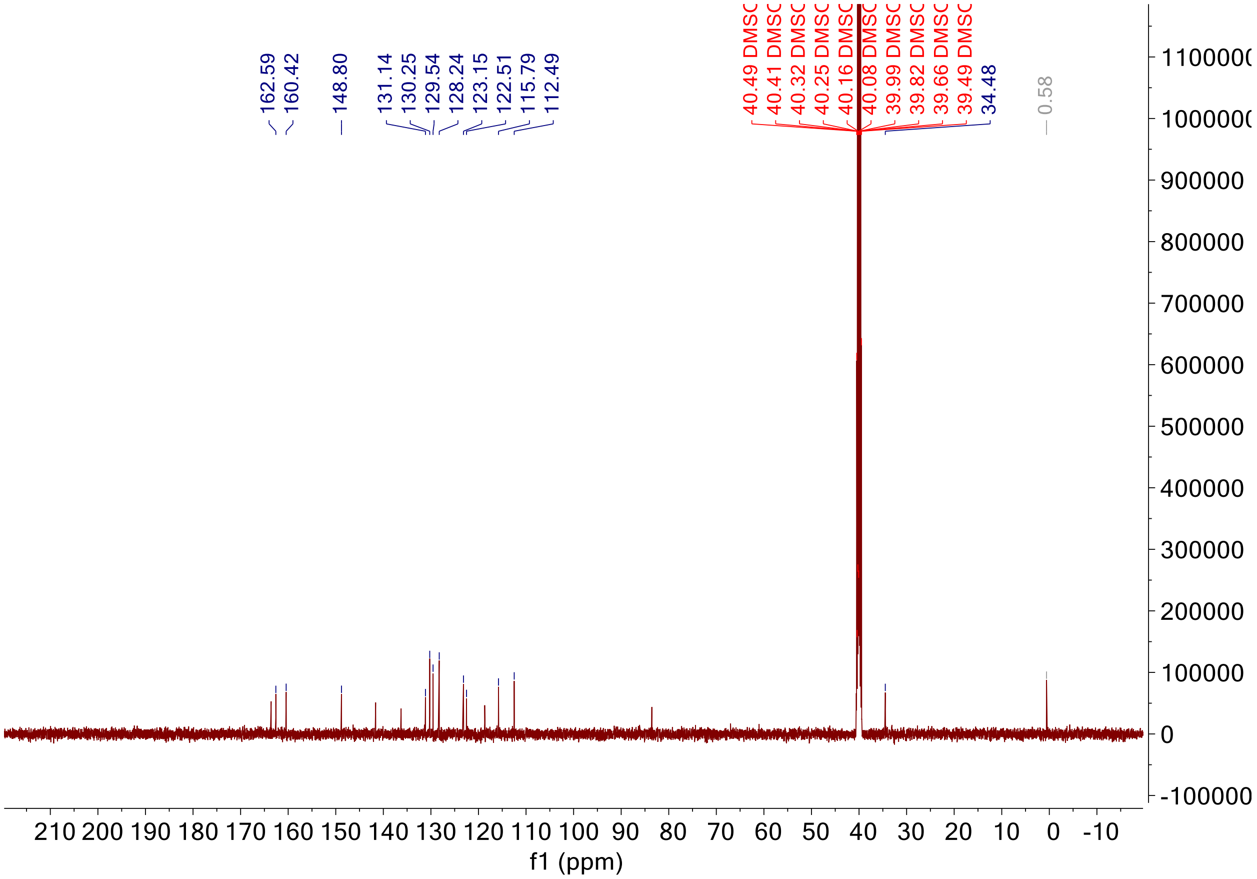

Signal DAD1 B, Sig=210,4 Ref=360,100 (bw-11_Mohan_walkuptest05092019_0332.D)

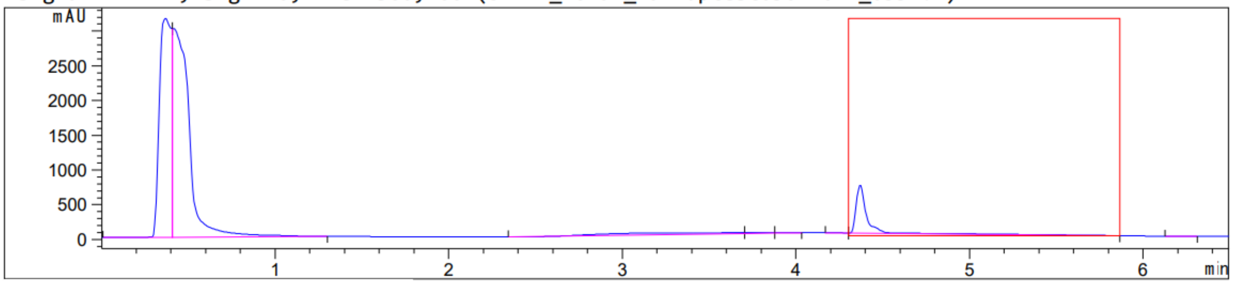

Peak :18 at 4.370 min Name : ?

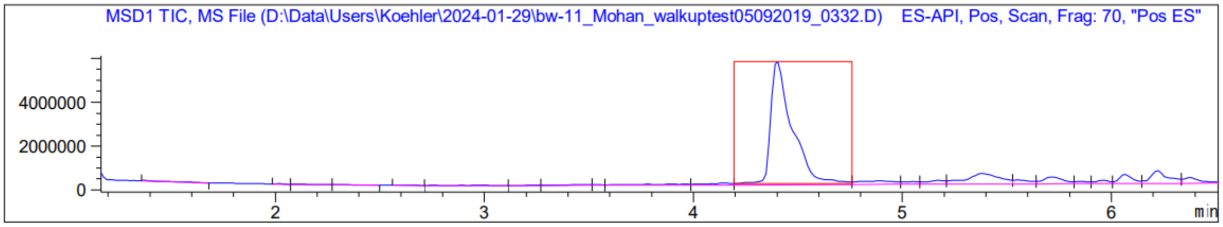
MSD1 TIC, MS File (D:\Data\Users\Koehler\2024-01-29\bw-11_Mohan_walkuptest05092019_0332.D) ES-API, Pos, Scan, Frag: 70, "Pos ES"
4000000
2000000
0
2
3
4
5
6
min

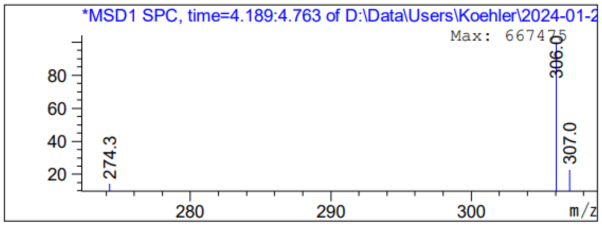
*MSD1 SPC, time=4.189:4.763 of D:\Data\Users\Koehler\2024-01-2
Max: 667475
306.0
307.0
274.3
80
60
40
20
280
290
300
m/z

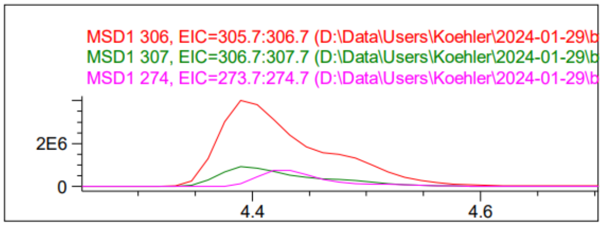
MSD1 306, EIC=305.7:306.7 (D:\Data\Users\Koehler\2024-01-29\b
MSD1 307, EIC=306.7:307.7 (D:\Data\Users\Koehler\2024-01-29\b
MSD1 274, EIC=273.7:274.7 (D:\Data\Users\Koehler\2024-01-29\b
2E6
0
4.4
4.6

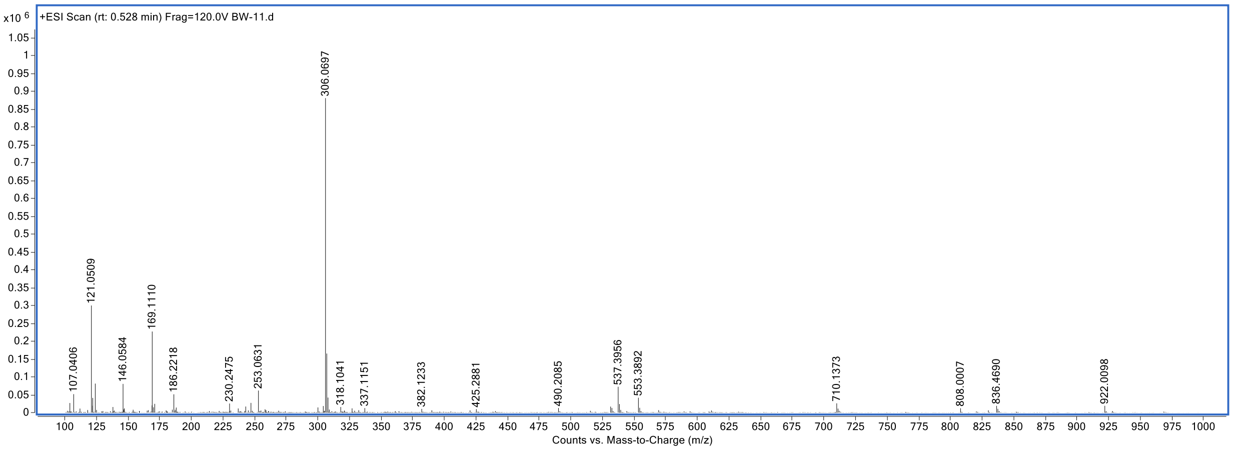
x10 6
+ESI Scan (rt: 0.528 min) Frag=120.0V BW-11.d
306.0697
121.0509
169.1110
107.0406
146.0584
186.2218
230.2475
253.0631
318.1041
337.1151
382.1233
425.2881
490.2085
537.3956
553.3892
710.1373
808.0007
836.4690
922.0098
Counts vs. Mass-to-Charge (m/z)

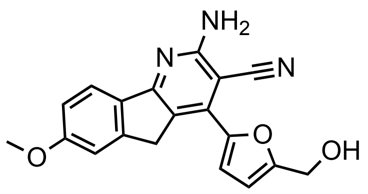

KI-TOX-A3-0174

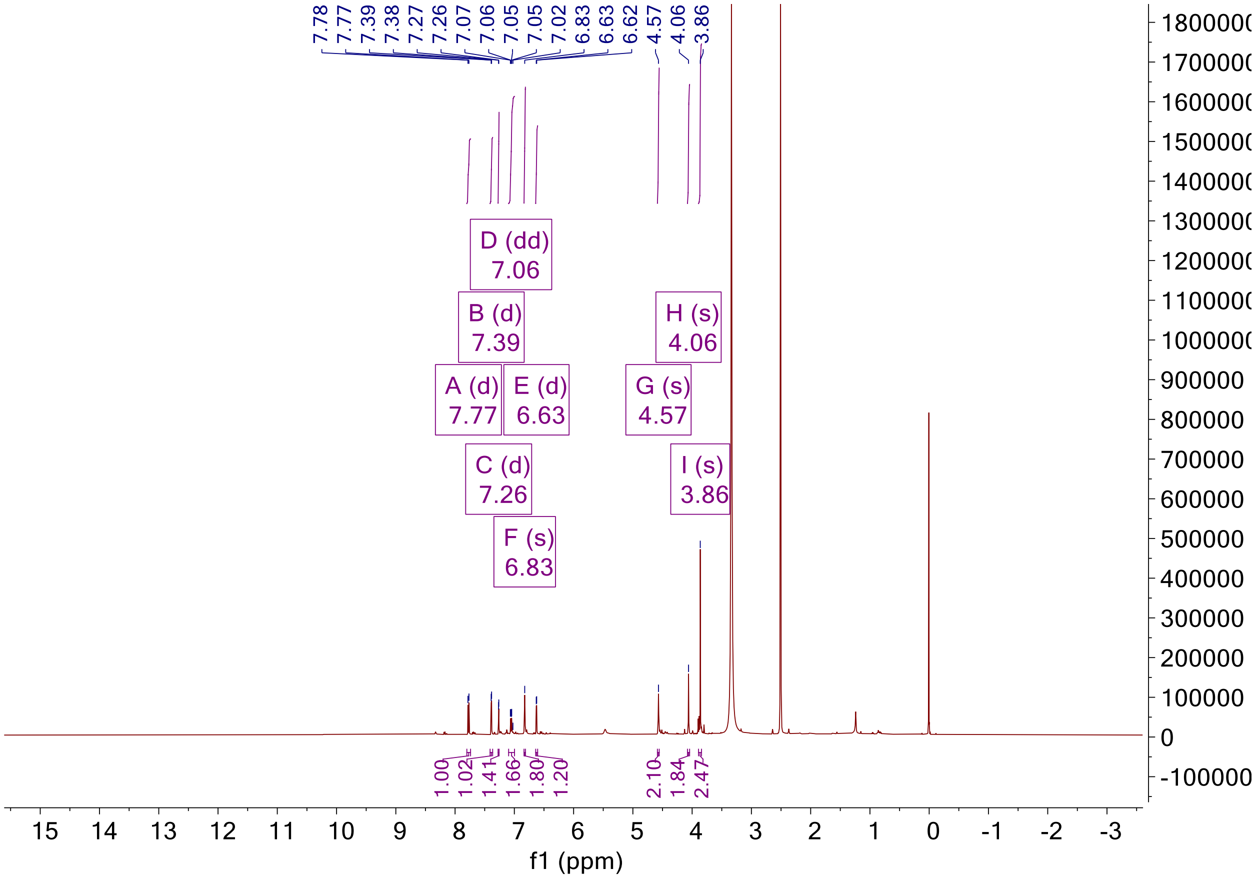

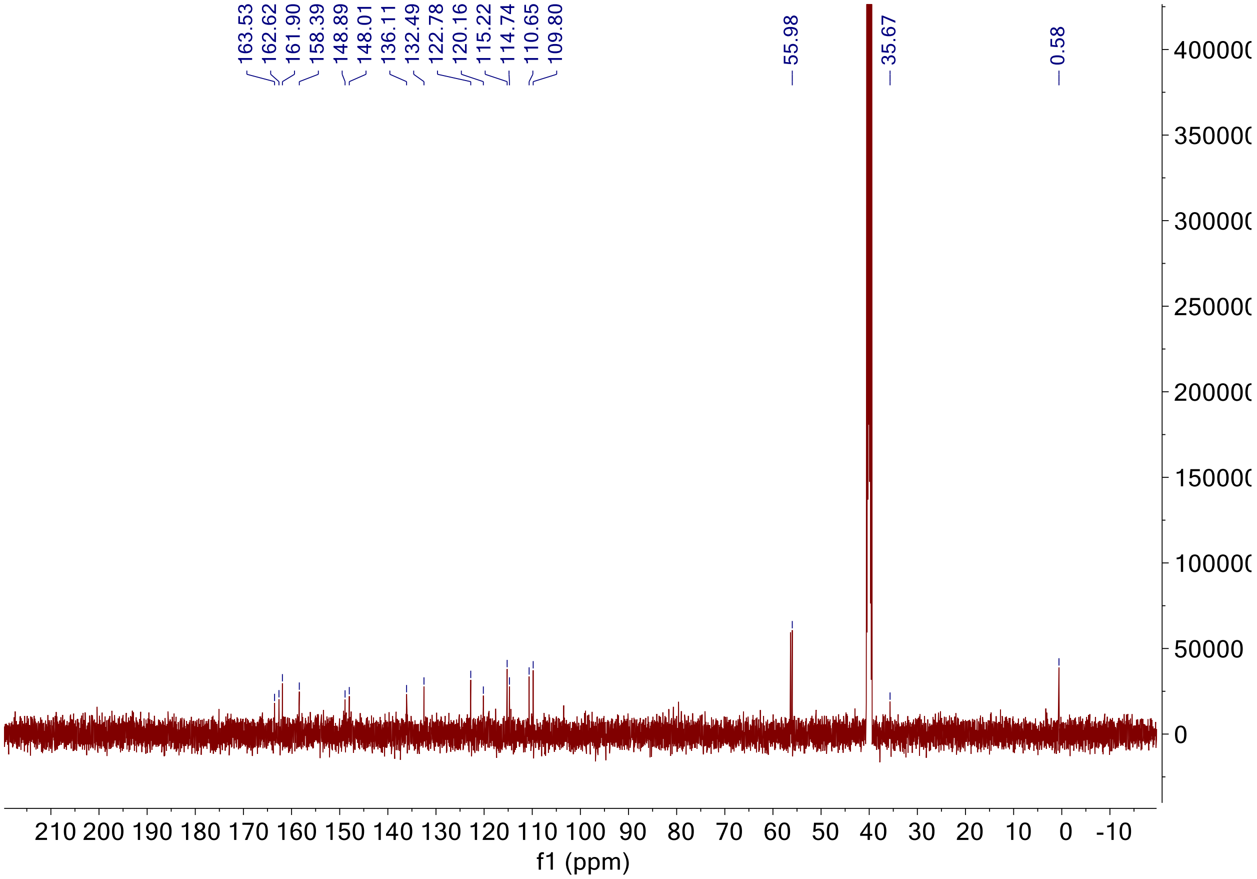

Signal DAD1 B, Sig=210,4 Ref=360,100 (bw-12_Mohan_walkuptest05092019_0331.D)

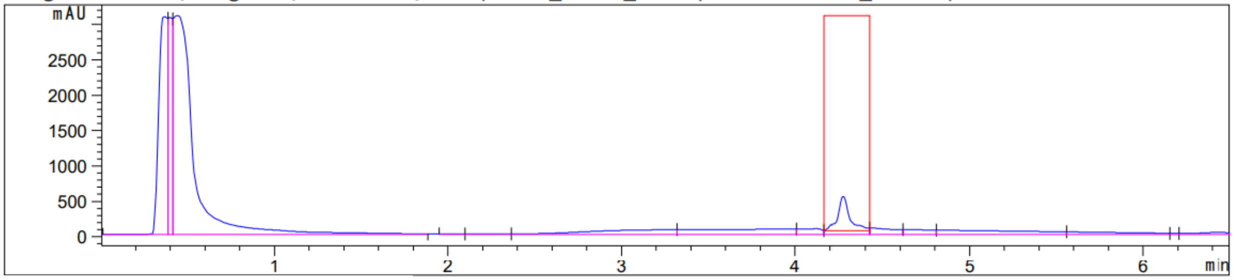

Peak :19 at 4.272 min Name : ?

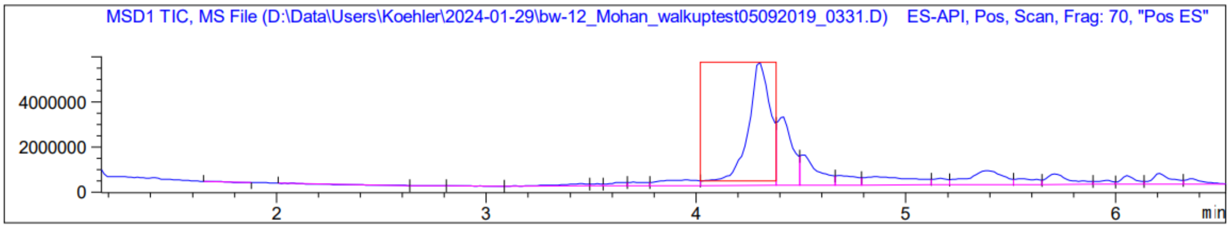
MSD1 TIC, MS File (D:\Data\Users\Koehler\2024-01-29\bw-12_Mohan_walkuptest05092019_0331.D)    ES-API, Pos, Scan, Frag: 70, "Pos ES"
4000000
2000000
0
2
3
4
5
6
min

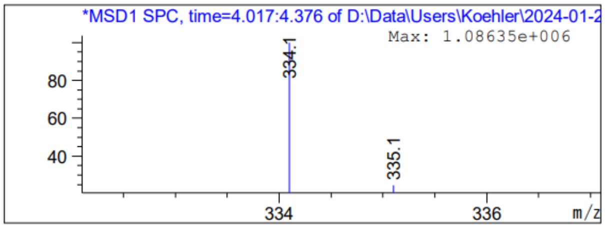
*MSD1 SPC, time=4.017:4.376 of D:\Data\Users\Koehler\2024-01-2
Max: 1.08635e+006
334.1
335.1
80
60
40
334
336
m/z

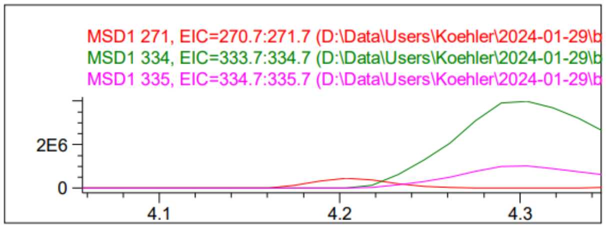
MSD1 271, EIC=270.7:271.7 (D:\Data\Users\Koehler\2024-01-29\b
MSD1 334, EIC=333.7:334.7 (D:\Data\Users\Koehler\2024-01-29\b
MSD1 335, EIC=334.7:335.7 (D:\Data\Users\Koehler\2024-01-29\b
2E6
0
4.1
4.2
4.3

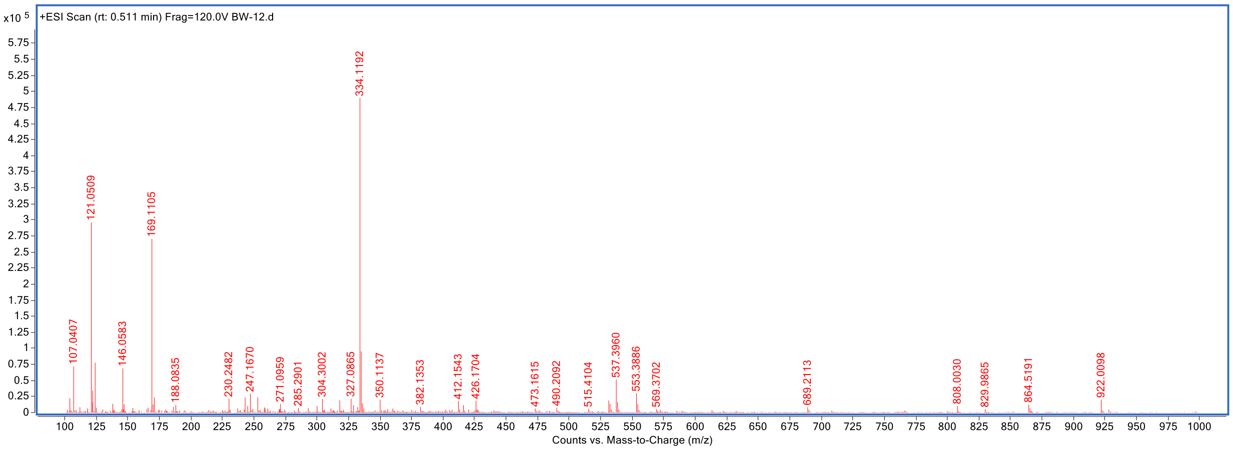
x10 5
+ESI Scan (rt: 0.511 min) Frag=120.0V BW-12.d
334.1192
121.0509
169.1105
107.0407
146.0583
188.0835
230.2482
247.1670
271.0959
285.2901
304.3002
327.0865
350.1137
382.1353
412.1543
426.1704
473.1615
490.2092
515.4104
537.3960
553.3886
569.3702
689.2113
808.0030
829.9865
864.5191
922.0098
Counts vs. Mass-to-Charge (m/z)

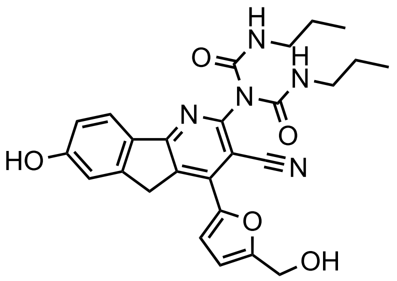

KI-TOX-A3-0189

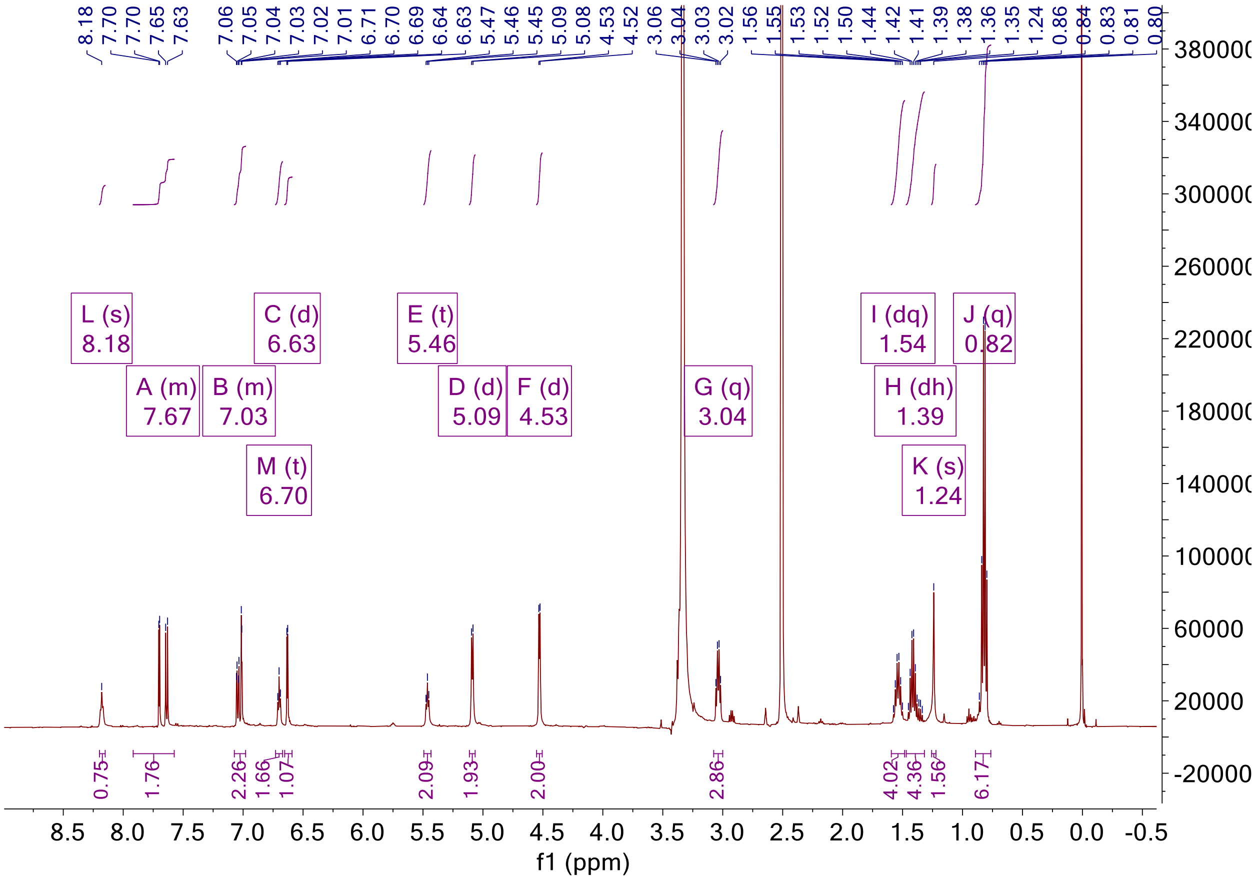

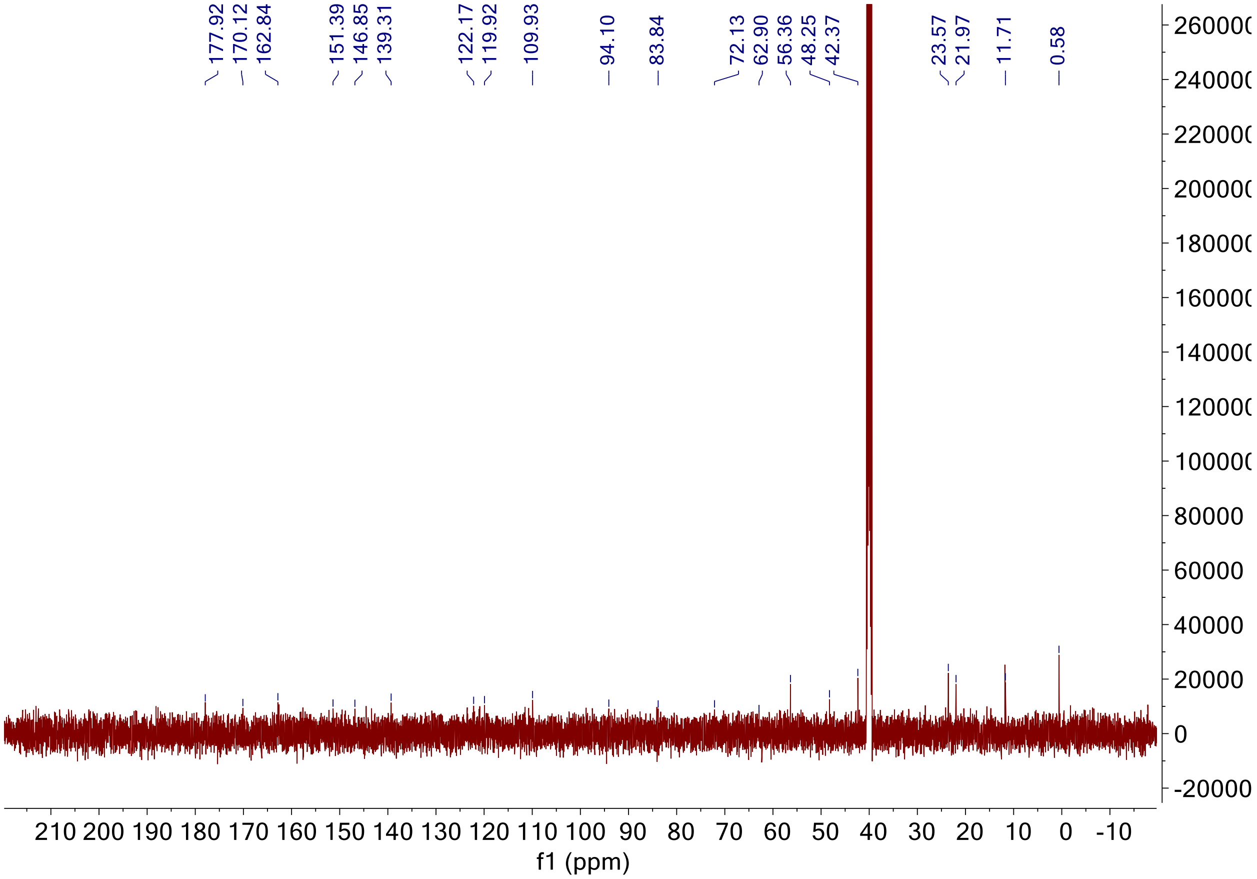

Signal DAD1 G, Sig=280,4 Ref=360,100 (bw-13L_Mohan_walkuptest05092019_0330.D)

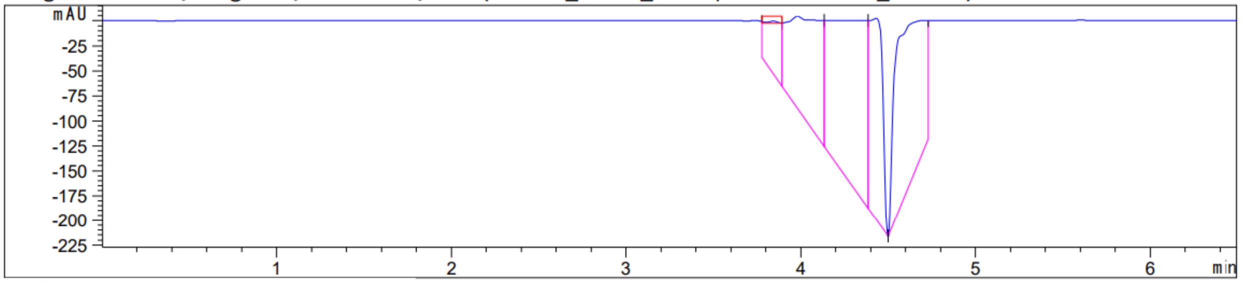

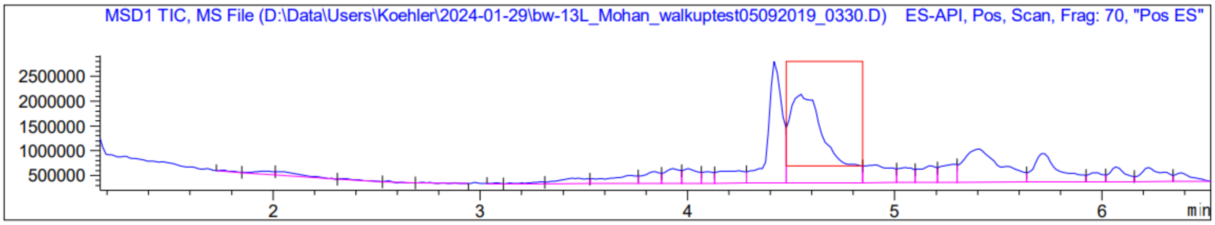
MSD1 TIC, MS File (D:\Data\Users\Koehler\2024-01-29\bw-13L_Mohan_walkuptest05092019_0330.D) ES-API, Pos, Scan, Frag: 70, "Pos ES"
min

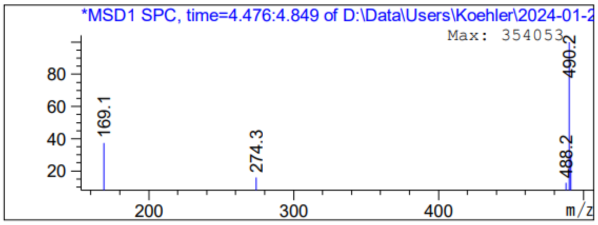
*MSD1 SPC, time=4.476:4.849 of D:\Data\Users\Koehler\2024-01-2
Max: 354053
169.1
274.3
488.2
490.2
m/z

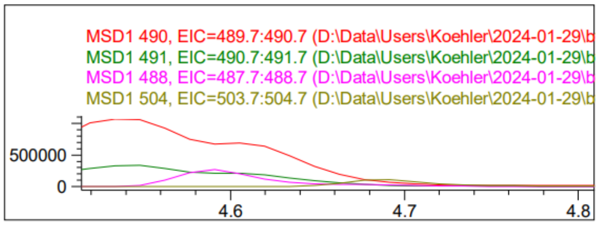
MSD1 490, EIC=489.7:490.7 (D:\Data\Users\Koehler\2024-01-29\b
MSD1 491, EIC=490.7:491.7 (D:\Data\Users\Koehler\2024-01-29\b
MSD1 488, EIC=487.7:488.7 (D:\Data\Users\Koehler\2024-01-29\b
MSD1 504, EIC=503.7:504.7 (D:\Data\Users\Koehler\2024-01-29\b

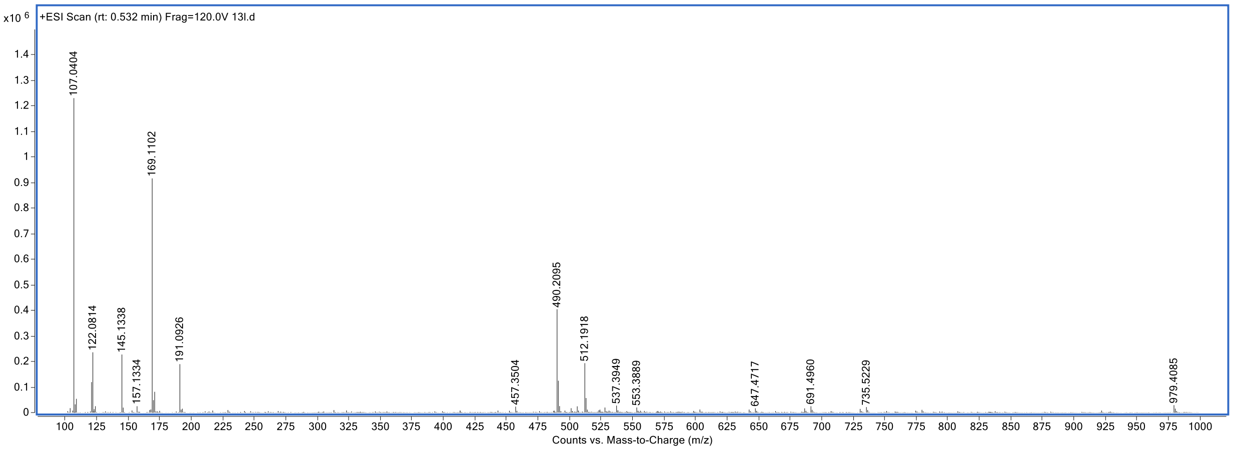
+ESI Scan (rt: 0.532 min) Frag=120.0V 13l.d
x10 6
107.0404
122.0814
145.1338
157.1334
169.1102
191.0926
457.3504
490.2095
512.1918
537.3949
553.3889
647.4717
691.4960
735.5229
979.4085
Counts vs. Mass-to-Charge (m/z)

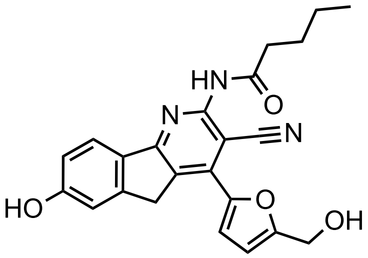

KI-TOX-A3-0190

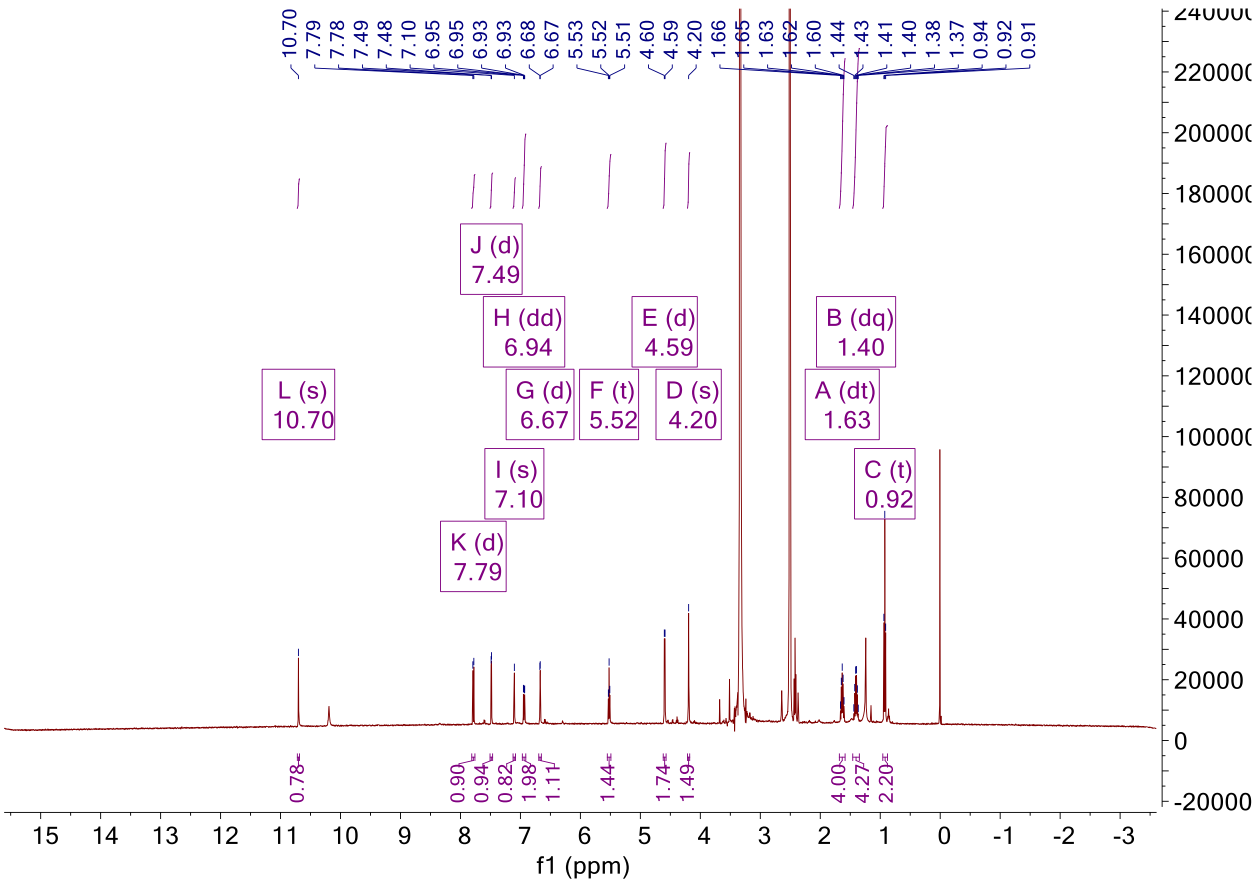

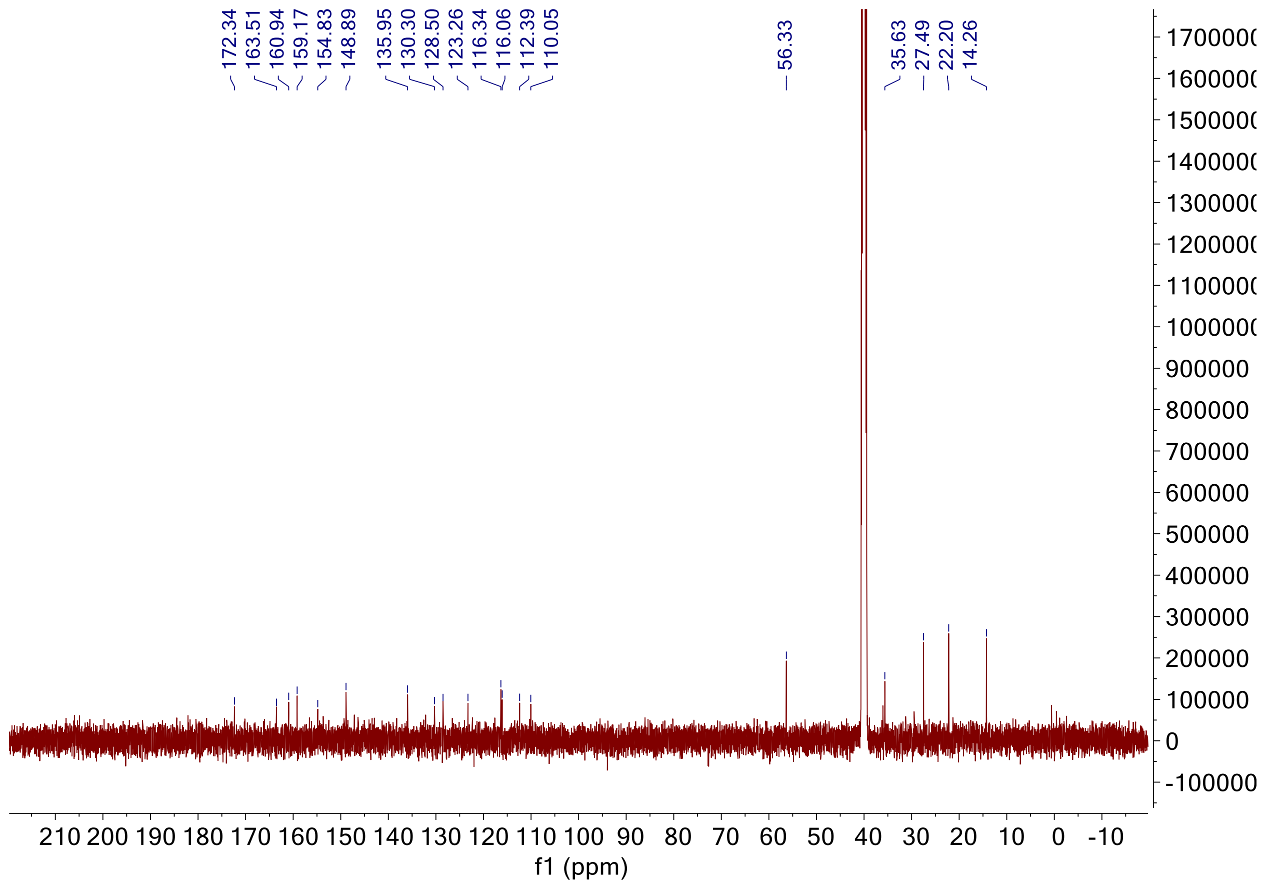

172.34
163.51
160.94
159.17
154.83
148.89
135.95
130.30
128.50
123.26
116.34
116.06
112.39
110.05
56.33
35.63
27.49
22.20
14.26
f1 (ppm)

Signal DAD1 G, Sig=280,4 Ref=360,100 (bw-14_Mohan_walkuptest05092019_0329.D)

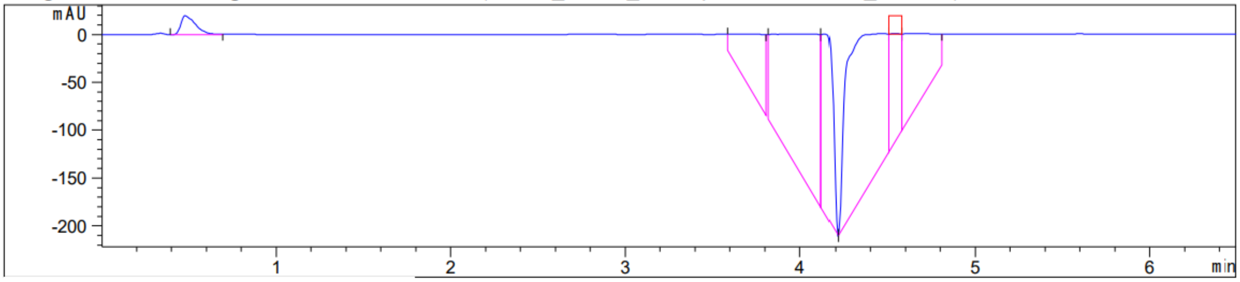

Peak :26 at 4.541 min  Name : ?

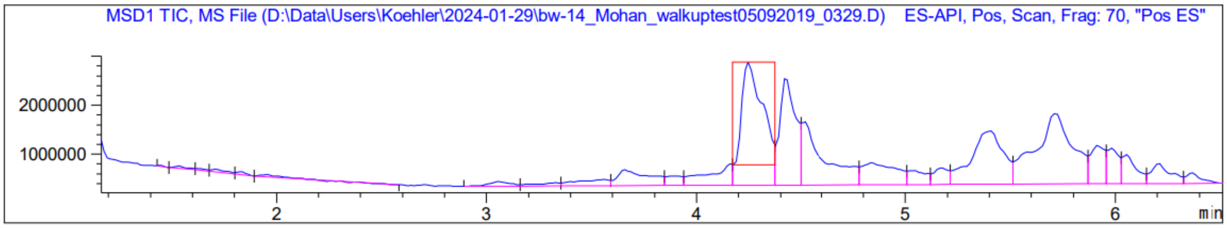

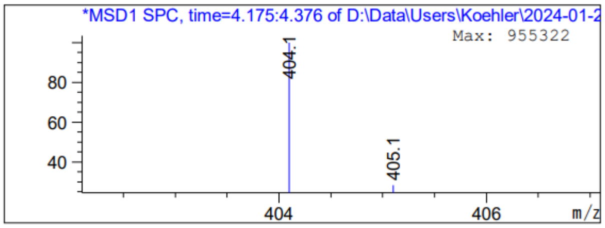

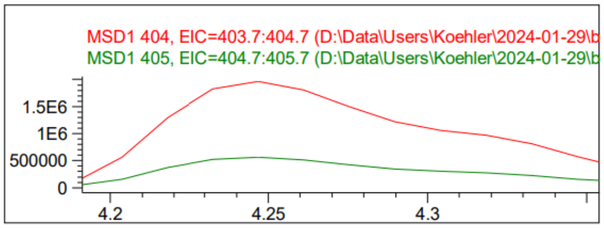

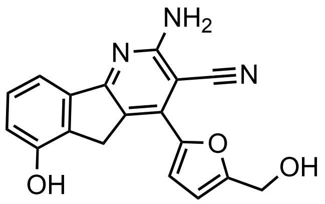

KI-TOX-A3-0179

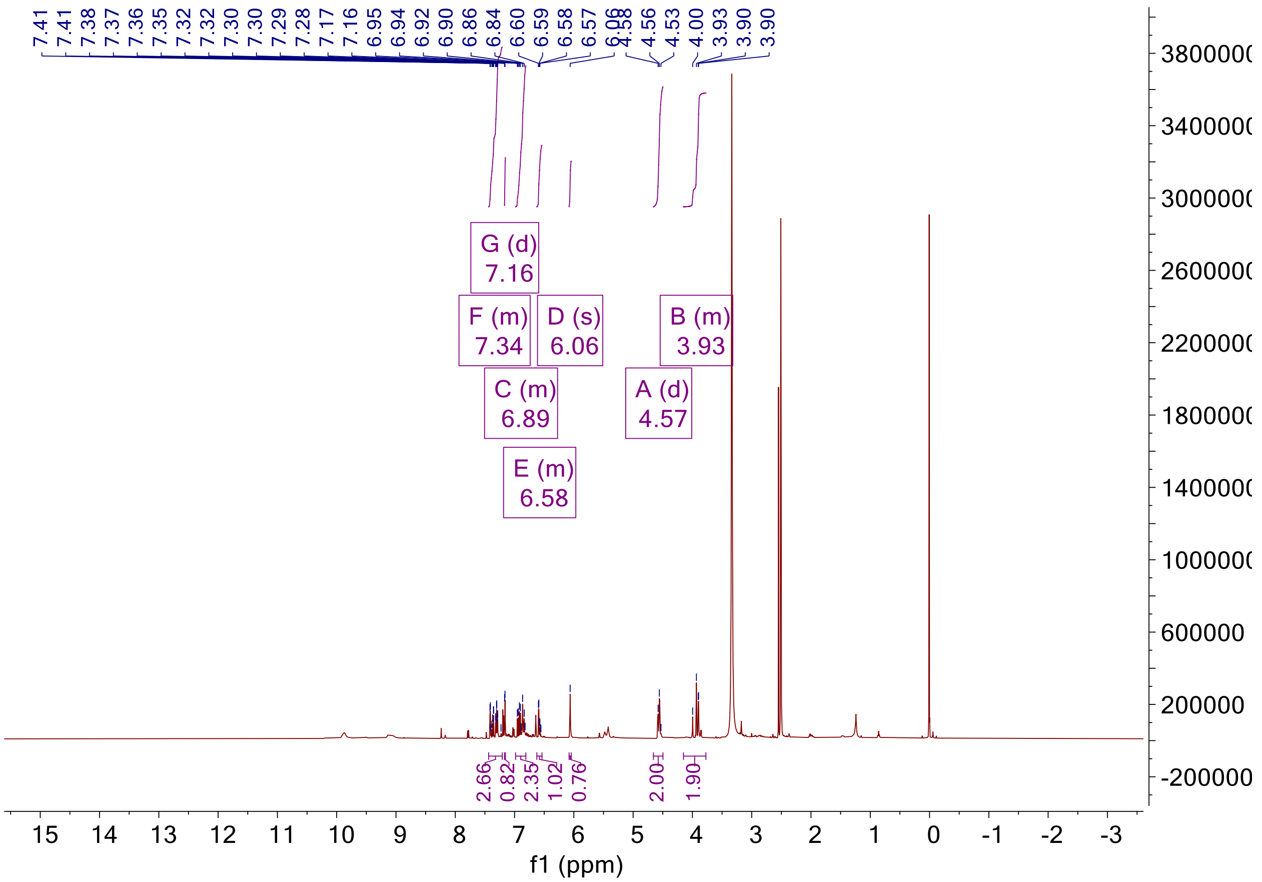

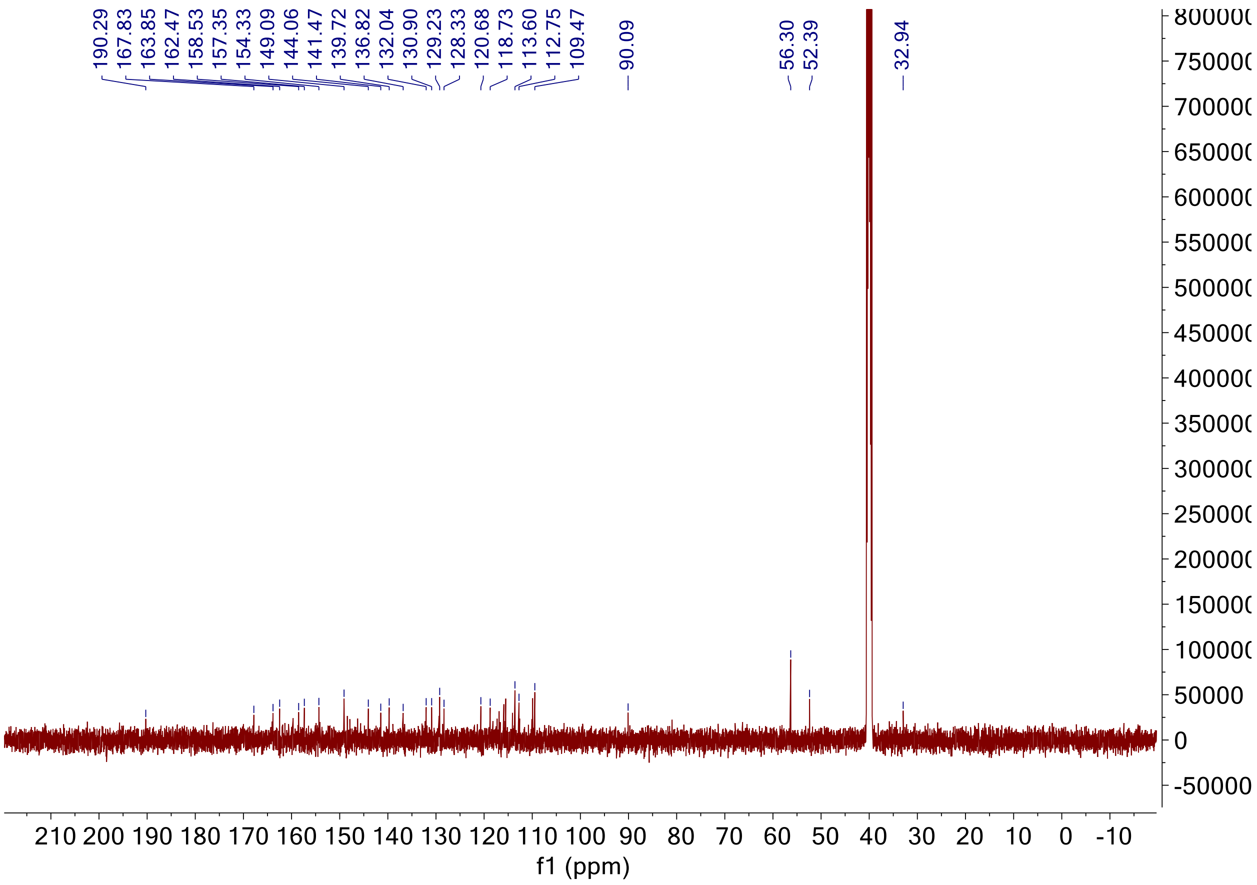

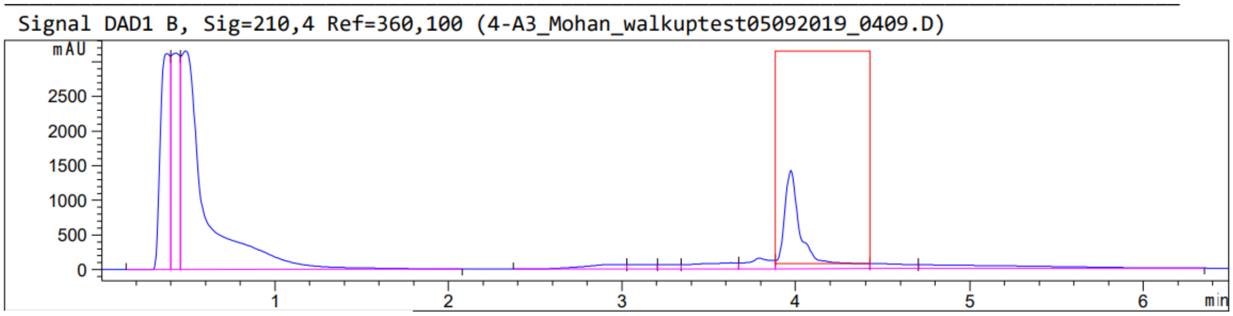

Peak :21 at 3.974 min Name : ?

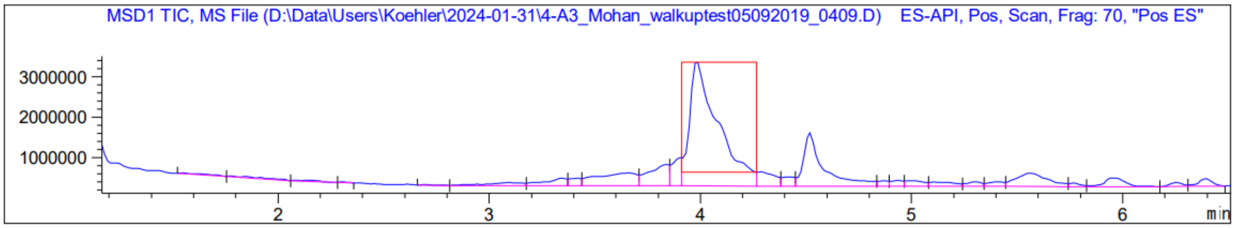
MSD1 TIC, MS File (D:\Data\Users\Koehler\2024-01-31\4-A3_Mohan_walkuptest05092019_0409.D) ES-API, Pos, Scan, Frag: 70, "Pos ES"
3000000
2000000
1000000
2
3
4
5
6
min

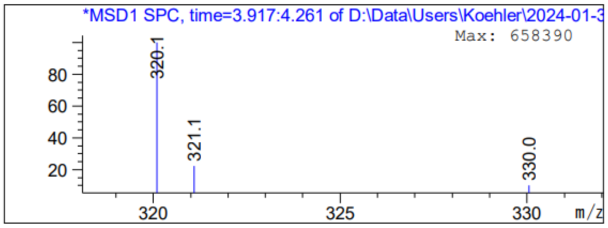
*MSD1 SPC, time=3.917:4.261 of D:\Data\Users\Koehler\2024-01-3
Max: 658390
320.1
321.1
330.0
80
60
40
20
320
325
330
m/z

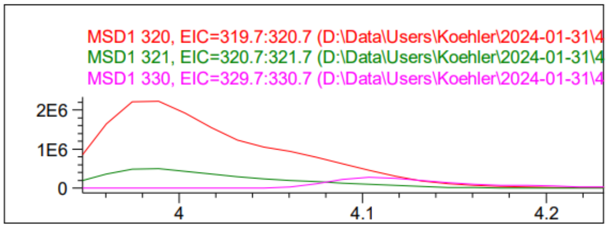
MSD1 320, EIC=319.7:320.7 (D:\Data\Users\Koehler\2024-01-31\4
MSD1 321, EIC=320.7:321.7 (D:\Data\Users\Koehler\2024-01-31\4
MSD1 330, EIC=329.7:330.7 (D:\Data\Users\Koehler\2024-01-31\4
2E6
1E6
0
4
4.1
4.2

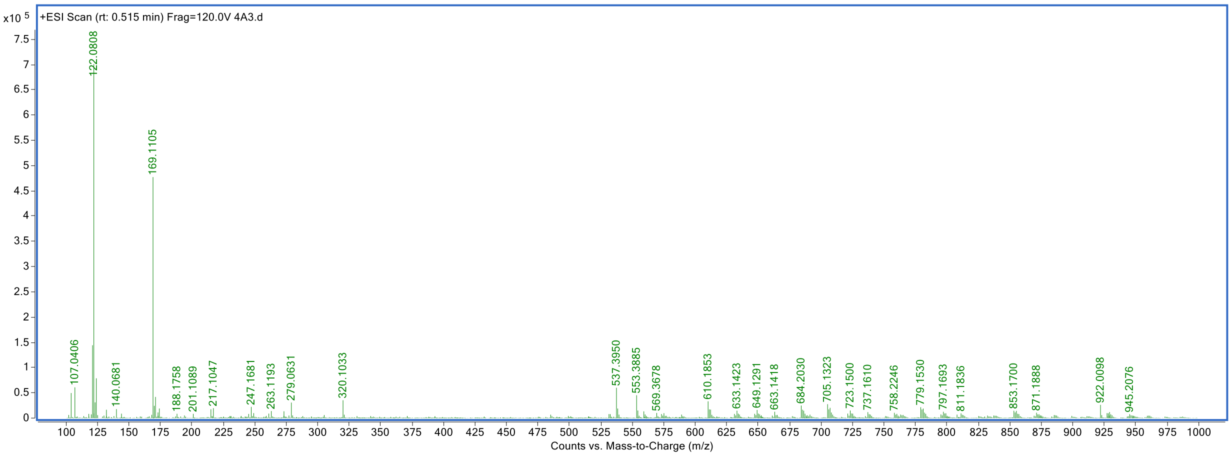
x10 5
+ESI Scan (rt: 0.515 min) Frag=120.0V 4A3.d
122.0808
169.1105
107.0406
140.0681
188.1758
201.1089
217.1047
247.1681
263.1193
279.0631
320.1033
537.3950
553.3885
569.3678
610.1853
633.1423
649.1291
663.1418
684.2030
705.1323
723.1500
737.1610
758.2246
779.1530
797.1693
811.1836
853.1700
871.1888
922.0098
945.2076
Counts vs. Mass-to-Charge (m/z)

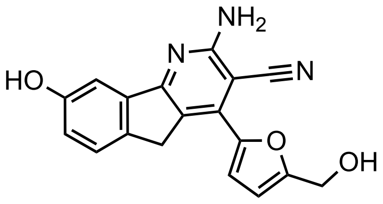

KI-TOX-A3-0178

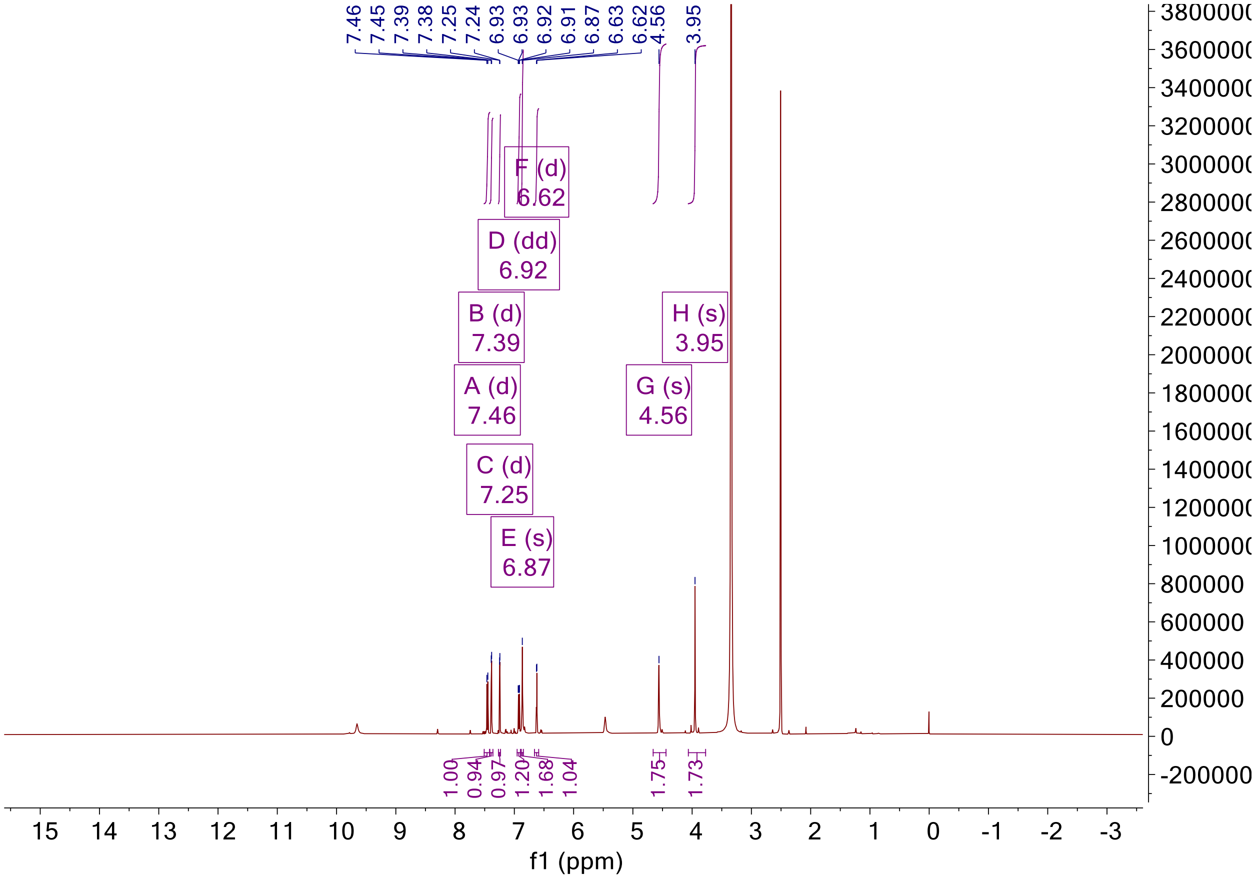

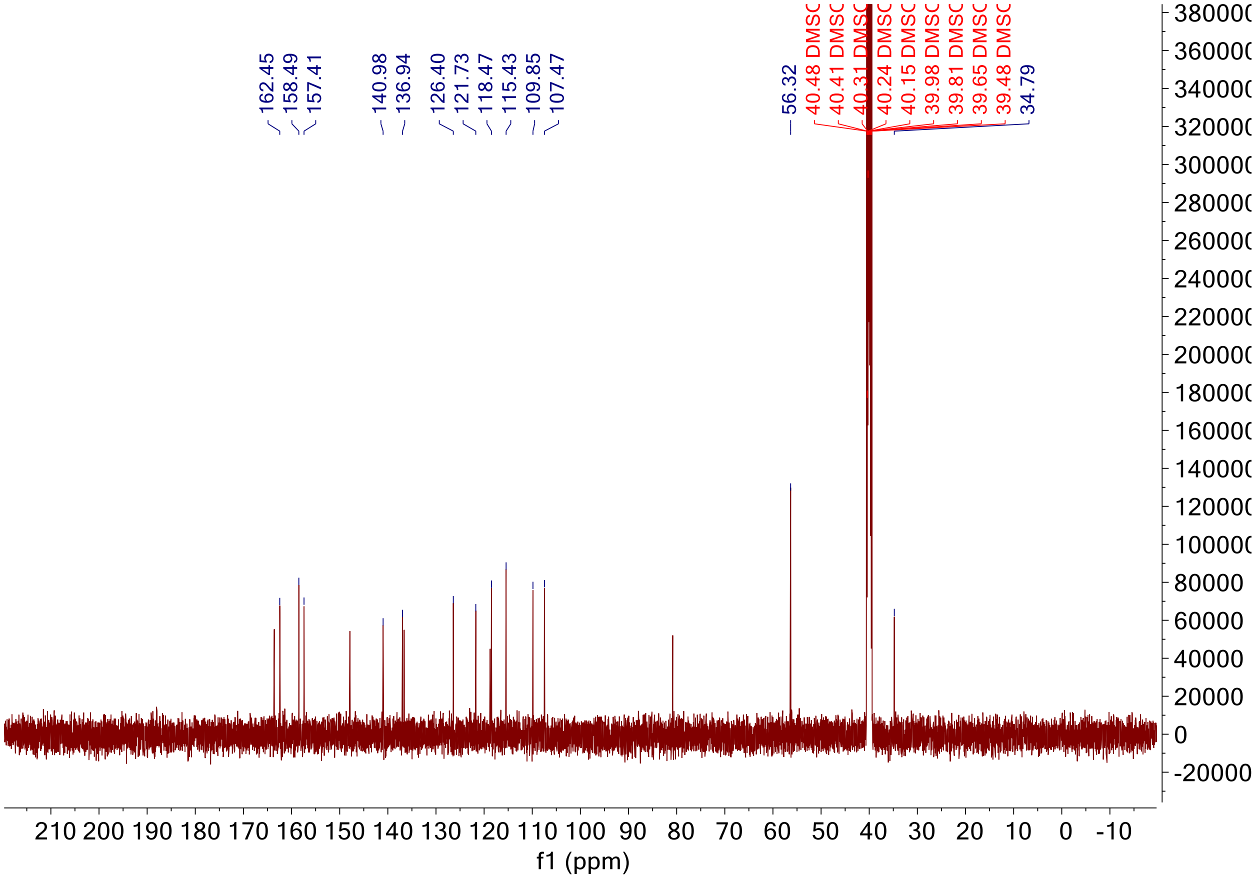

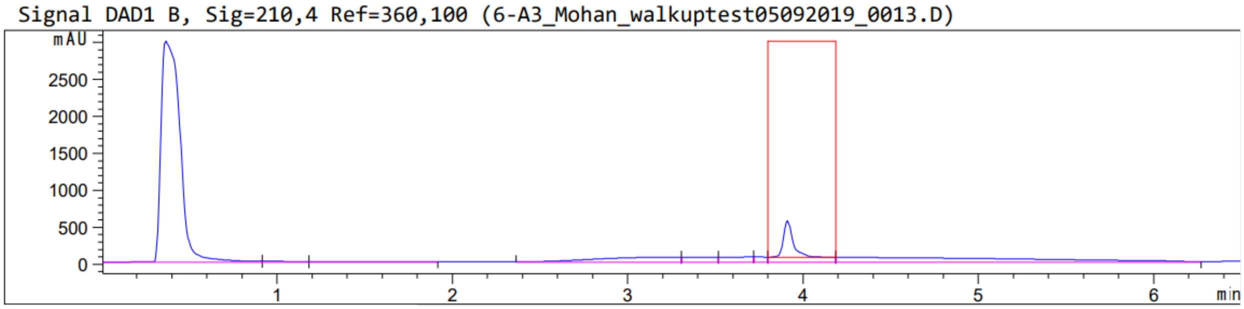

Peak :21 at 3.911 min Name : ?

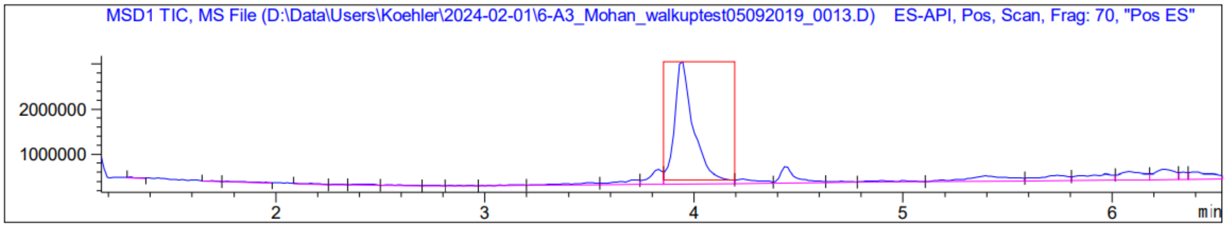
MSD1 TIC, MS File (D:\Data\Users\Koehler\2024-02-01\6-A3_Mohan_walkuptest05092019_0013.D) ES-API, Pos, Scan, Frag: 70, "Pos ES"
2000000
1000000
2
3
4
5
6
min

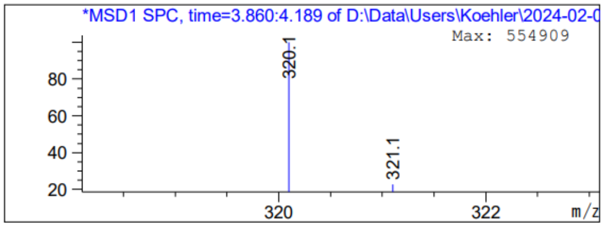
*MSD1 SPC, time=3.860:4.189 of D:\Data\Users\Koehler\2024-02-0
Max: 554909
320.1
321.1
80
60
40
20
320
322
m/z

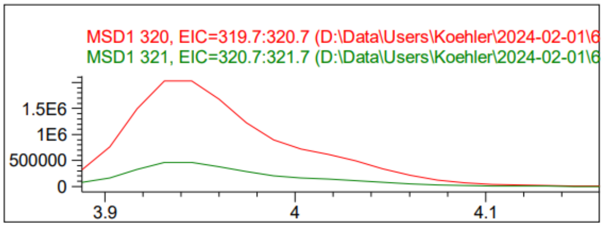
MSD1 320, EIC=319.7:320.7 (D:\Data\Users\Koehler\2024-02-01\6
MSD1 321, EIC=320.7:321.7 (D:\Data\Users\Koehler\2024-02-01\6
1.5E6
1E6
500000
0
3.9
4
4.1

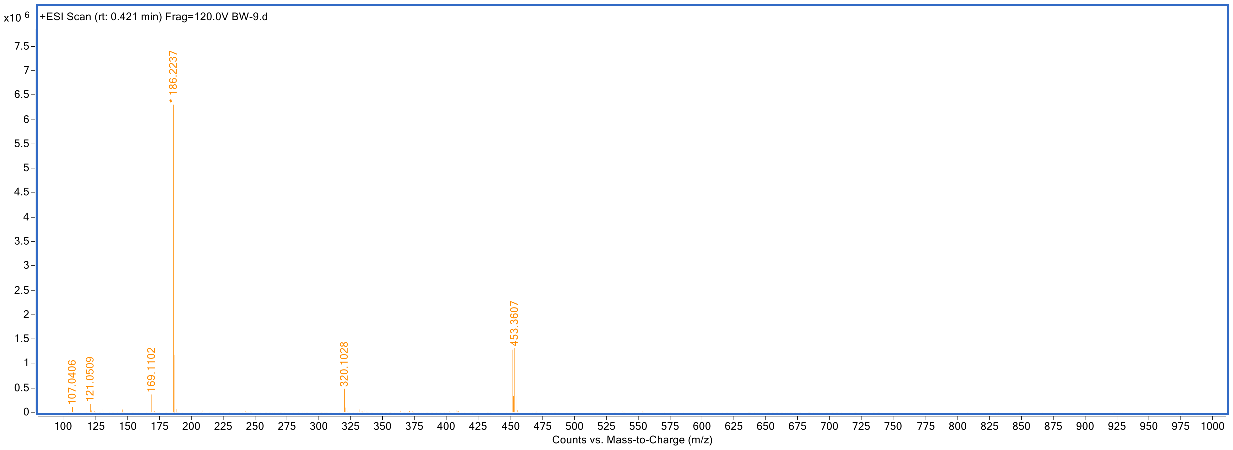
x10 6
+ESI Scan (rt: 0.421 min) Frag=120.0V BW-9.d
* 186.2237
107.0406
121.0509
169.1102
320.1028
453.3607
Counts vs. Mass-to-Charge (m/z)

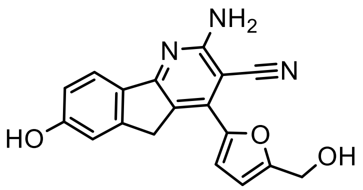
NH2
N
N
HO
O
OH

KI-TOX-A3

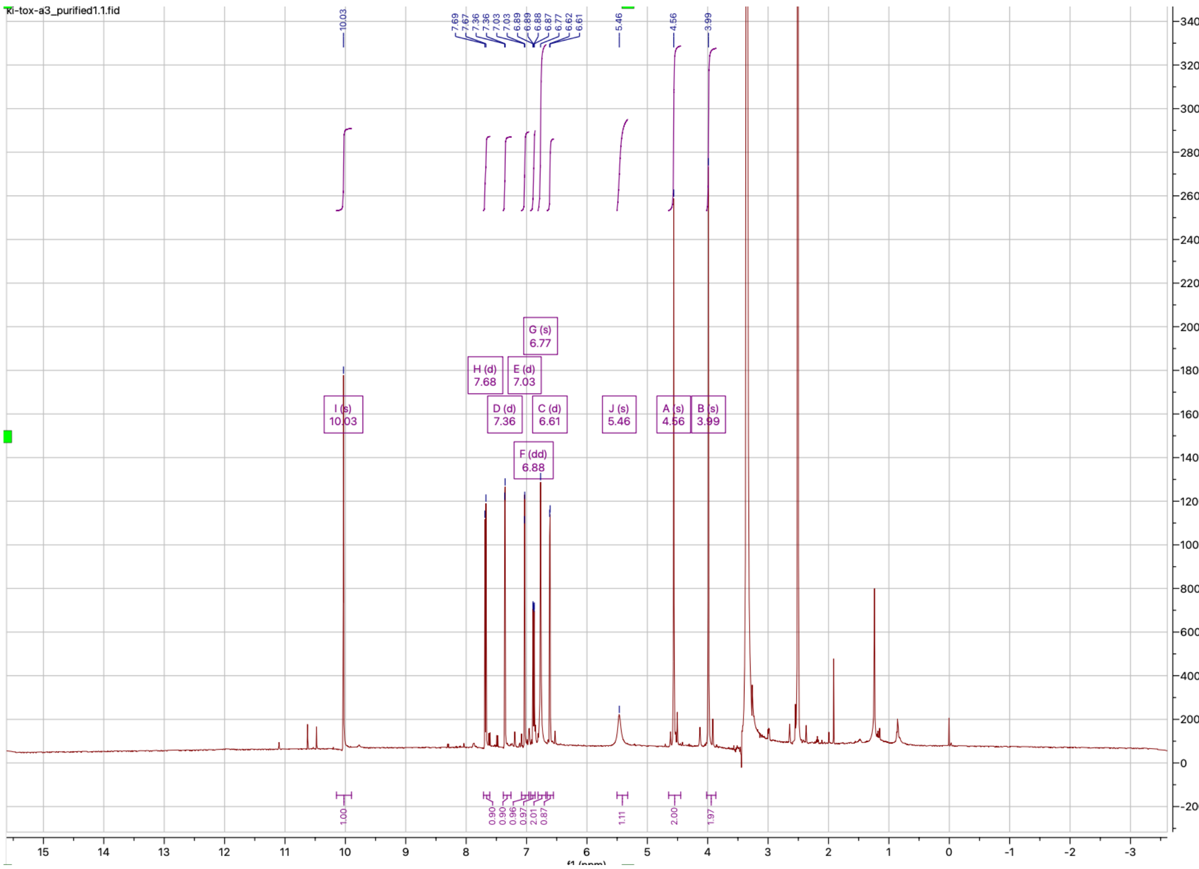
ki-tox-a3_purified1.1.fid
10.03
7.69
7.67
7.36
7.36
7.03
7.03
6.89
6.89
6.88
6.87
6.77
6.62
6.61
5.46
4.56
3.99
G (s)
6.77
H (d)
7.68
E (d)
7.03
I (s)
10.03
D (d)
7.36
C (d)
6.61
J (s)
5.46
A (s)
4.56
B (s)
3.99
F (dd)
6.88
1.00
0.90
0.90
0.96
0.97
2.01
0.87
1.11
2.00
1.97
340000
320000
300000
280000
260000
240000
220000
200000
180000
160000
140000
120000
100000
80000
60000
40000
20000
0
-20000
15
14
13
12
11
10
9
8
7
6
5
4
3
2
1
0
-1
-2
-3

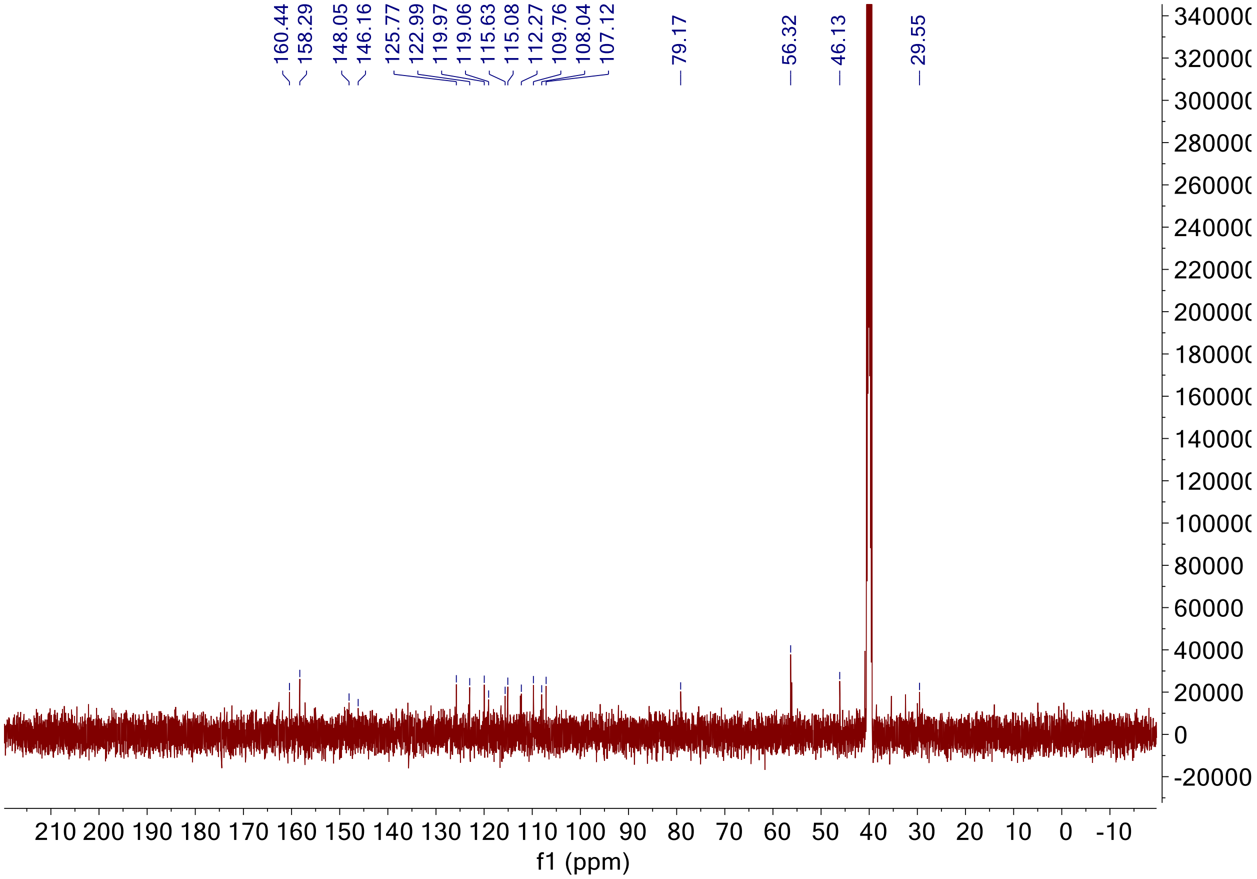
160.44
158.29
148.05
146.16
125.77
122.99
119.97
119.06
115.63
115.08
112.27
109.76
108.04
107.12
79.17
56.32
46.13
29.55
f1 (ppm)

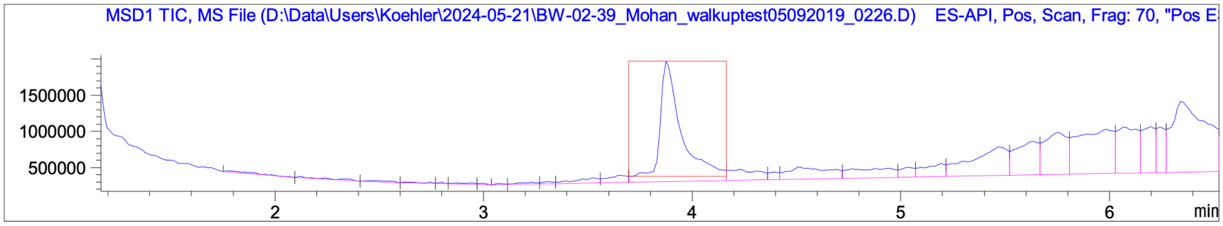
MSD1 TIC, MS File (D:\Data\Users\Koehler\2024-05-21\BW-02-39_Mohan_walkuptest05092019_0226.D) ES-API, Pos, Scan, Frag: 70, "Pos E
min

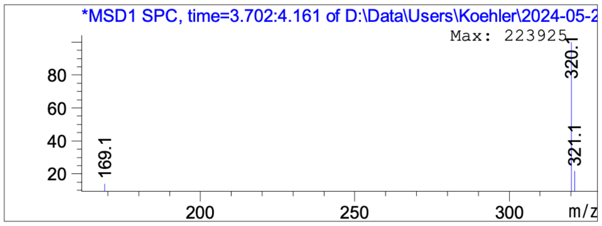
*MSD1 SPC, time=3.702:4.161 of D:\Data\Users\Koehler\2024-05-2
Max: 223925
320.1
321.1
169.1
m/z

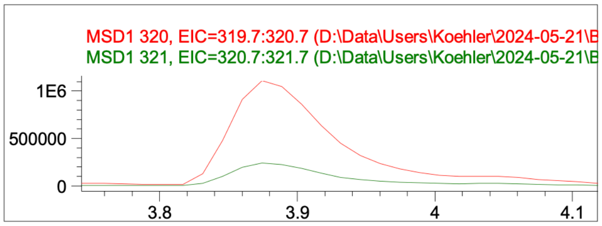
MSD1 320, EIC=319.7:320.7 (D:\Data\Users\Koehler\2024-05-21\E
MSD1 321, EIC=320.7:321.7 (D:\Data\Users\Koehler\2024-05-21\E

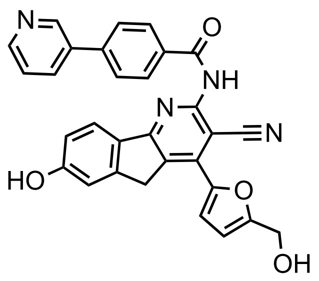

KI-TOX-A3-0245

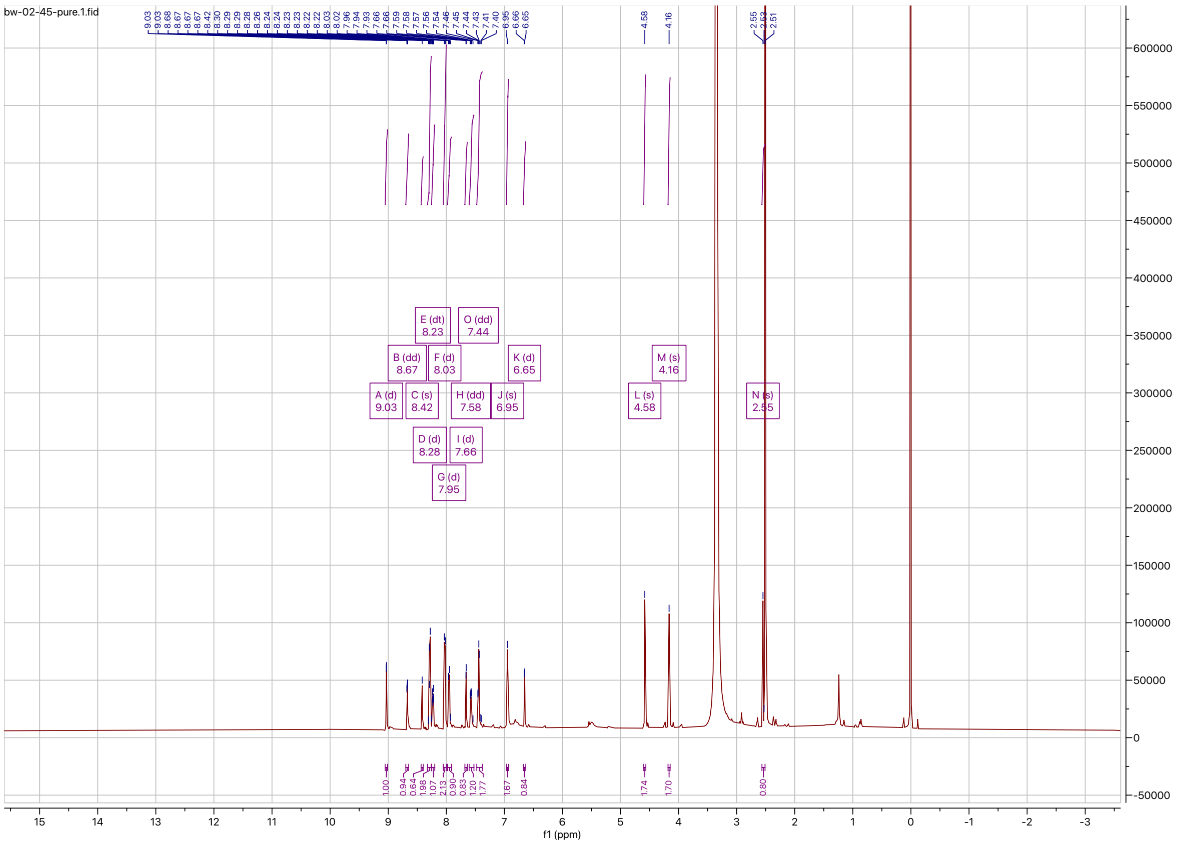

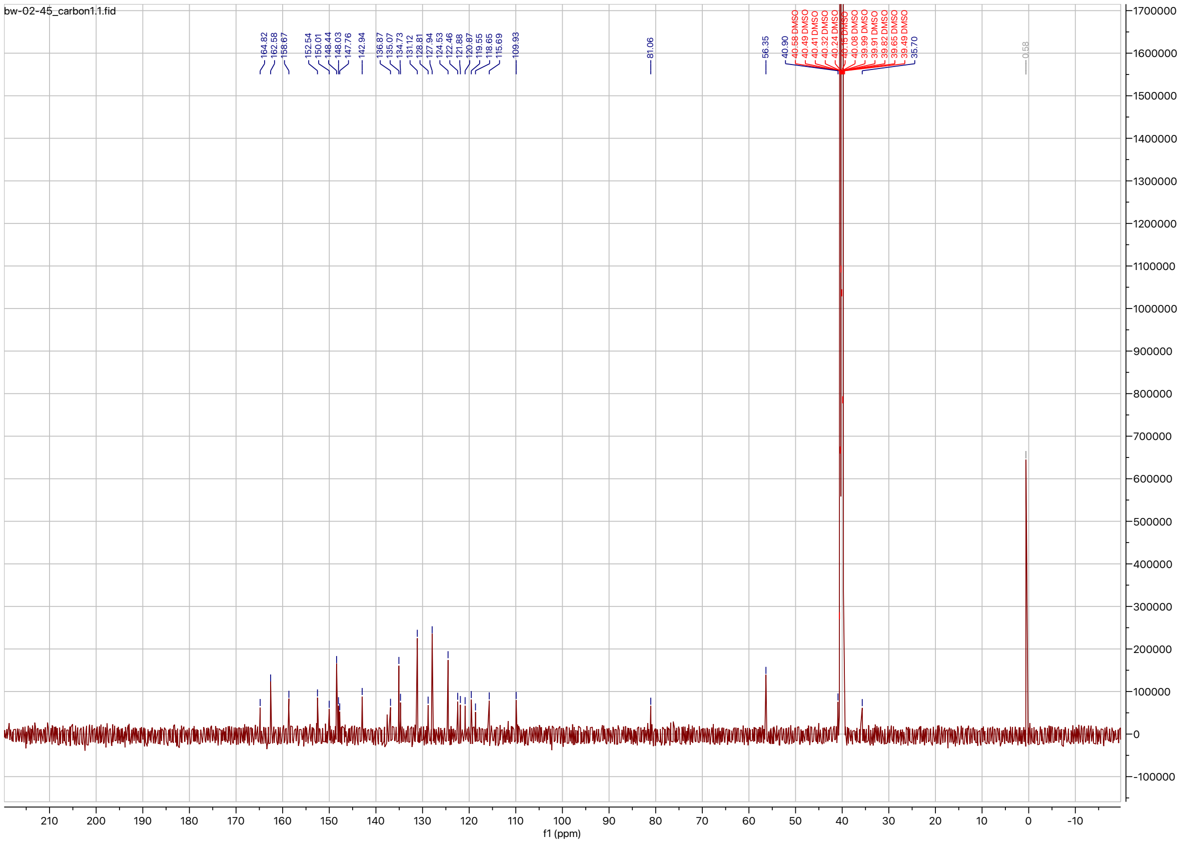

Signal DAD1 B, Sig=210,4 Ref=360,100 (BW-02-45_Mohan_walkuptest05092019_0258.D)

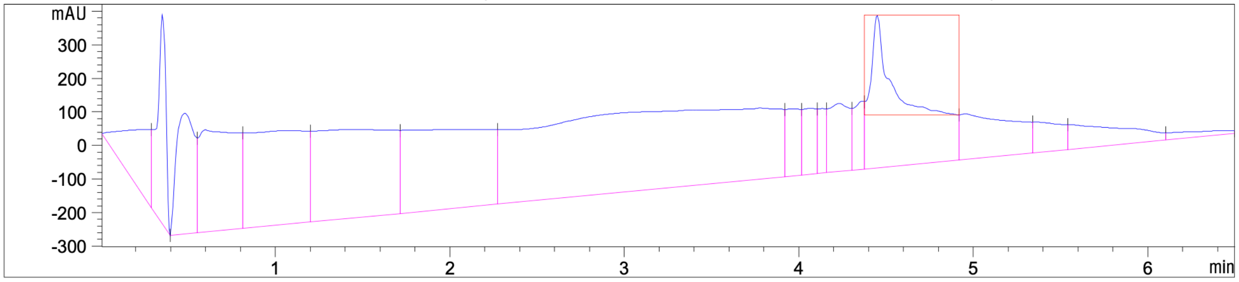

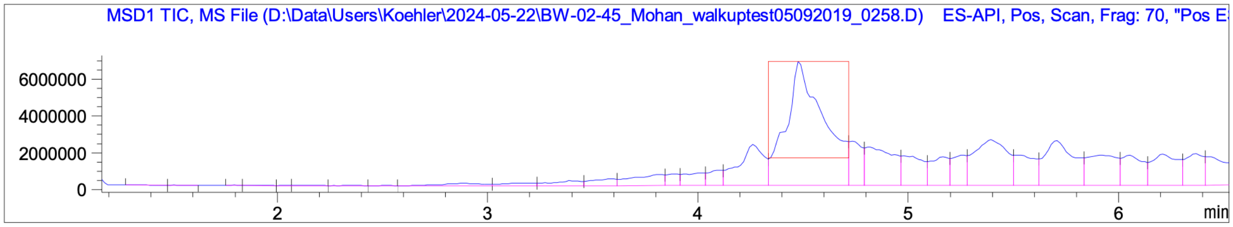

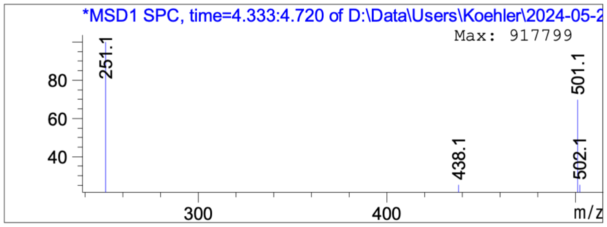

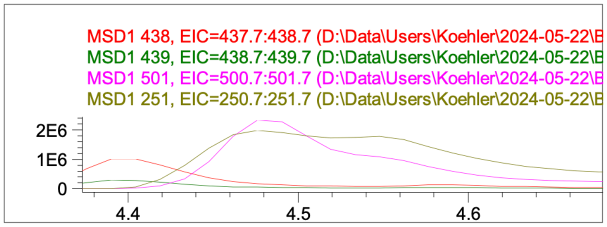

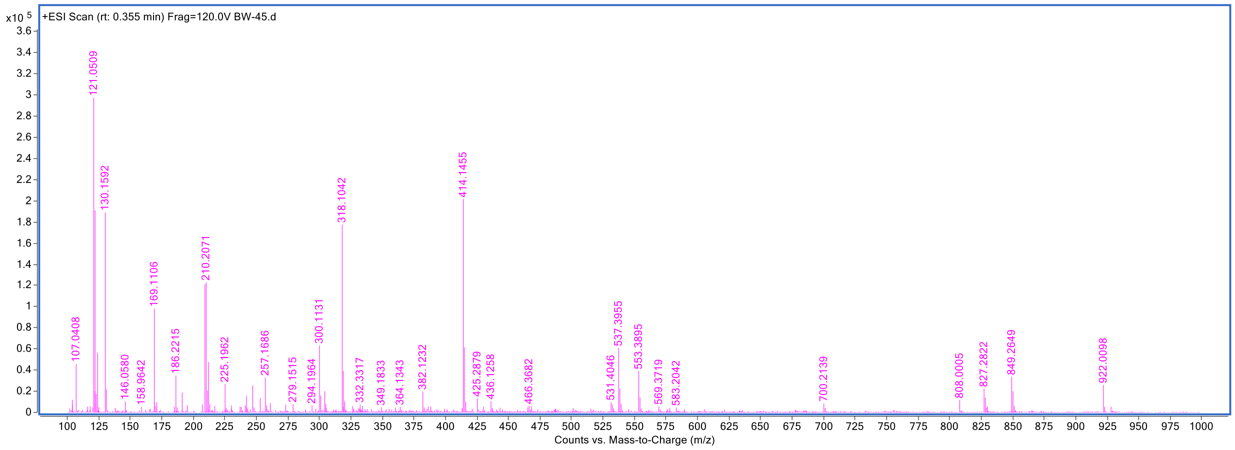

+ESI Scan (rt: 0.355 min) Frag=120.0V BW-45.d
x10 5
121.0509
130.1592
414.1455
318.1042
210.2071
169.1106
537.3955
300.1131
107.0408
186.2215
225.1962
257.1686
553.3895
849.2649
827.2822
922.0098
382.1232
146.0580
158.9642
279.1515
294.1964
332.3317
349.1833
364.1343
425.2879
436.1258
466.3682
531.4046
569.3719
583.2042
700.2139
808.0005
Counts vs. Mass-to-Charge (m/z)

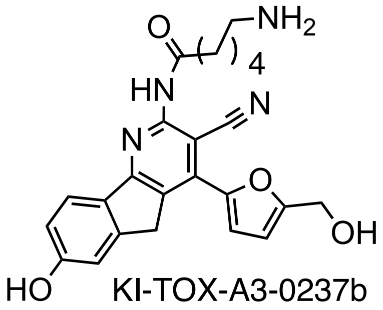

KI-TOX-A3-0237b

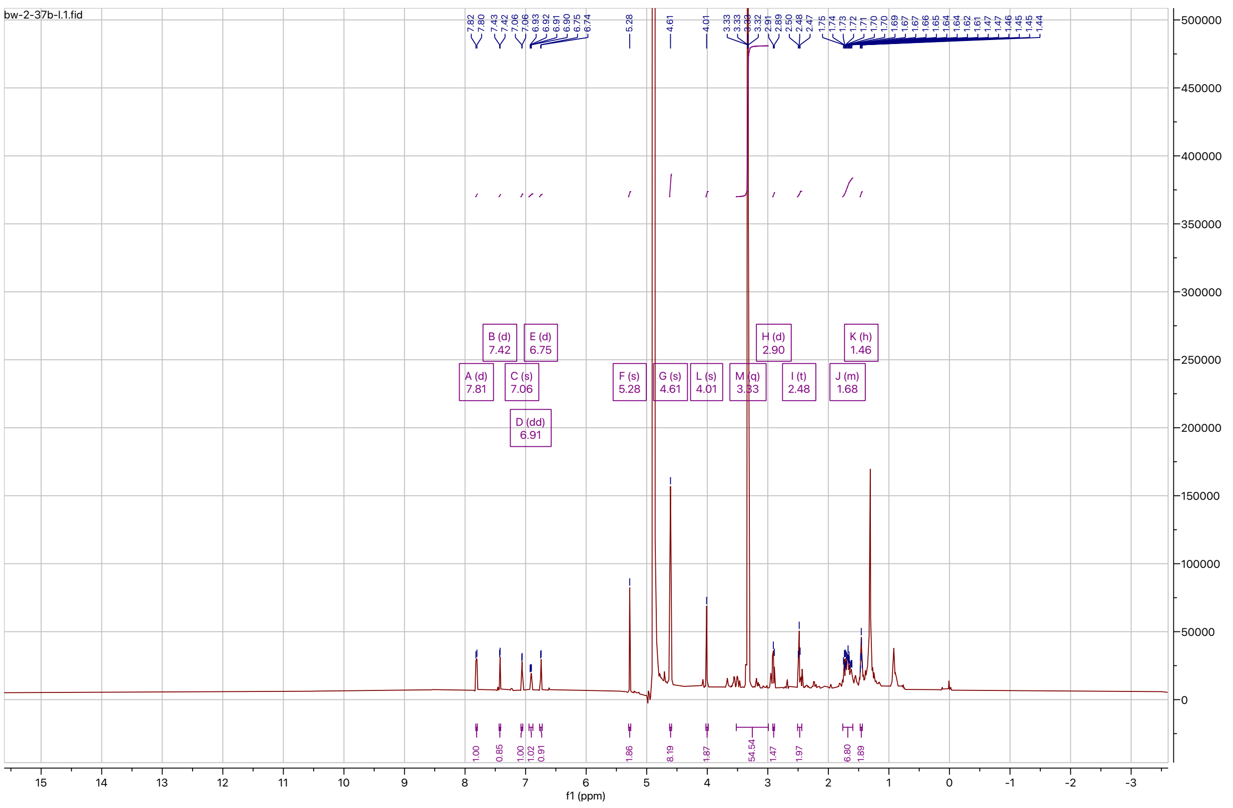

Signal DAD1 B, Sig=210,4 Ref=360,100 (BW-02-37B-14_Mohan_walkuptest05092019_0333.D)

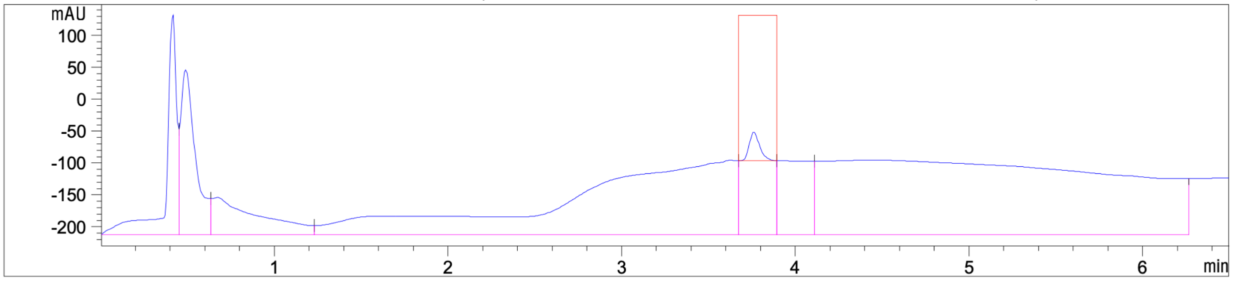

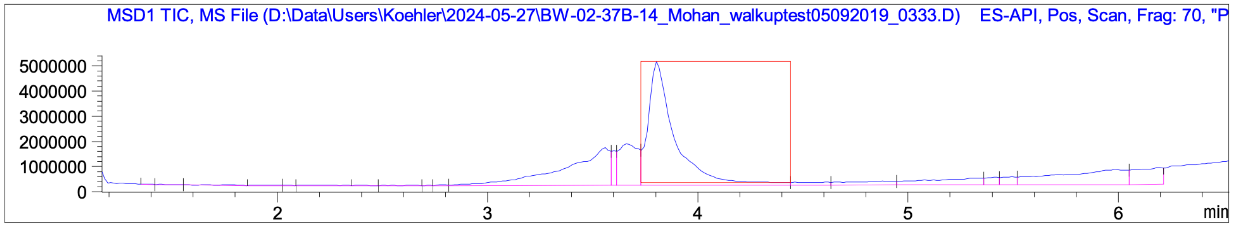
MSD1 TIC, MS File (D:\Data\Users\Koehler\2024-05-27\BW-02-37B-14_Mohan_walkuptest05092019_0333.D) ES-API, Pos, Scan, Frag: 70, "P
5000000
4000000
3000000
2000000
1000000
0
2
3
4
5
6
min

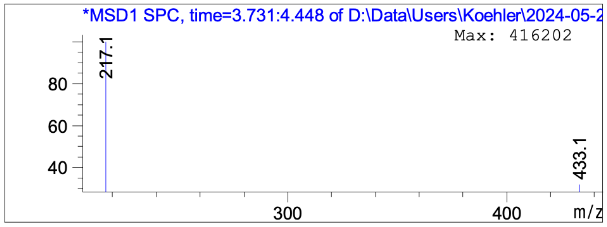
*MSD1 SPC, time=3.731:4.448 of D:\Data\Users\Koehler\2024-05-2
Max: 416202
217.1
433.1
80
60
40
300
400
m/z

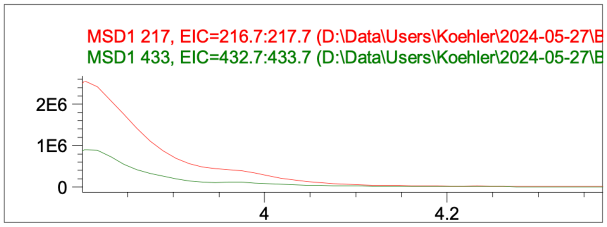
MSD1 217, EIC=216.7:217.7 (D:\Data\Users\Koehler\2024-05-27\B
MSD1 433, EIC=432.7:433.7 (D:\Data\Users\Koehler\2024-05-27\B
2E6
1E6
0
4
4.2

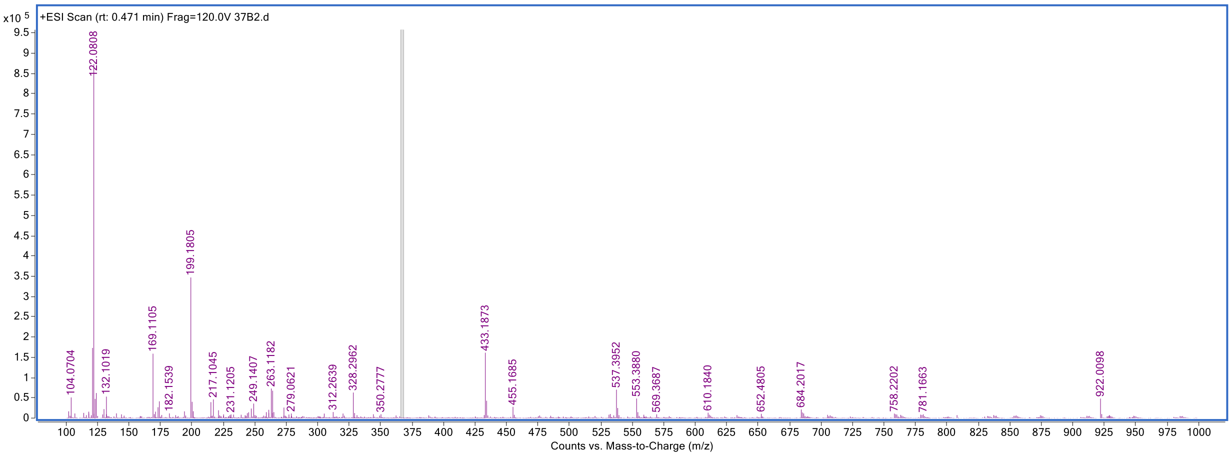
x10 5
+ESI Scan (rt: 0.471 min) Frag=120.0V 37B2.d
122.0808
199.1805
169.1105
433.1873
537.3952
553.3880
922.0098
104.0704
132.1019
182.1539
217.1045
231.1205
249.1407
263.1182
279.0621
312.2639
328.2962
350.2777
455.1685
569.3687
610.1840
652.4805
684.2017
758.2202
781.1663
Counts vs. Mass-to-Charge (m/z)
